# Bacterial surface properties influence the activity of the TAT-RasGAP_317-326_ antimicrobial peptide

**DOI:** 10.1101/2020.10.02.321802

**Authors:** Maria Georgieva, Tytti Heinonen, Alessandra Vitale, Simone Hargraves, Senka Causevic, Trestan Pillonel, Leo Eberl, Christian Widmann, Nicolas Jacquier

## Abstract

Antibiotic resistance is an increasing threat for public health, underscoring the need for new antibacterial agents. Antimicrobial peptides (AMPs) represent an alternative to classical antibiotics. TAT-RasGAP_317-326_ is a recently described AMP effective against a broad range of bacteria, but little is known about the conditions that may influence its activity. Using RNA-sequencing and screening of mutant libraries, we show that *Escherichia coli* and *Pseudomonas aeruginosa* respond to TAT-RasGAP_317-326_ by regulating metabolic and stress response pathways, possibly implicating two-component systems. Our results also indicate that bacterial surface properties, in particular integrity of the lipopolysaccharide layer, influence peptide binding and entry. Finally, we found differences between bacterial species with respect to their rate of resistance emergence against this peptide. Our findings provide the basis for future investigation on the mode of action of this peptide and its potential clinical use as an antibacterial agent.

## Introduction

The spread of antibiotic resistance in many bacterial species is severely limiting the benefits of antibiotics and a growing number of infections are becoming harder to treat (O’Neill, 2016). Therefore, there is a need for new antimicrobials that could be used in the treatment of bacterial infections. Antimicrobial peptides (AMPs), several of which are already in clinical trials with promising results, represent a large source of antibacterial agents (Kumar et al., 2018). They are attractive alternatives to classical antibiotics due to their broad-spectrum activity that allows the targeting of a wide variety of bacterial species (Di Somma et al., 2020). In addition, AMPs have a relatively simple structure that can be bioengineered to increase, for example, their stability under physiological conditions or their resistance to degradation by gastrointestinal tract enzymes after oral administration (Kong et al., 2020).

AMPs were first described as naturally occurring peptides produced by many different organisms. Thousands have been identified (Wang et al., 2016, Kumar et al., 2018). In bacteria, AMP-producing strains have an advantage over other strains or species during competitive colonization of ecological niches (Hassan et al., 2012). In multicellular organisms, AMPs such as the human cathelicidin LL-37 and the bovine bactenecin, are part of the innate immune system involved in the destruction of various microorganisms (Gennaro et al., 1989, Xhindoli et al., 2016).

Despite their diversity, AMPs share a number of common features: they are short peptides rich in cationic and hydrophobic amino acids and display an overall positive charge. To exert their biological activity, positively charged AMPs first interact with the negatively charged bacterial surface through electrostatic interactions (Brogden, 2005). This initial interaction with the bacterial surface is followed, for a majority of AMPs described up to date, by permeabilization and disruption of the membrane bilayer resulting in bacterial death. For example, this mechanism of killing has been demonstrated for melittin isolated from bee venom (Hong et al., 2019), human cathelicidin LL-37 (Mendez-Samperio, 2010), and polymyxin B derived from the Gram-positive bacterium *Bacillus polymyxa* (Srinivas and Rivard, 2017, Kumar et al., 2018).

TAT-RasGAP_317-326_ is a recently identified antimicrobial peptide that kills both Gram-positive and Gram-negative bacteria and has antibiofilm activity *in vitro* (Heulot et al., 2017, Heinonen et al., 2021). This peptide is composed of a cell permeable moiety, the TAT HIV 48-57 sequence, and a 10 amino acid sequence derived from the Src homology 3 domain of p120 RasGAP. TAT-RasGAP_317-326_ was initially identified as an anticancer compound that sensitizes cancer cells to genotoxins (Michod et al., 2004, Michod et al., 2009) and to radiotherapy (Tsoutsou et al., 2017). This peptide also inhibits cell migration and invasion (Barras et al., 2014c) and possesses anti-metastatic activity *in vivo* (Barras et al., 2014b). It can also directly lyse a subset of cancer cells by targeting plasma membrane inner leaflet-enriched phospholipids (Serulla et al., 2020) in a manner that does not involve known programmed cell death pathways (Annibaldi et al., 2014, Heulot et al., 2016). We have previously shown that the tryptophan residue at position 317 of the TAT-RasGAP_317-326_ peptide is essential for its activity against both eukaryotic and bacterial cells (Barras et al., 2014a, Heulot et al., 2017). Furthermore, we have reported that, despite its potent *in vitro* antimicrobial activity, TAT-RasGAP_317-326_ showed limited protection in a mouse model of *Escherichia coli* (*E. coli*)-induced peritonitis (Heulot et al., 2017). Physiological factors may have contributed to the poor biodistribution and rapid clearance of TAT-RasGAP_317-326_ and, subsequently, to its low efficacy in this setting (Michod et al., 2009).

In this study, we assessed TAT-RasGAP_317-326_ activity under various experimental settings to better characterize the peptide activity as well as the bacterial response to peptide exposure. Our findings provide important initial insights into the activity of the TAT-RasGAP_317-326_ peptide that will pave the way for further investigation on its antimicrobial properties.

## Results

### Divalent cations reduce TAT-RasGAP_317-326_ surface binding and entry into bacteria

*P. aeruginosa* grown under low Mg^2+^ conditions is resistant to EDTA, gentamicin and polymyxin B via a mechanism that involves outer membrane modifications (Macfarlane et al., 1999, McPhee et al., 2003, Olaitan et al., 2014). To assess whether the antimicrobial activity of TAT-RasGAP_317-326_ is also affected by Mg^2+^ levels, we assessed how Mg^2+^ modulated its minimal inhibitory concentration (MIC) and concentration inhibiting growth by 50% (IC_50_) in three laboratory strains, *E. coli* MG1655, *E. coli* ATCC 25922, and *P. aeruginosa* PA14. The results of these experiments are summarized in Table 1 and shown in detail in Supplementary Figures 1–3. The MIC of TAT-RasGAP_317-326_ in standard Luria-Bertani (LB) medium for *E. coli* and *P. aeruginosa* was determined to be 8 μM and 32 μM, respectively. There was a small difference in peptide MIC levels between LB and BM2 medium supplemented with 2 mM MgSO_4_ (BM2 Mg^high^) for both *E. coli* and *P. aeruginosa*. However, these two bacterial species displayed an 8-fold decrease in MIC of TAT-RasGAP_317-326_ in BM2 containing 20 μM MgSO_4_ (BM2 Mg^low^) relative to 2 mM MgSO_4_ (BM2 Mg^high^). Low magnesium in BM2 medium resulted in a 4-fold increase of MIC of polymyxin B in *P. aeruginosa* but had no impact in *E. coli*. Our results agree with earlier data that low Mg^2+^ increases *P. aeruginosa* resistance to polymyxin B (Macfarlane et al., 1999, McPhee et al., 2003). Altogether these findings show that low Mg^2+^ in culture medium renders bacterial cells more susceptible to TAT-RasGAP_317-326_ but not to polymyxin B, suggesting that TAT-RasGAP_317-326_ and polymyxin B attack bacterial cells through different mechanisms.

**Table 1:**
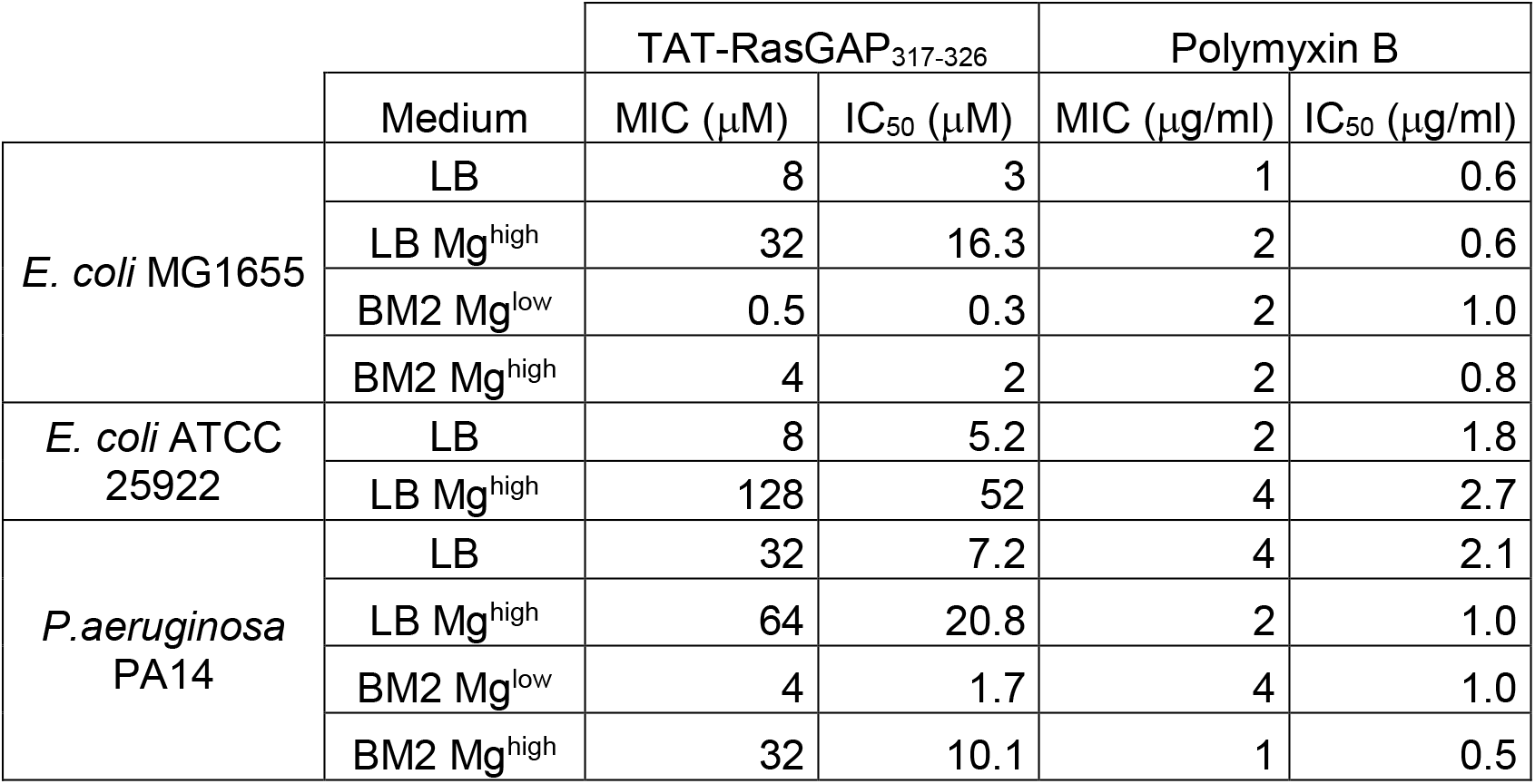
*E. coli* MG1655, ATCC25922 and *P. aeruginosa* PA14 sensitivity to TAT-RasGAP_317-326_ varies depending on the growth medium used. The indicated strains were grown overnight in LB, LB with 2 mM MgSO_4_ (LB Mg^high^), BM2 with 20 μM MgSO_4_ (BM2 Mg^low^) or BM2 with 2 mM MgSO_4_ (BM2 Mg^high^). Culture was then diluted to OD_600_ = 0.1 and grown for 1 hour. Bacterial suspension was further diluted 20 times for *E. coli* and 10 times for *P. aeruginosa* and 10 μl was added per well of a 96-well plate containing serial dilutions of TAT-RasGAP_317-326_.or polymyxin B. OD_590_ was measured after 16 hours of incubation. MIC is defined as the lowest concentration of TAT-RasGAP_317-326_ that completely inhibits bacterial proliferation. IC_50_ is defined as the concentration required to inhibit 50% of growth and was calculated using GraphPad Prism 8. The detailed growth curves are presented in Supplementary Fig. 1 to 3.

Because BM2 is a defined bacteriological medium, we also investigated whether Mg^2+^ affects the sensitivity to TAT-RasGAP_317-326_ in complex medium such as LB. We compared peptide MIC in LB and LB supplemented with 2 mM MgSO_4_ (LB Mg^high^) and found that high Mg^2+^ increased peptide MIC in both *E. coli* and *P. aeruginosa* (Table 1), consistent with the data obtained with these two bacterial strains in BM2 medium. Moreover, the ability of TAT-RasGAP_317-326_ to hamper *E. coli* growth rate was clearly inhibited by 2 mM MgSO_4_ (Fig. 1, panels A and B). Therefore, high Mg^2+^ levels decreased bacterial sensitivity to TAT-RasGAP_317-326_ and this result was independent of the medium used. We also determined that the MIC of polymyxin B for *P. aeruginosa* decreased in the presence of high Mg^2+^ in LB (Table 1), which is in accordance with our data for *P. aeruginosa* in BM2 medium.

**Figure 1.**
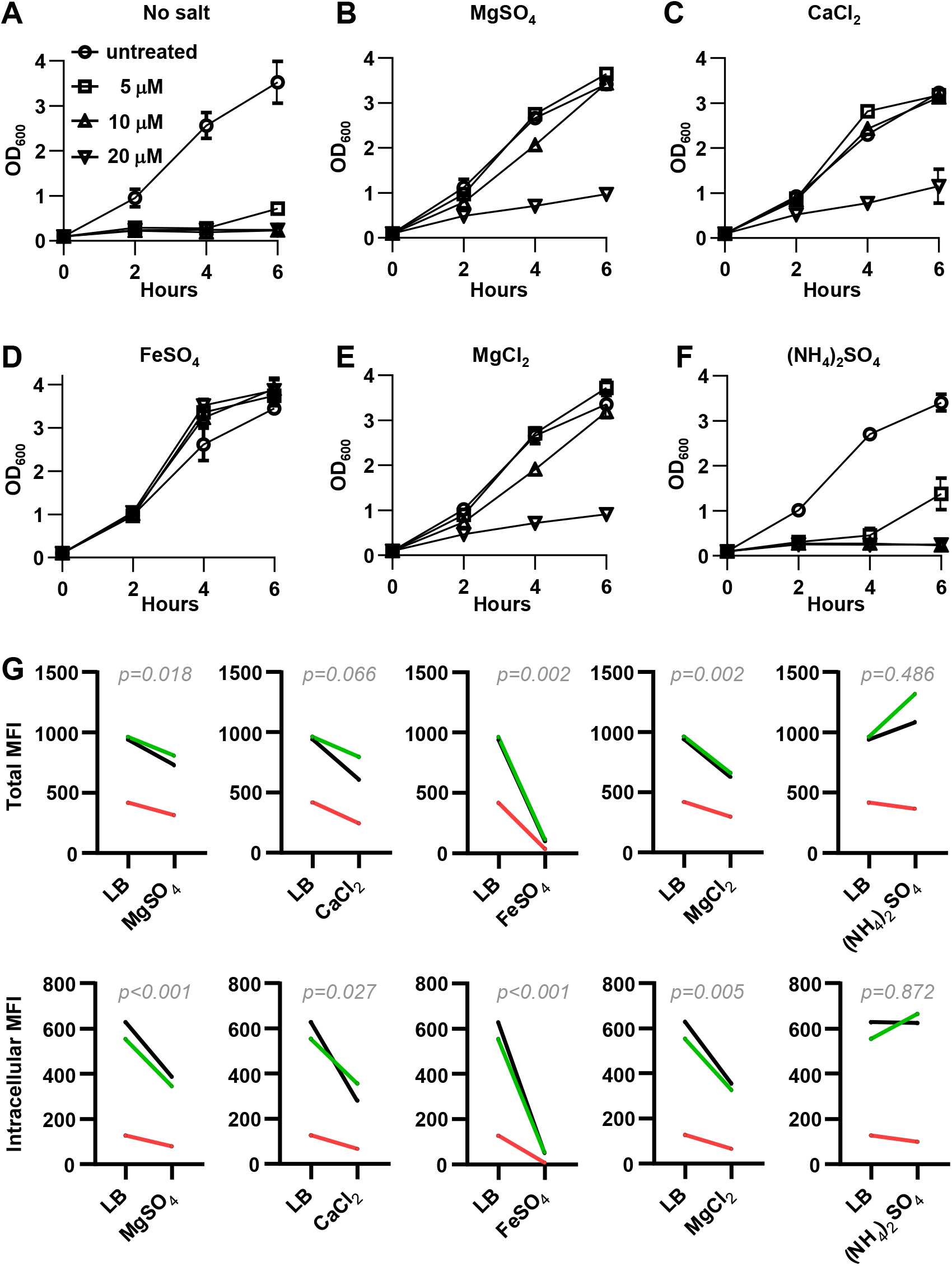
Divalent cations affect bacterial sensitivity towards TAT-RasGAP_317-326_ and decrease binding and entry of the peptide in *E. coli*. **(A-F)** *E. coli* MG1655 were grown overnight at 37°C in LB supplemented with 2 mM of the indicated salt and diluted to OD_600_ = 0.1. Bacterial suspension was then grown 1 hour at 37°C before addition of the indicated concentrations of TAT-RasGAP_317-326_. OD_600_ was measured at the indicated times after the initial dilution. The results correspond to the mean ± the range of two independent experiments. **(G)** *E. coli* MG1655 were grown overnight at 37°C in LB containing 2 mM of the indicated salts and diluted to OD_600_ = 0.1. Bacterial binding and uptake of 10 μM FITC-labelled TAT-RasGAP_317-326_ was recorded in triplicate (each shown with a different color on the graph) via flow cytometry with (Intracellular) or without (Total) quenching with 0.2 % trypan blue. P values were calculated by ratio paired t-test between the indicated condition and the LB control.

Would cations other than Mg^2+^ also render bacteria less sensitive to TAT-RasGAP_317-326_? Addition of Fe^2+^ and Ca^2+^ in culture medium decreased the sensitivity of *E. coli* to peptide (Fig. 1 panels C and D). We also found that high Mg^2+^ concentrations decreased bacterial sensitivity to TAT-RasGAP_317-326_ both in the context of sulfate and chloride counterions (Fig. 1 panels B and E). In contrast, ammonium sulfate did not affect bacterial susceptibility to TAT-RasGAP_317-326_ (Fig. 1F). Collectively, these results indicate that the Fe^2+^, Ca^2+^ and Mg^2+^ divalent cations in culture medium both hamper the ability of TAT-RasGAP_317-326_ to kill bacteria.

Since divalent cations are known to influence outer membrane characteristics, such as lipopolysaccharide (LPS) integrity (Hancock, 1997) we questioned whether TAT-RasGAP_317-326_ binding and internalization were altered by these cations. Fe^2+^, and to a lower extent Ca^2+^ and Mg^2+^, decreased the levels of FITC-labelled TAT-RasGAP_317-326_ bound to the surface of *E. coli* as well as the amount of internalized peptide (Fig. 1G). However, we found that peptide binding and internalization were not affected by ammonium sulfate – a finding that is consistent with our data that ammonium sulfate does not impact peptide MIC (Fig. 1F). Altogether, this data suggests that divalent cations decrease bacterial sensitivity to TAT-RasGAP_317-326_ peptide via a mechanism that restricts peptide binding and entry in bacteria.

### TAT-RasGAP_317-326_ is bactericidal against *E. coli* and *P. aeruginosa*

To determine the relationship between peptide exposure and the number of viable (culturable) bacteria, we performed colony formation unit (CFU) assays. For *E. coli* grown in LB, 10 μM of TAT-RasGAP_317-326_ induced an initial 2- to 5- fold decrease in the number of surviving bacteria but there was no further decrease upon longer incubation times (Fig. 2A). A more pronounced decrease in bacterial viability was observed at peptide concentrations ≥ 15 μM, indicating that the peptide is bactericidal at these concentrations (Fig. 2A). Confocal microscopy studies showed that *E. coli* accumulated TAT-RasGAP_317-326_ intracellularly when exposed to a concentration leading to bacterial killing (Fig. 2E). Furthermore, peptide exposure at this concentration led to changes in bacterial morphology as seen by electron microscopy (Fig. 2F). For *P. aeruginosa* grown in BM2 Mg^low^ medium, 0.5-2 μM TAT-RasGAP_317-326_ had a small impact on bacterial growth relative to the no peptide control, while 5-10 μM strongly reduced bacterial numbers (Fig. 2B). In order to analyze the kinetics of peptide activity at early time points, we performed survival curves using 20 μM of TAT-RasGAP_317-326_ peptide for *E. coli* and 10 μM for *P. aeruginosa*. These concentrations correspond to 2.5 times the MIC of TAT-RasGAP_317-326_ (Table 1) and were shown to kill a majority of bacteria (Fig. 2A-B). We monitored bacterial killing for the first two hours of peptide exposure and compared bacterial killing by TAT-RasGAP_317-326_ and polymyxin B, the latter also added at 2.5 times its MIC (2.5 μg/ml for *E. coli* and 10 μg/ml for *P. aeruginosa*). Interestingly, TAT-RasGAP_317-326_ displayed slow time-kill kinetics in comparison to polymyxin B against *E. coli* (Fig. 2C), suggesting that these two peptides have different killing mechanisms in this bacterial species. In *P. aeruginosa* however, the killing kinetics were similar between TAT-RasGAP_317-326_ and polymyxin B (Fig. 2D).

**Figure 2.**
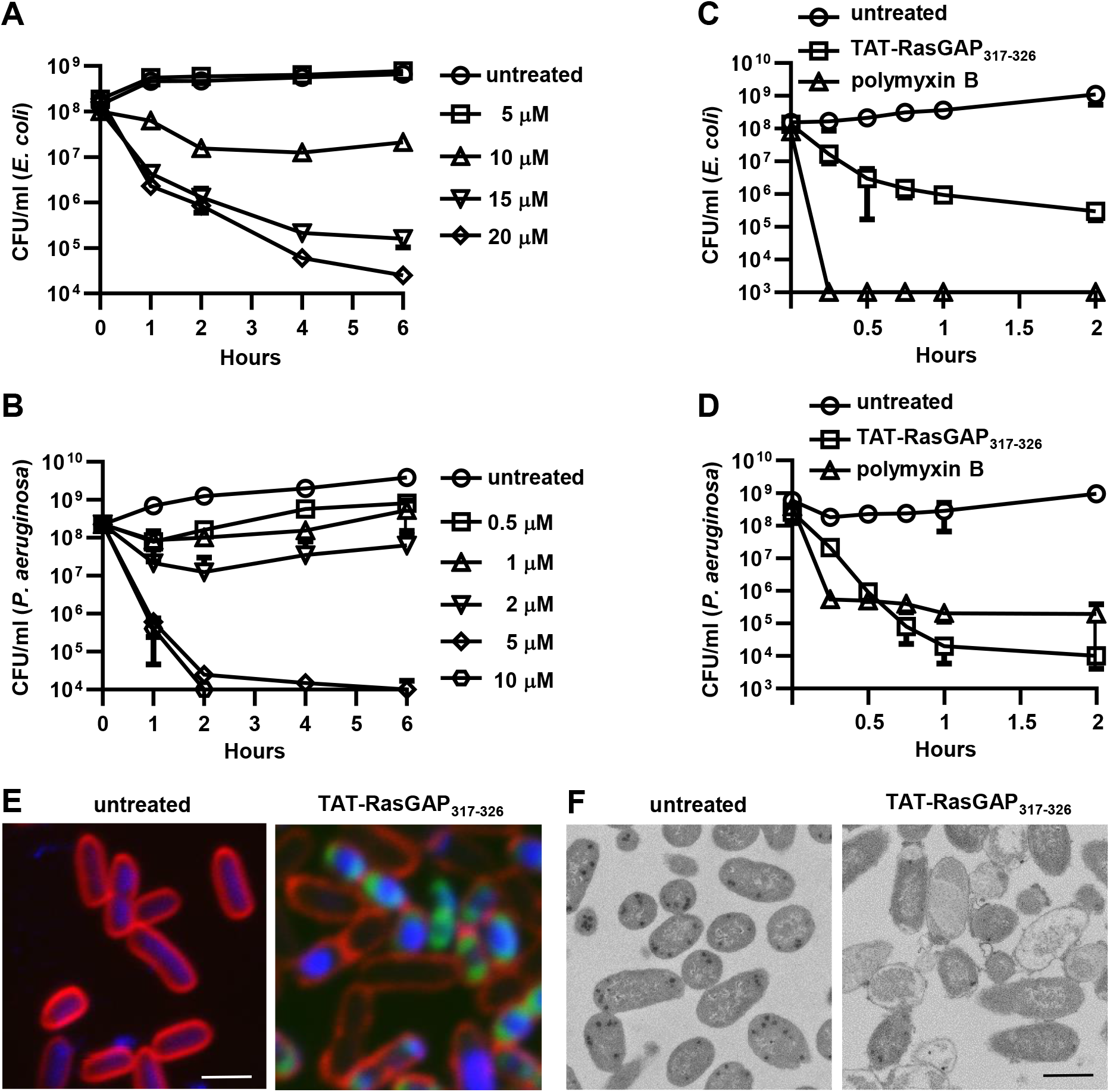
TAT-RasGAP_317-326_ is bactericidal against *E. coli* and *P. aeruginosa*. **(A-B)** Overnight cultures of *E. coli* MG1655 in LB (A) and *P. aeruginosa* PA14 in BM2 Mg^low^ (B) were diluted to OD_600_ = 0.1 and incubated at 37°C for 1 hour. TAT-RasGAP_317-326_ was then added at the indicated concentrations. Samples were taken at the indicated time points, serially diluted 10-fold in fresh LB and plated on LB agar plates. Number of colony forming units per ml (CFU/ml) in the original culture was calculated. **(C)** *E. coli* cultures were treated as in (A). TAT-RasGAP_317-326_ (20 μM) or polymyxin B (2.5 μg/ml) were added as indicated. Samples were taken at the indicated time points and CFU/ml were determined as in (A). **(D)** *P. aeruginosa* cultures were treated as in (B). TAT-RasGAP_317-326_ (10 μM) or polymyxin B (10 μg/ml) were added as indicated. Samples were taken at the indicated time points and CFU/ml were determined as in (A). Panels A-D: the results correspond to the mean ± standard deviation from at least two independent experiments. **(E)** *E. coli* MG1655 grown overnight and diluted to OD^600^ = 0.1 were incubated for 1 hour with or without 20 μM FITC-labelled TAT-RasGAP_317-326_ (green). The bacteria were then labelled with 5 μg/ml FM4-64 (red) and fixed with 4% paraformaldehyde. Incubation with DAPI (blue) was subsequently performed. Pictures were taken with a Zeiss LSM710 confocal microscope and analyzed using ImageJ software. Bar = 2 μm. **(F)** *E. coli* bacteria treated for 1 hour with 20 μM TAT-RasGAP_317-326_ were fixed with glutaraldehyde and prepared for electron microscopy as described in Material and Methods section. Samples were imaged via transmission electron microscopy. Images were analyzed using ImageJ software. Bar = 2 μm.

### TAT-RasGAP_317-326_ alters the transcriptional landscape of *E. coli*

RNA sequencing analysis was performed to evaluate the impact of TAT-RasGAP_317-326_ on *E. coli* transcriptome. For this, we used 10 μM of the peptide, a concentration that prevents *E. coli* proliferation but does not lead to a dramatic drop in bacterial numbers (Fig. 2A). Among the 4419 transcripts predicted from the *E. coli* MG1655 genome, 95.6% (n = 4223) were detected in at least one condition (Dataset 1). Figure 3A presents the fold change in gene expression between bacteria incubated with and without TAT-RasGAP_317-326_ as well as the average level of expression for each gene. We excluded from our analysis genes whose expression was below the threshold set at 16 reads per kilobase of transcripts per million reads (RPKM). Overall, TAT-RasGAP_317-326_ treatment affected the expression of 962 genes (fold change > 4): 11.0% of total detected genes were upregulated (red dots in Fig. 3A) while 11.8% were downregulated (blue dots in Fig. 3A). Detailed lists of upregulated and downregulated genes can be found in Supplementary Tables 1 and 2, respectively.

**Figure 3.**
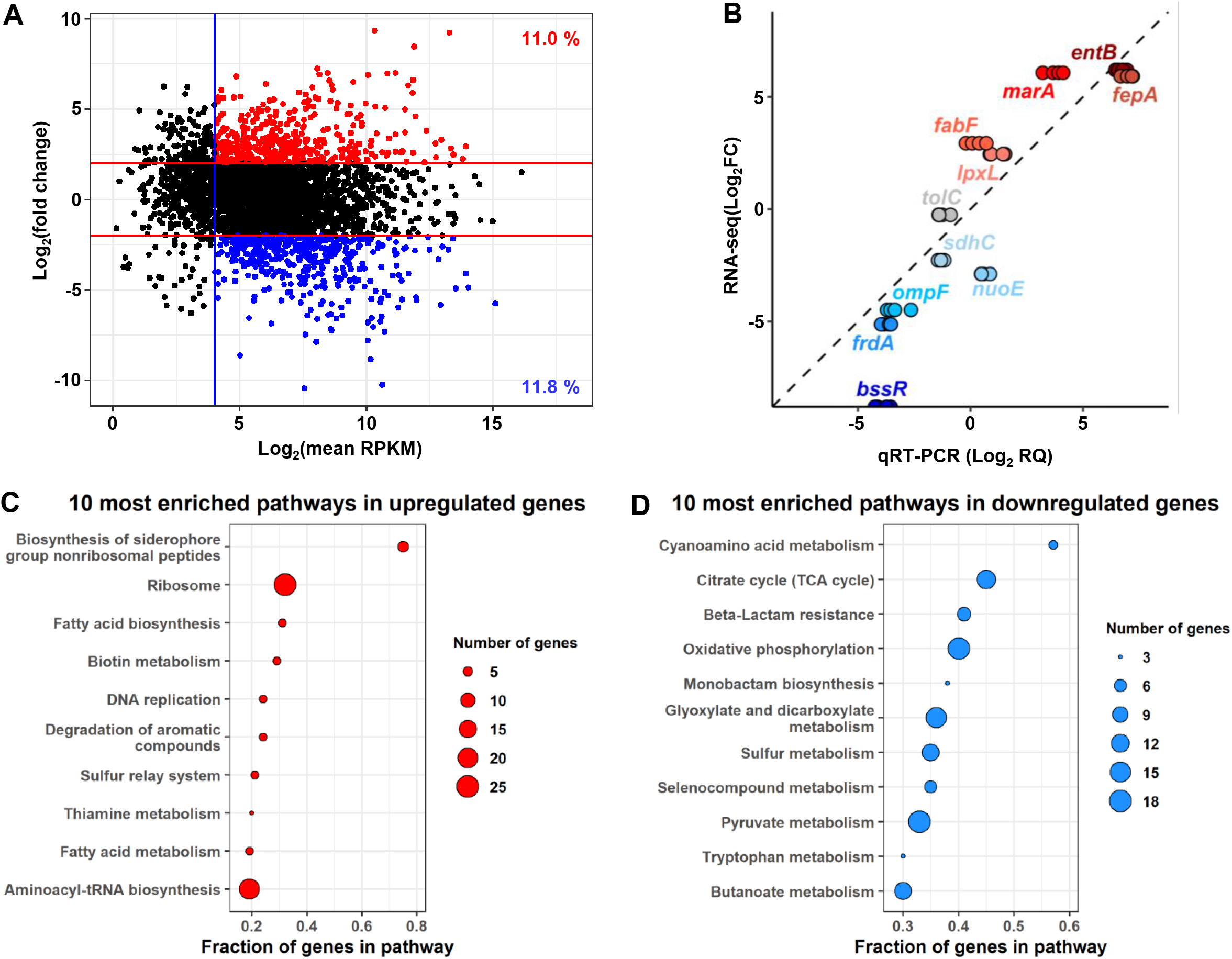
TAT-RasGAP_317-326_ alters the transcriptional landscape of *E. coli*. RNA-seq analysis was performed on *E. coli* MG1655 incubated for 1 hour with or without 10 μM TAT-RasGAP_317-326_. **(A)** MA-plot of the average gene expression (x-axis, RPKM: read per kilobase million) vs the differential expression (y-axis). Threshold for gene expression is indicated with the blue vertical line. The red lines indicate the cut-off limit for upregulated (red dots) and downregulated (blue dots) genes. **(B)** Correlation between RNA-seq (log_2_ Fold Change) and qRT-PCR (log_2_ Relative Quantification) differential expression performed on RNA extracted from *E. coli* treated for one hour with or without 10 μM TAT-RasGAP_317-326_ for a set of genes detected by RNA-seq as downregulated by the peptide (blue), not changed (grey) or upregulated (red). Gene expression was measured in duplicates on two independent extracted RNA sets. **(C-D)** Fraction of KEGG pathway genes that are upregulated **(C)** or downregulated **(D)** after treatment with TAT-RasGAP_317-326_. Dot size indicates the number of genes in the selection.

We assessed and validated twelve genes from the RNA-Seq data by qRT-PCR on RNA extracted under the same conditions as for the RNA-Seq analyses. Five of these (*lpxL*, *fabF*, *marA*, *entB,* and *fepA*, depicted in red in Fig. 3B) were reported by RNA-Seq as upregulated, five as downregulated (*bssR*, *frdA*, *ompF*, *nuoE,* and *sdhC*, depicted in blue in Fig. 3B) and two as unchanged according to the RNA-Seq analysis (*tolC* and *ompR*). One of the unchanged genes, *ompR*, was used as the housekeeping reference gene for normalization. We obtained good correlation between the fold changes obtained with RNA-Seq and with qRT-PCR, confirming the validity of the RNA-Seq data (Fig. 3B).

Using the gene expression profiles we obtained from RNA sequencing, we investigated which biological pathways were associated with the *E. coli* response upon exposure to TAT-RasGAP_317-326_. To accomplish this in a systematic manner, we performed Gene Ontology (GO) and Kyoto Encyclopedia of Genes and Genomes (KEGG) pathway enrichment analyses (Kanehisa and Goto, 2000, Kanehisa et al., 2019, Kanehisa, 2019, Ashburner et al., 2000, The Gene Ontology, 2019) on the subset of differentially expressed genes. The analysis of KEGG pathways revealed that several metabolic and information-processing pathways were enriched among differentially expressed genes (Fig. 3, panels C and D). For example, seven of the eight genes responsible for enterobactin synthesis in *E. coli* (included in “biosynthesis of siderophore group nonribosomal peptides” KEGG pathway) were upregulated upon peptide treatment. Other metabolic pathways such as carbon metabolism (citrate cycle, pyruvate metabolism) and oxidative phosphorylation were downregulated. Similarly, GO term analysis revealed that upregulated and downregulated genes in response to TAT-RasGAP_317-326_ were enriched in biological processes related to general bacterial metabolism and stress response (Supplementary Fig. 4). From our data, we could not distinguish peptide-specific gene expression changes from gene expression changes mediating general bacterial adaptation to stress. To address this question, we decided to perform a screening of a comprehensive *E. coli* deletion mutant library to determine which genes are directly involved in the bacterial response to TAT-RasGAP_317-326_.

### Screening of the Keio *E. coli* deletion mutant library uncovers genes that affect bacterial responses to TAT-RasGAP_317-326_

The Keio collection of *E. coli* deletion mutants consists of single gene deletion clones for each non-essential gene in *E. coli* (Baba et al., 2006). To perform the screening of the collection, we exposed each Keio strain to 5 μM TAT-RasGAP_317-326_, a non-bactericidal concentration of peptide (Fig. 2A) and monitored bacterial growth by OD_590_ measurement at specific time points (detailed results of growth measurements for all individual mutants are available in Dataset 2). For each strain, we determined the relative growth of the deletion strain compared to the wild-type strain when incubated with TAT-RasGAP_317-326_ for 6 hours and 24 hours (Fig. 4, panels A and B). We identified 27 strains showing decreased sensitivity to the peptide, thus having a normalized growth at 6 hours higher than the average of 270 replicates of the parental strain + 2 times the standard deviation (Fig. 4A and Supplementary Table 3). Furthermore, we identified 356 hypersensitive strains (having deletions in 279 different genes) that showed a normalized growth at 24 hours lower than average of the parental strain – 3 times the standard deviation (Fig. 4B). While the wild-type strains grew more slowly in the presence than in the absence of TAT-RasGAP_317-326_, strains showing decreased sensitivity to the peptide grew similarly in both conditions and hypersensitive strains showed no detectable growth in the presence of the peptide (Fig. 4C). It has to be mentioned that Keio collection is composed of two independent deletion mutants for each gene. We could not observe decreased sensitivity, as defined by our criteria, in both strains having the same gene deleted (Supplementary Table 3). However, some decreased sensitivity, approaching the threshold, could be observed in the second strain for a few genes such as *crr* and *rfaY*, for example. The *crr* gene product is involved in glucose uptake and phosphorylation, and in carbon metabolism regulation (Deutscher et al., 2006). The *rfaY* gene product is part of the LPS biogenesis pathway (Yethon et al., 1998). Interestingly, inactivation of *rfaY* by transposon mutagenesis was shown to affect *E. coli* susceptibility to another AMP, LL-37 (Bociek et al., 2015).

**Figure 4.**
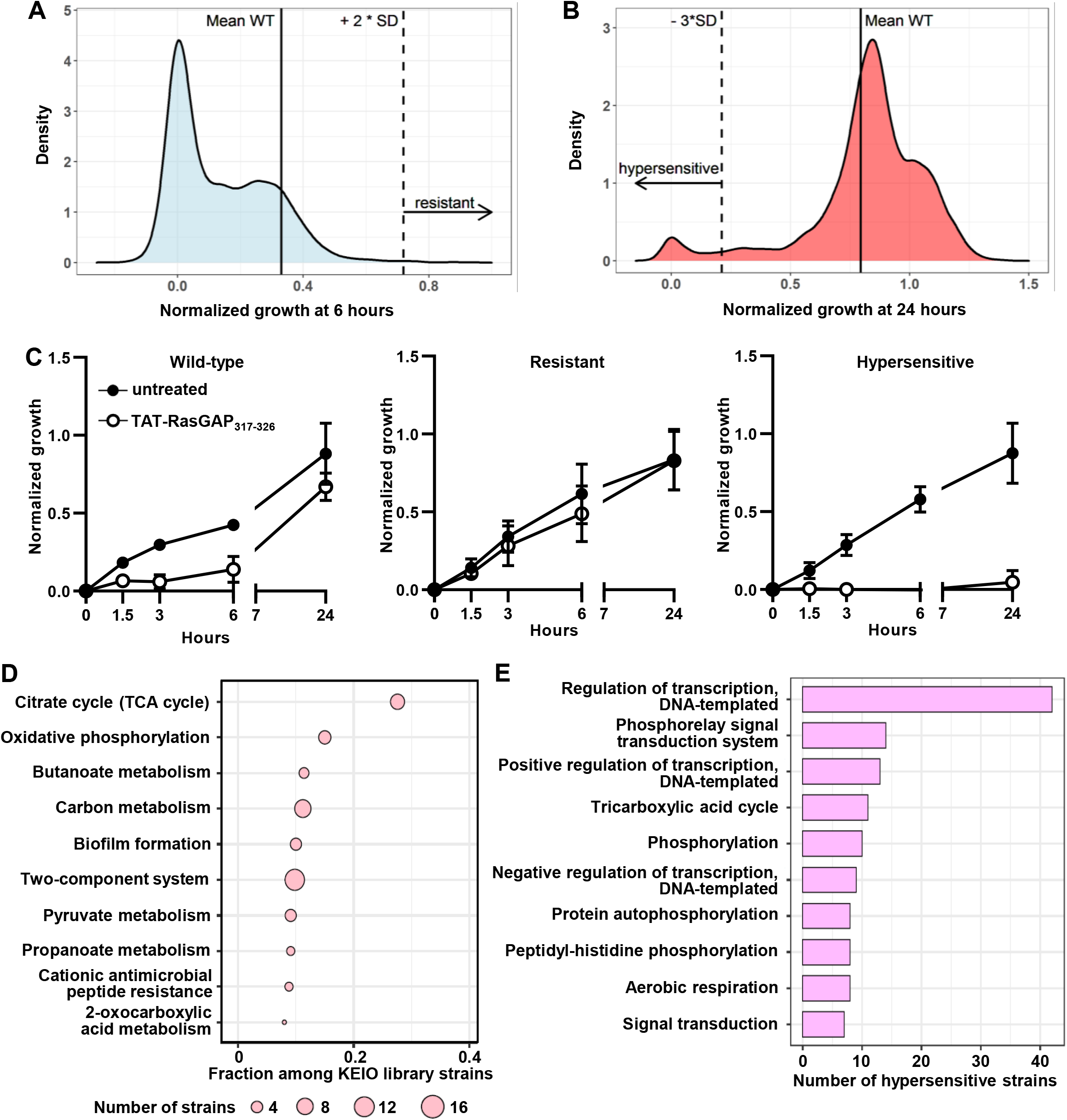
Selection of hypersensitive and resistant *E. coli* deletion mutants from the KEIO collection. Deletion mutants and the corresponding wild-type strain were grown in LB medium with or without 5 μM TAT-RasGAP_317-326_. OD_590_ was measured at 0, 1.5, 3, 6, and 24 hours. **(A-B)** Distribution of the normalized growth (NG; see the methods for the calculation of NG) of bacteria incubated with TAT-RasGAP_317-326_ at 6 hours **(A)** and 24 hours **(B)**. The mean NG of the wild-type strain (mean WT) is indicated with a vertical solid line. Strains with NG_6 hours_ > [mean WT + 2 standard deviations (SDs)] and with NG_24 hours_ < [mean WT - 3 SDs] are defined here as resistant and hypersensitive strains, respectively. **(C)** Growth curves of wild-type (n=270), hypersensitive (n=356) and resistant (n=20) mutants in presence or absence of 5 μM TAT-RasGAP_317-326_. Data are mean ± SD. **(D)** Top 10 most represented KEGG pathways among hypersensitive strains. The number of hypersenstive strains in each pathway was normalized to the number of KEIO collection strains in the corresponding pathway. **(E)** Biological processes GO term enrichment analysis with the 10 most represented terms among the hypersensitive strains.

On the other hand, 77 gene deletions caused hypersensitivity for both replicates present in the Keio collection (Supplementary Table 4). KEGG pathway and GO term analyses were thus performed on genes for which both deletion mutants showed hypersensitivity. The results of this analysis indicate that deletion of genes involved in bacterial metabolism and two component systems were associated with TAT-RasGAP_317-326_ bacterial sensitivity (Fig. 4D-E, Table 2).

**Table 2:** List of two-component systems, for which deletion of at least one of the components caused increased sensitivity to TAT-RasGAP_317-326_. Data were extracted from the Keio collection screening and genes annotated as two-component system components and showing increased sensitivity for both duplicates were selected. Systems are highlighted in bold when both components were retrieved in the screening. Conditions regulating the systems were extracted from the Ecocyc.org database.

| Two-component system | Component(s), which deletion cause(s) sensitivity | Conditions regulating the system | Potential link(s) with resistance (Ref) |
| --- | --- | --- | --- |
| *baeSR* | *baeS* (Sensory kinase) | Envelope stress | Overexpression of *baeR* causes resistance to novobiocin and deoxycholate (Baranova and Nikaido, 2002) |
| ***citAB*** | *citB* (DNA binding) | Citrate/anaerobic conditions |  |
| *cpxAR* | *cpxA* (Sensory kinase) | Inner membrane stress | Upregulates multidrug resistance cascade (Weatherspoon-Griffin et al., 2014) |
| ***creCB*** | **Both** | Carbone source | Hyperactivation causes colicin E2 tolerance (Cariss et al., 2010) |
| ***dcuSR*** | **Both** | Dicarboxylate |  |
| ***envZ-ompR*** | **Both** | Medium osmolality |  |
| *kdpDE* | *kdpD* (Sensory kinase) | Potassium concentration |  |
| ***phoQP*** | **Both** | Low magnesium | Involved in resistance to AMPs (Bader et al., 2005) |
| ***rcsBC*** | *rcsC* (Sensory kinase) | Envelope stress | Contributes to intrinsic antibiotic resistance (Laubacher and Ades, 2008) |

Of interest, we found that a subset of less and more sensitive Keio strains were deletion mutants in LPS biogenesis genes. We confirmed differences in sensitivity for ΔrfaY and ΔlpxL deletion mutants by measuring MIC and IC_50_ of TAT-RasGAP_317-326_ on these mutants and could confirm that deletion of *rfaY* caused a decreased sensitivity and deletion of *lpxL* an increased sensitivity to the peptide (Fig. 5A). This raises the possibility that TAT-RasGAP_317-326_ directly interacts with bacterial LPS. In such a case, soluble LPS should compete with the peptide for binding to bacterial cells and reduce peptide efficacy, an effect that has been reported for polymyxin B (Domingues et al., 2012). Figure 5B shows indeed that soluble LPS greatly diminishes the efficacy of polymyxin B but has no impact on the sensitivity of *E. coli* towards TAT-RasGAP_317-326_. The potential role played by genes involved in LPS synthesis in TAT-RasGAP_317-326_ sensitivity remains therefore to be uncovered.

**Figure 5.**
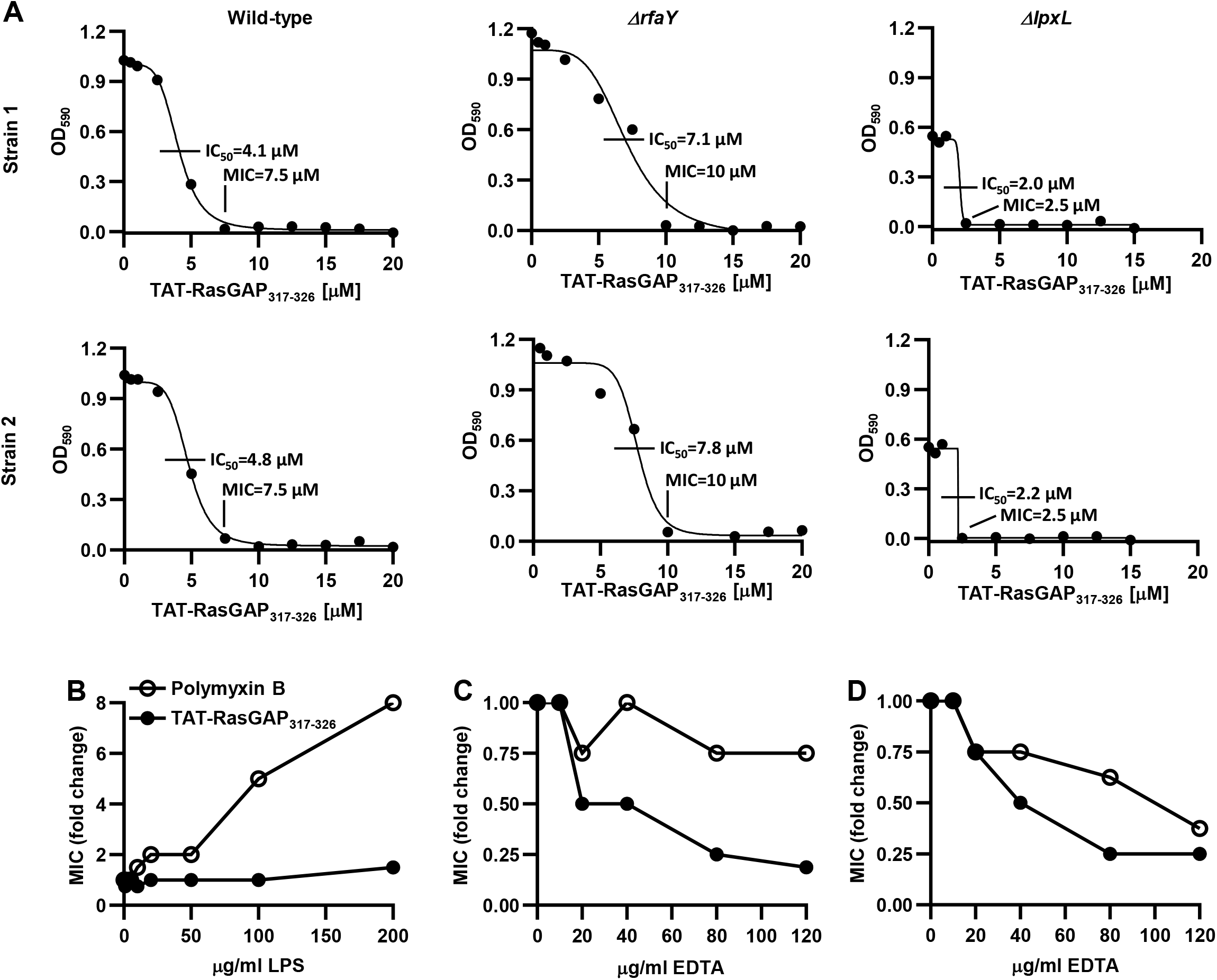
Changes in LPS integrity influence TAT-RasGAP_317-326_ activity. **(A)** Deletion of LPS biosynthesis genes have diverse effect on TAT-RasGAP_317-326_ activity. MICs and IC_50_ of TAT-RasGAP_317-326_ against wild-type strain or the two deletion mutants ΔrfaY resp. ΔlpxL from the Keio deletion library were measured as previously described. **(B-D)** LPS supplementation or EDTA differentially influence activity of TAT-RasGAP_317-326_ and polymyxin B. MICs of TAT-RasGAP_317-326_ and polymyxin B on *E. coli* MG1655 (B-C) or ATCC25922 (D) were measured as previously described in LB containing the indicated concentrations of purified LPS or EDTA. Data are averages of two independent experiments.

We further investigated whether LPS integrity was required for survival of *E. coli* in the presence of TAT-RasGAP_317-326_. For this purpose, we used EDTA to destabilize the LPS structure (Hancock, 1984) and measured how this impacted the MIC of TAT-RasGAP_317-326_ and polymyxin B on two *E. coli* strains lacking (the MG1655 strain) or not (the ATCC 25922 strain) O-antigen moieties (Eder et al., 2009). EDTA, at concentrations that do not affect bacterial proliferation (Supplementary Fig. 5), sensitized both strains to TAT-RasGAP_317-326_ (Figure 6B-C), suggesting that compromised LPS integrity favors the antimicrobial activity of the peptide. Polymyxin B sensitivity was less affected by EDTA, indicating again that polymyxin B and TAT-RasGAP_317-326_ use different mechanisms to inhibit bacterial growth or survival.

**Figure 6.**
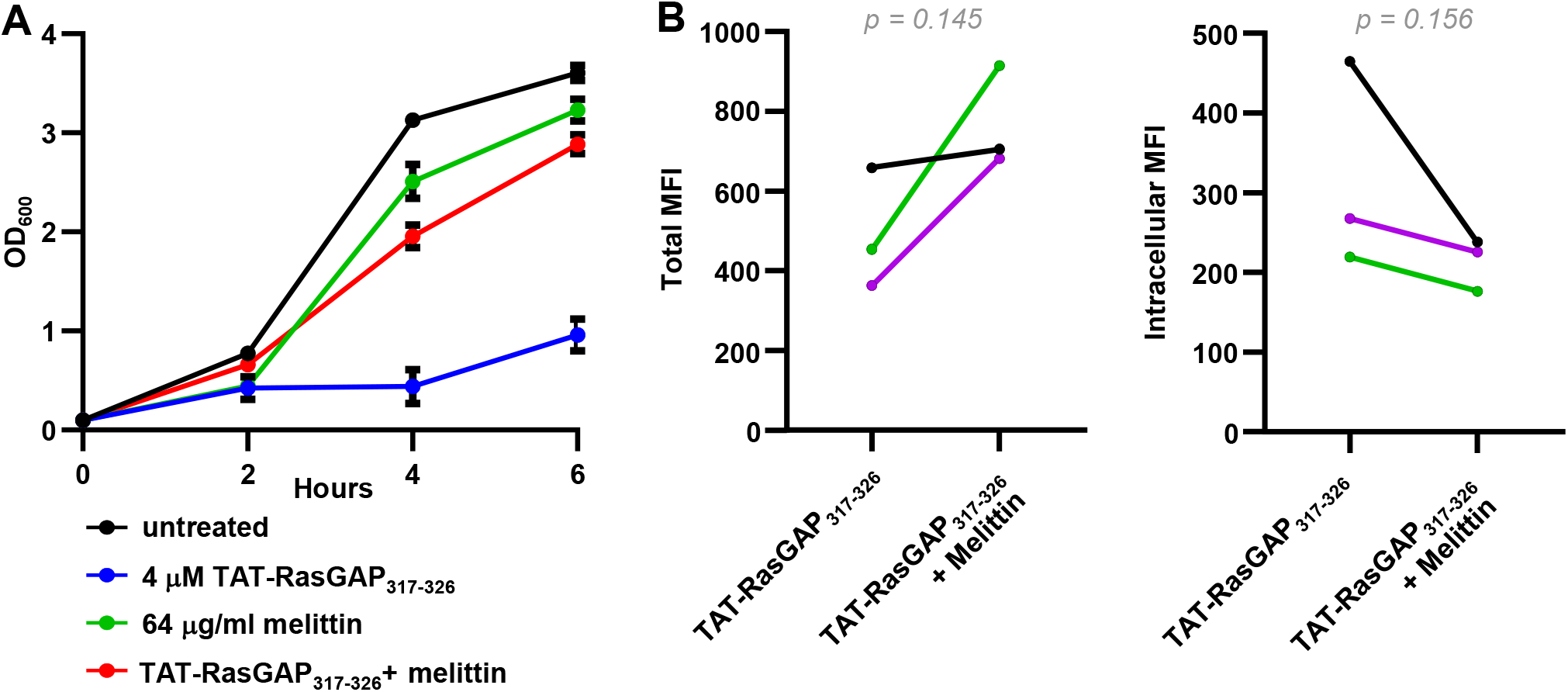
Melittin has an inhibitory effect on TAT-RasGAP_317-326_ activity. **(A)** Sub-inhibitory concentrations of melittin interfere with TAT-RasGAP_317-326_ activity. Indicated concentrations of AMPs were added and OD_600_ was measured as previously described. Average and range of two independent experiments are shown. **(B)** *E. coli* MG1655 was grown overnight at 37°C, diluted to OD_600_ = 0.1 and grown during 1 hour before addition or not of 10 μM FITC-labelled TAT-RasGAP_317-326_ with or without 64 μg/ml melittin. Cells were incubated for 1 hour at 37°C, extracellular fluorescence was quenched (Intracellular) or not (Total) using 0.2% trypan blue before sample acquisition. Mean fluorescence intensities (MFI) were measured for triplicates (shown with different colors). P values were calculated using ratio paired t-test between the indicated conditions.

### Transposon screening in *P. aeruginosa*

Since TAT-RasGAP_317-326_ is active against both *E. coli* and *P. aeruginosa*, we investigated whether some of the pathways that play a role in peptide resistance are shared between the two bacterial species. We thus exposed a *P. aeruginosa* transposon mutant library (Vitale et al., 2020) to 0.5 μM TAT-RasGAP_317-326_ for 12 generations and performed deep sequencing. This allowed us to compare level of transposons in different genes between a bacterial population treated with the peptide and another that was not. Prevalence of strains having a disruption of a gene required for growth in presence of the peptide would decrease compared to strains having integrated the transposon in an unrelated region (detailed results of this deep sequencing are presented as Dataset 3). We thus defined lower prevalence of transposon insertion as a read-out of hypersensitivity to TAT-RasGAP_317-326_. By this way, we identified 75 genes, for which prevalence of disruption via transposon insertion decreased in presence of the peptide (Supplementary Table 5). Interestingly, 26 of these (35%) are associated with hypersensitivity to other antimicrobial peptides (Vitale et al., 2020). Some of these genes code for LPS modifying enzymes such as ArnA, ArnB and ArnT, and for two-component regulators such as ParS and ParR that are involved in the regulation of LPS modifications (Fernandez et al., 2010). Among the genes, for which prevalence of transposon insertion was decreased in presence of TAT-RasGAP_317-326_ but not with other AMPs, we identified *algJ*, *algK* and *algX*, genes of the biosynthesis pathway of the extracellular polysaccharide alginate. We also observed that mutants in genes coding for the RND efflux transporter MdtABC and CusC, a component of the trans-periplasmic Cu^2+^ transporter CusCFBA, are potentially associated with hypersensitivity to TAT-RasGAP_317-326_. Other pathways that seem to be important for TAT-RasGAP_317-326_ resistance are related to carbon metabolism, redox reactions and translation regulation (Supplementary Table 5).

We next compared the lists of potential hypersensitive strains found in screenings in *E. coli* and in *P. aeruginosa*. We identified six gene orthologues, whose disruption is associated with hypersensitivity to the TAT-RasGAP_317-326_ peptide in both *E. coli* and *P. aeruginosa* (Table 3). Among them, four are coding for two-component system proteins: *parR* and *parS* (*rtsA* and *rstB* in *E. coli*), *phoP*, and *pmrB* (*qseC* in *E. coli*). These four mutants were associated with hypersensitivity to polymyxin B in *P. aeruginosa* (Vitale et al., 2020), indicating that these regulatory pathways may be required for a general adaptation to AMPs. This is of interest, since RstAB system is regulated by PhoQP system in *E. coli* (Ogasawara et al., 2007) and PhoQP system was shown to be involved in resistance to AMPs (Yadavalli et al., 2016). Two other genes conserved between *P. aeruginosa* and *E. coli* are associated specifically with hypersensitivity to TAT-RasGAP_317-326_. One is a transcriptional regulator and the other is involved in LPS biosynthesis, further highlighting a potential role for cell surface composition in the sensitivity of bacteria towards TAT-RasGAP_317-326_ (Table 3).

**Table 3:** Genes found as more sensitive in both *P. aeruginosa* by transposon library screening and in *E. coli* by Keio collection screening. List of the genes, whose disruption generated mutant strains found as sensitive in transposon library screening in *P. aeruginosa* and whose orthologues in *E. coli* were detected as sensitive in Keio collection screening. Transposon mutants of *P. aeruginosa* were incubated in presence or absence of 0.5 μM TAT-RasGAP_317-326_ in BM2 Mg^low^ medium for 12 generations. Transposon junctions were amplified and sequenced. Fold change (FC) between abundance of transposon mutants with incubation in absence (BM2) or presence (TATRasGAP) of the peptide was calculated and values are presented as Log_2_ of the FC. Inf indicates that no mutant was detected upon peptide treatment, and therefore Log_2_ of the FC could not be calculated. Shown also are results obtained in a former study using the same transposon library that indicate which gene disruptions also cause hypersensitivity to polymyxin B (Vitale et al., 2020). Ratio in Keio screening column indicates whether sensitivity was detected in one (1/2) or both (2/2) replicates of the Keio collections.

| Locus_tag | Gene_symbol | Description | Category | Log<sub>2</sub>(FC_BM2vsTAT-RasGAP) | Hypersensitivity to Polymyxin B | *E. coli* homologues |
| --- | --- | --- | --- | --- | --- | --- |
| PA14_41260 | *parR* | two-component response regulator | Two-component regulator system | -4.268533546 | Yes | *rstA* |
| PA14_41270 | *parS* | two-component sensor | Two-component regulator system | Inf | Yes | *rstB* |
| PA14_49180 | *phoP* | two-component response regulator PhoP | Two-component regulator system | Inf | Yes | *phoP* |
| PA14_63160 | *pmrB* | PmrB: two-component regulator system signal sensor kinase PmrB | Two-component regulator system | -3.492272393 | Yes | *qseC* |
| PA14_02390 |  | transcriptional regulator | Transcription regulation | -5.436478184 |  | *cynR* |
| PA14_71970 | *wbpW* | GDP-mannose pyrophosphorylase | LPS biosynthesis | -4.335647742 |  | *cpsB* |

### Effect of combining TAT-RasGAP_317-326_ with other AMPs

To determine whether TAT-RasGAP_317-326_ activity is affected by other AMPs, we performed growth tests of *E. coli* in presence of different combinations of TAT-RasGAP_317-326_, melittin, LL-37 and polymyxin B. Concentrations were chosen so that a clear difference in growth was observed when “half” concentrations were used (Supplementary Fig. 6A) as compared with “full” concentrations, that correspond to the double of “half” concentrations (Supplementary Fig. 6B). The effect of combining pairs of AMPs using “half” concentrations is shown in Supplementary Fig. 6C as percentage of growth compared to an untreated control. We did not observe an increase in the effect of TAT-RasGAP_317-326_ when combined with the three other AMPs. However, the combination of melittin and polymyxin B (2.7% of growth) and the combination of LL-37 and polymyxin B (0.6% of growth) showed increased activity. Notably, the combination of melittin and polymyxin B caused stronger growth inhibition (>95%) than obtained by either compound at the “full” concentration (~20% and ~35% growth inhibition for melittin and polymyxin B, respectively; Supplementary Fig. 6C). This observation is consistent with previous reports of the synergism between melittin and antibiotics such as doripenem and ceftazidime (Akbari et al., 2019). In contrast, we observed an apparent lower effect of TAT-RasGAP_317-326_ in presence of melittin (Supplementary Fig. 6C). Since this effect was very weak in these conditions, we combined the “half” concentration of melittin with the “full” concentration of TAT-RasGAP_317-326_ and could observe a clear inhibition of the antimicrobial activity of this peptide by melittin (Fig. 6A). To better understand the mechanism behind this observation, we assessed peptide binding and entry into bacteria using a fluorescently labelled version of TAT-RasGAP_317-326_ peptide. We found that, in the presence of melittin, binding of FITC-labelled TAT-RasGAP_317-326_ to *E. coli* bacteria was not decreased, but apparently slightly increased when compared to the control condition where bacteria were incubated with FITC-labelled TAT-RasGAP_317-326_ alone. However, we observed an apparent lower intracellular accumulation of the labelled version of TAT-RasGAP_317-326_ peptide in presence of melittin (Fig. 6B).

### *In vitro* selection of resistant bacteria to TAT-RasGAP_317-326_ peptide

AMPs are less susceptible to bacterial resistance evolution than classical antibiotics (Lazar et al., 2018, Spohn et al., 2019, Lazzaro et al., 2020). To measure the propensity of bacteria to develop resistance against TAT-RasGAP_317-326_, we serially passaged several bacterial strains (*E. coli*, *P. aeruginosa*, *S. aureus* and *S. capitis*) in the presence of TAT-RasGAP_317-326_ peptide and recorded the number of passages required to detect the appearance of resistant mutants in each strain. First, we grew the parental bacterial strains overnight in presence of sub-inhibitory concentrations of the peptide. We then diluted this parent culture into two subcultures, one of which was exposed to an increased concentration of the TAT-RasGAP_317-326_ peptide while the other was kept in the same concentration of peptide as the parent culture. Once bacterial growth was detected in the culture exposed to an elevated concentration of the peptide, the process was repeated, thereby exposing the bacterial culture to sequentially increasing concentrations of peptide for a total of 20 passages. For each passage, we measured the corresponding MIC (Fig. 7A). Using this approach, we obtained strains with highly increased MICs (16-32 fold) for *E. coli, S. capitis*, but only a faint increase (2-4 fold) for *P. aeruginosa* (Fig. 7A, Table 4 and Supplementary Tables 6-8). It should be noted that the parental strain of *S. aureus* has a peptide MIC in the range 64-128 μM and this MIC rapidly increased to 256 μM (Supplementary Table 9). We did not expose bacteria to higher concentrations, as the peptide started to precipitate in these conditions. To test whether the strains recovered at passage 20 for *E. coli*, *P. aeruginosa* and *S. capitis* and passage 12 for *S. aureus* showed increased resistance to other AMPs as well, we determined the fold change of the MICs for polymyxin B, melittin and LL-37 relative to the corresponding parental strains that did not undergo selection (Table 4). Interestingly, peptide-resistant *E. coli* (gram-negative) did not show increased MICs to the other AMPs we tested as compared to the parental strain. In contrast, *P. aeruginosa* and the Gram-positive *S. aureus* and *S. capitis* selected for resistance to TAT-RasGAP_317-326_ showed increased MIC towards other AMPs (Table 4). Thus, our findings suggest that bacterial species differ in their tendency to develop cross-resistance to TAT-RasGAP_317-326_ peptide and other AMPs.

**Figure 7.**
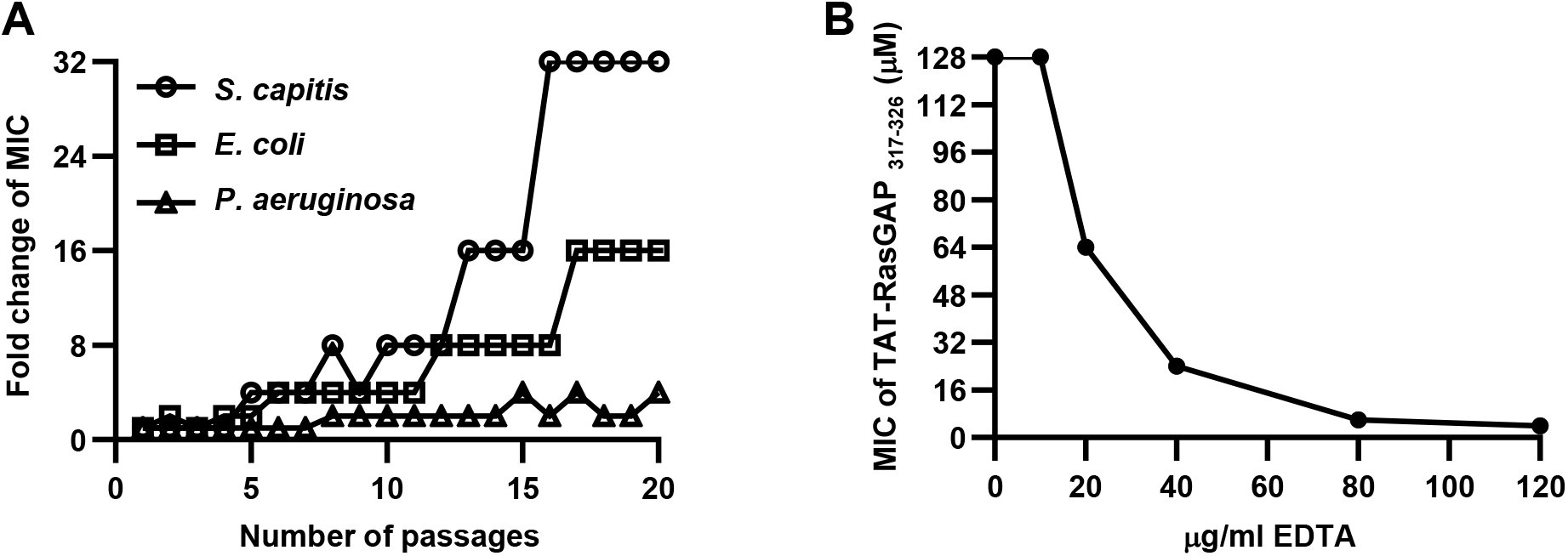
Bacterial resistance against TAT-RasGAP_317-326_ appears after selection with sub-inhibitory concentrations of peptide. **(A)** The indicated strains were incubated in presence or absence of 0.5 MIC of TAT-RasGAP_317-326_. Cultures were then diluted each day in medium containing either the same concentration of the peptide or twice the concentration. Once bacterial growth was detected in the culture exposed to an elevated concentration of the peptide, the process was repeated thereby exposing the bacterial culture to sequentially increasing concentrations of peptide for a total of 20 passages. MIC of each passage was then measured and is presented as a fold change compared to the MIC of the original strain passaged in the absence of peptide. **(B)** Peptide-resistant *E. coli* is susceptible to peptide activity during combination treatment with EDTA. MIC of *E. coli* strain selected for 20 passages from (A) was measured in presence of increasing concentrations of EDTA. Average of two independent experiments is presented.

**Table 4:** MICs of TAT-RasGAP_317-326_-resistant strains towards other AMPs. Fold change of MICs between the original strains (*E. coli* MG1655, *P. aeruginosa* PA14, *S. capitis* and *S. aureus* ATCC 29213) and strains exposed to increasing concentrations of TAT-RasGAP_317-326_ for 20 passages. MICs were measured as described for Fig. 1. n.d.: MIC of the strain could not be determined.

|  | TAT-RasGAP<sub>317-326</sub> | Polymyxin B | Melittin | LL-37 |
| --- | --- | --- | --- | --- |
| ***E. coli*** | 16 | 0.5 | 0.5 | 0.5 |
| ***P. aeruginosa*** | 4 | 2 | >2 | n.d. |
| ***S. capitis*** | 32 | >2 | 2 | n.d. |
| ***S. aureus*** | >2 | n.d. | 8 | n.d. |

Finally, we sought to investigate whether the peptide-resistant bacteria we obtained in our selection process remain targets for alternative treatments such as combination therapy with other antimicrobial agents. In particular, we tested whether EDTA, an agent known to enhance the efficacy of antimicrobials via a mechanism that weakens the outer cell wall of bacteria, could potentiate the effect of TAT-RasGAP_317-326_ against peptide-resistant *E. coli* (Leive, 1965). Importantly, the presence of EDTA alone at the concentrations tested was not associated with any significant change in bacterial numbers (Supplementary Figure 5). However, EDTA in combination with TAT-RasGAP_317-326_ (Fig. 7B) potentiated the ability of the peptide against the peptide-resistant *E. coli* strain. Our findings suggest that peptide-resistance remains treatable in combination therapy with other antimicrobial agents.

## Discussion

The activity of antimicrobial peptides can be affected by environmental factors, but we lack knowledge about how extracellular factors impact TAT-RasGAP_317-326_ activity. Here, we report that addition of divalent cations in LB medium resulted in decreased bacterial sensitivity to TAT-RasGAP_317-326_ peptide and reduced peptide binding and entry. The mechanism contributing to lower peptide binding might be due to competition between divalent ions in the culture medium and the cationic TAT-RasGAP_317-326_ peptide for binding to bacterial surface (Fig. 9). Alternatively, divalent cations, which are important for membrane stability, may influence binding and entry of TAT-RasGAP_317-326_ (Clifton et al., 2015).

RNA sequencing showed that genes involved in carbon metabolism were downregulated upon treatment with TAT-RasGAP_317-326_ (Fig. 3). Moreover, deletion or transposon mutants of genes involved in carbon metabolism and ATP production were more sensitive towards TAT-RasGAP_317-326_ (Fig. 4 and Table 3), indicating that energy production pathways may be important for resistance towards this peptide. Adaptation to environmental stimuli might also be of importance for survival to TAT-RasGAP_317-326_, since mutants lacking genes coding for some two-component systems show increased sensitivity towards TAT-RasGAP_317-326_ (Table 2). Several of these two-component systems are known to play role in resistance to antibiotics or AMPs, such as PhoPQ, whose importance in response to AMPs is well described (Bader et al., 2005, Yadavalli et al., 2016). This further highlights the importance of two-component systems for adaptability and survival of bacteria in harsh conditions.

Another pathway that may be involved in sensitivity to TAT-RasGAP_317-326_ is the LPS biosynthesis pathway. We found that some mutations affecting this pathway cause either moderate resistance or hypersensitivity to the peptide (Fig. 5A). We could further confirm the importance of LPS integrity for survival to TAT-RasGAP_317-326_ using EDTA that destabilizes LPS. This is consistent with the protective effect of divalent cations (Fig. 1), which can bind and stabilize LPS (Pelletier et al., 1994). Importance of bacterial surface composition in sensitivity towards TAT-RasGAP_317-326_ is further highlighted by the fact that *P. aeruginosa* transposon mutants affecting the alginate biosynthesis pathway are more sensitive to TAT-RasGAP_317-326_ than the control strain (Supplementary Table 5). Alginate is an anionic extracellular polysaccharide that is involved in virulence, antimicrobial resistance and biofilm formation in *P. aeruginosa* (Franklin et al., 2011).

Interestingly, screening of the Keio deletion collection did not allow to unearth mutants showing complete resistance towards TAT-RasGAP_317-326_. This indicates that resistance may not be obtained by the loss of function of one gene. Resistance towards TAT-RasGAP_317-326_ that we obtained by selection (Fig. 7A) may thus have acquired point mutations that modulate activity through activation of some pathways or modifications of essential components. This needs now further investigations in order to describe mechanisms of resistance towards TAT-RasGAP_317-326_ in particular and AMPs in general.

On the other hand, Keio collection screening highlighted pathways that are apparently required for *E. coli* to respond to TAT-RasGAP_317-326_. Whether these pathways are specifically required for response to TAT-RasGAP_317-326_ or play a role in a general response to AMPs needs further investigation. Interestingly, we observed, using a *P. aeruginosa* transposon mutants library, that 21% (16 out of 75) of the genes which mutation was association with hypersensitivity to TAT-RasGAP_317-326_ were associated with hypersensitivity towards other AMPs (Supplementary Table 5)(Vitale et al., 2020).

Combinatorial therapies are gaining interest in the treatment of multi-resistant bacteria (Leon-Buitimea et al., 2020). We thus investigated whether combination with other AMPs might influence the activity of TAT-RasGAP_317-326_. In general, activity of TAT-RasGAP_317-326_ was not influenced by other AMPs. However, melittin had an inhibitory effect on TAT-RasGAP_317-326_ activity, affecting its entry in bacteria (Fig. 6). This rather peculiar effect might be explained by the hypothesized mode of action of melittin (i.e. carpet model), in which melittin first interacts with the bacterial surface, before reaching a concentration threshold that leads to the disruption of the bacterial membrane (Lee et al., 2013). Sub-inhibitory concentrations of melittin might thus block binding of TAT-RasGAP_317-326_ to the bacterial membrane.

Finally, we investigated the potency of bacteria to develop resistance towards TAT-RasGAP_317-326_. Resistance could be obtained upon passages in sub-inhibitory concentrations of the peptide (Fig. 7A), but bacterial strains differed with respect to the rate of resistance emergence. Interestingly, peptide-resistant *E. coli* remains treatable by peptide in combination with EDTA, a chemical agent that compromises the integrity of the bacterial outer membrane. Future work should examine the mechanism of *E. coli* resistance to peptide and will help elucidate how EDTA, which targets the bacterial envelope, helps potentiate peptide activity in resistant backgrounds. Overall, our data highlight the potential benefit of combination therapies, which might not only prevent the development of such resistance, but also potentiate treatment of resistant strains, as shown here by EDTA in combination with TAT-RasGAP_317-326_.

The schemes presented in Figure 8 highlight the factors that may influence TAT-RasGAP_317-326_ activity and present hypotheses about underlying mechanisms. The positively charged TAT-RasGAP_317-326_ peptide interacts with the negative surface charges of the bacterial membrane, allowing its binding and entry in the bacterial cell (Fig. 8A). Presence of divalent cations in the culture medium compete with TAT-RasGAP_317-326_ peptide for binding to the negative charges on LPS, lowering the activity of the peptide. Similarly, modifications of LPS structure can also lower interaction between TAT-RasGAP_317-326_ and bacterial surface. We hypothesize this lower activity to be due to a decrease of the net charge of bacterial surface, causing a lower affinity of the peptide to bacteria (Fig. 8B). In contrast, destabilization of LPS by EDTA or by deletion of genes involved in biosynthesis of LPS precursors increases the bactericidal activity of TAT-RasGAP_317-326_. This is possibly due to a defect of the integrity of the bacterial envelope, decreasing bacterial defenses towards TAT-RasGAP_317-326_. (Fig. 8C).

**Figure 8.**
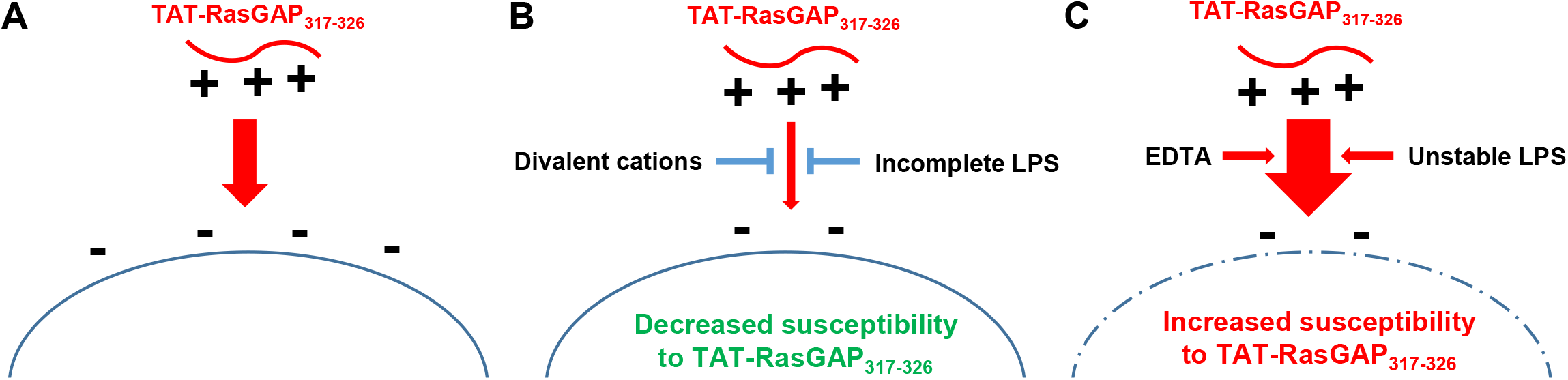
Model of interaction of TAT-RasGAP_317-326_ with bacterial surface. **(A)** The positively charged peptide interacts with negative charges on bacterial surfaces. **(B)** This interaction may be lowered by presence of divalent cations, which compete for the negative charges of the LPS, or by mutations that decrease the net negative charge of LPS. **(C)** Chemicals that target the bacterial outer membrane, such as EDTA and bacterial mutants with defects in LPS biosynthesis are associated with increased susceptibility to TAT-RasGAP_317-326_.

In summary, the results presented in this article bring a better understanding of the factors that influence the antimicrobial activity of TAT-RasGAP_317-326_. We describe the importance of bacterial envelope integrity on the sensitivity towards TAT-RasGAP_317-326_. Factors such as divalent salts, EDTA and LPS structure influence the concentration of peptide needed to inhibit bacterial growth. Furthermore, we report the effect of TAT-RasGAP_317-326_ on the transcriptional landscape of *E. coli* and highlight the importance of a broad range of two-component systems in the adaptation of bacteria towards this AMP. We finally investigated the effect of other AMPs on the activity of TAT-RasGAP_317-326_ and could select TAT-RasGAP_317-326_-resistant bacteria. Our observation that sensitivity could be increased and resistance could be reversed by addition of EDTA is important in the perspective of a clinical use of this peptide to improve its efficiency and to prevent rapid emergence of resistance.

### Limitations of the study

Results presented in this study originate from *in vitro* studies. They might thus only be partially representative of which interactions would happen in an *in vivo* model of infection. Indeed, several factors such as presence of endogenous AMPs, as well as proteins or other components with which TAT-RasGAP_317-326_ may interact are not present in our system. Moreover, interactions between TAT-RasGAP_317-326_ and other AMPs need to be investigated in further details using checkerboard assays, in order to determine putative synergisms. Similarly, mechanisms of action of the peptide and mechanisms of resistance towards the peptide that were selected need to be further investigated in the future, in order to describe how TAT-RasGAP_317-326_ interacts with bacteria at the molecular level.

## Acknowledgments

We would like to thank Sébastien Aeby and Yasmina Merzouk for technical support, Valentin Scherz for support in bioinformatics analyses and Prof. Gilbert Greub for sharing equipment and laboratories. This study was supported by an interdisciplinary grant of the Faculty of Biology and Medicine of the University of Lausanne.

## Author contributions

MG, TH, NJ, AV, SH and SC performed experiments. MG, TH, LE, CW and NJ were involved in the planning of the project and discussed the results. MG, TH, NJ and TP analysed the results. MG, CW and NJ wrote the manuscript. All the authors proofread the manuscript.

## Declaration of interest

The authors declare no competing interests.

## Material and methods

### Strains, growth conditions and chemicals

*E. coli* strains K-12 MG1655, ATCC 25922 and BW25113 were grown in LB or Basal Medium 2 (BM2; 62 mM potassium phosphate buffer [pH 7.0], 7 mM (NH_4_)_2_SO_4_, 10 μM FeSO_4_, 0.4% (wt/v) glucose and 0.5% tryptone) with high (2 mM) or low (20 μM) concentration of magnesium (MgSO_4_) (Fernandez et al., 2012). *Pseudomonas aeruginosa* strain PA14 was grown either in LB or BM2 medium. *Staphylococcus capitis* (Heulot et al., 2017) and *S. aureus* (ATCC 29213) strains were grown in tryptic soy broth (TSB) (Missiakas and Schneewind, 2013). All strains were stored at −80°C, in their respective medium, supplemented with ~25% glycerol. When required, antibiotics were added at final concentrations of 50 μg/mL (kanamycin), 20 μg/mL (gentamycin), or 100 μg/mL (carbenicillin). The retro-inverse TAT-RasGAP_317-326_ peptide (amino acid sequence DTRLNTVWMWGGRRRQRRKKRG) and the N-terminal FITC-labelled version of this peptide were synthesized by SBS Genetech (Beijing, China) and stored at −20°C. Chemicals were purchased from Sigma-Aldrich (St-Louis, MO, USA), unless otherwise specified.

### MIC measurements

The minimum inhibitory concentration (MIC) of peptide was defined as the lowest concentration of peptide that resulted in no visible growth. Overnight cultures were diluted to OD_600_ = 0.1 and grown with shaking at 37°C for 1 hour. MICs were measured by diluting these cultures (1:20 for LB and TSB cultures and 1:8 for BM2 cultures) and then adding these dilutions to 2-fold sequential dilutions of the peptides in 96-well plates. Volume of media (with peptide) per well was 100 μl and 10 μl of diluted cultures were added to each well. Cell growth was monitored via OD_590_ measurement after overnight growth with shaking at 37°C. OD_590_ readings were measured by FLUOstar Omega microplate reader (BMG Labtech, Ortenberg, Germany). Peptide-free growth control wells and bacteria-free contamination control wells were included. First concentration at which no bacterial growth could be detected was defined as the MIC.

For MIC measurements in presence of *E. coli* LPS or EDTA, the indicated concentrations of these substances were dissolved in LB and distributed in 96-well plates prior to addition of the peptides. OD_590_ measurements of control wells without peptides were used to calculate the percentage of growth in presence of EDTA.

### Growth curves

Overnight cultures were diluted to OD_600_ = 0.1 and grown with shaking at 37°C for 1 hour, before addition of peptide. Cell growth was monitored via OD_600_ measurement by Novaspec II Visible spectrophotometer (Pharmacia LKB Biotechnology, Cambridge, England) at 2, 4 and 6 hours.

Combinations of antimicrobial peptides were tested using the above methods and combining “half” concentrations of the different peptides (2 μM TAT-RasGAP_317-326_, 64 μg/ml melittin, 32 μg/ml LL-37 or 1 μg/ml polymyxin B) to produce supplementary Figure 6C.

### CFU measurements

Overnight cultures were diluted to OD_600_ = 0.1 and grown with shaking at 37°C for 1 hour, before addition of the peptide. Each time point was taken by removing 10 μl and performing 10-fold serial dilutions. Dilutions of each condition were then plated in the absence of peptide and grown at 37°C overnight. CFU were measured by counting the number of colonies on the plates after overnight incubation.

### Confocal microscopy

Overnight cultures of *E. coli* MG1655 were diluted to OD_600_ = 0.1, grown for 1 hour, incubated for 1 hour with 10 μM FITC-labelled TAT-RasGAP_317-326_, stained with 5 μg/ml FM4-64 and fixed with 4% paraformaldehyde solution. Incubation with DAPI was subsequently performed and pictures were acquired on a LSM710 confocal microscope (Zeiss, Oberkochen, Germany). Images were analyzed with ImageJ software (Schneider et al., 2012).

### Electron microscopy

Bacteria were fixed with 2.5% glutaraldehyde solution (EMS, Hatfield, PA) in Phosphate Buffer (PB 0.1 M pH 7.4) for 1 hour at room temperature. Then, bacterial samples were incubated in a freshly prepared mix of 1% osmium tetroxide (EMS) and 1.5% potassium ferrocyanide in phosphate buffer for 1 hour at room temperature. The samples were then washed three times in distilled water and spun down in 2% low melting agarose, solidified on ice, cut into 1 mm^3^ cubes and dehydrated in acetone solution at graded concentrations (30% for 40 minutes; 50% for 40 minutes; 70% for 40 minutes and 100% for 3 times 1 hour). This was followed by infiltration in Epon at graded concentrations (Epon 1/3 acetone for 2 hours; Epon 3/1 acetone for 2 hours, Epon 1/1 for 4 hours and Epon 1/1 for 12 hours) and finally polymerization for 48 hours at 60°C in a laboratory oven. Ultrathin sections of 50 nm were cut on a Leica Ultramicrotome (Leica Mikrosysteme GmbH, Vienna, Austria) and placed on a copper slot grid 2×1mm (EMS) coated with a polystyrene film. The bacterial sections were stained in 4% uranyl acetate for 10 minutes, rinsed several times with water, then incubated in Reynolds lead citrate and finally rinsed several times with water before imaging.

Micrographs (10×10 tiles) with a pixel size of 1.209 nm over an area of 40×40 μm were taken with a transmission electron microscope Philips CM100 (Thermo Fisher Scientific, Waltham, MA) at an acceleration voltage of 80kV with a TVIPS TemCam-F416 digital camera (TVIPS GmbH, Gauting, Germany). Large montage alignments were performed using Blendmont command-line program from the IMOD software (Kremer et al., 1996) and treated with ImageJ software.

### Flow cytometry

Overnight cultures of *E. coli* MG1655 were diluted 1:100 and grown to mid exponential phase (OD_600_ = 0.4-0.6) with shaking at 37°C. Each culture was then diluted to OD_600_ = 0.1, grown with shaking at 37°C for 1 hour and then treated with 10 μM FITC-labelled peptide for 1 hour. Following peptide treatment, bacterial cells were washed in PBS and diluted 1:5 before acquisition on a CytoFLEX benchtop flow cytometer (Beckman Coulter). For each sample, 10,000 events were collected and analyzed. Extracellular fluorescence was quenched with 0.2% Trypan Blue (TB). TB is an efficient quencher of extracellular fluorescence (Sahlin et al., 1983, Loike and Silverstein, 1983, Jevprasesphant et al., 2004, Wan et al., 1993) and allows quantification of fluorescent signal from intracellular peptide (not subject to quenching by TB). P values were calculated using ratio paired t-test.

### RNA-Seq

Overnight cultures of *E. coli* MG1655 were diluted to OD_600_ = 0.1 and grown with shaking at 37°C for one hour to mid exponential phase (OD_600_ = 0.4-0.6). Cultures were then treated with TAT-RasGAP_317-326_ (10 μM) or left untreated (negative control), and grown with shaking at 37°C for an additional hour. For RNA extraction, protocol 1 in the RNAprotect Bacteria Reagent Handbook (Enzymatic lysis of bacteria) was followed using the RNeasy Plus Mini Kit (Qiagen) using TE buffer (10 mM Tris-HCl, 1 mM EDTA, pH 8.0) containing 1 mg/ml lysozyme (AppliChem, Chicago, IL). In the last step, RNA was eluted in 30 μl RNase-free water. Next, any contaminating DNA was removed using the DNA-free™ DNA Removal Kit (Invitrogen, Carlsbad, CA). 10x DNase buffer was added to the 30 μl eluted RNA with 2 μl rDNase I. This mix was incubated for 30 minutes at 37°C followed by rDNase I inactivation with 7 μl DNase Inactivation Reagent for 2 minutes with shaking (700 rpm) at room temperature. Samples were then centrifuged for 90 seconds at 10,000 x *g*, supernatant was transferred to a new tube, and stored at −80°C. Integrity of the samples was verified using the Standard Sensitivity RNA Analysis kit (Advanced Analytical, Ankeny, IA) with the Fragment Analyser Automated CE System (Labgene Scientific, Châtel-Saint-Denis, Switzerland). Samples that met RNA-Seq requirements were further processed and sent for sequencing. Preparation of the libraries and Illumina HiSeq platform (1×50 bp) sequencing were performed by Fasteris (Plan-les-Ouates, Switzerland). Raw reads were trimmed with trimmomatic version 0.36 (Bolger et al., 2014) (parameters: ILLUMINACLIP: NexteraPE-PE.fa:3:25:6, LEADING: 28, TRAILING: 28 MINLEN: 30). Trimmed reads were mapped to the genome of *E. coli* K-12 MG1655 (accession: NC_000913.3) with bwa mem version 0.7.17 (https://arxiv.org/abs/1303.3997) using default parameters. Htseq version 0.11.2 (Anders et al., 2015) was used to count reads aligned to each gene (parameters: --stranded=no -t gene). Normalized expression values were calculated as Reads Per Kilobase of transcript per Million mapped reads (RPKM) with edgeR (Robinson et al., 2010).

### Keio collection screening

Deletion mutants from the Keio collection (Baba et al., 2006, Yamamoto et al., 2009) were used, along with the corresponding wild-type, which was added as a control on each test plate. Overnight cultures were diluted 1:100 in LB medium. Bacteria were incubated at 37°C for 1 hour before adding TAT-RasGAP_317-326_ (5 μM final concentration). Plates were incubated statically at 37°C and OD_590_ was measured at 0 hour, 1.5 hour, 3 hours, 6 hours and 24 hours with FLUOstar Omega plate reader. Measurements were combined and analysed with R (version 3.6.1, (Team, 2019)). Data analysis and visualisation were performed with the *dplyr* (version 0.8.5) and *ggplot2* (version 3.3.0) packages from the *tidyverse* (version 1.3.0) environment. Since starting OD_590_ (OD in equations) varied between strains and conditions, the OD_590_ starting values in each well was subtracted from corresponding measurements made at time t in the presence (P) or absence (noP) of TAT-RasGAP_317-326_. For each strain, NG^m_t_^(P), the normalized growth value for a mutant strain at time t in the presence of the peptide was calculated with the following formula:

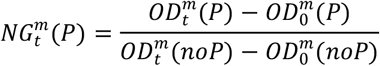

Normalized growths of wild-type strain (mean WT), as presented on Fig. 4a and b were calculated by averaging normalized growths of all the wild-type controls performed (N=270). To normalize the growth of a mutant (m) to the growth of control (c) bacteria (wild-type) on the same plate, the NG^m^_t_(noP) factor was calculated with the following formula:

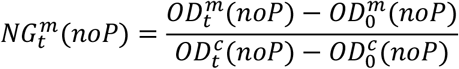

Gene ontology (GO) annotation (The Gene Ontology, 2019) was obtained from GO database (2020-09-01, “http://current.geneontology.org/annotations”) and assigned to the list of gene deletion inducing hypersensitivity with the GO.db package (version 3.10.0 (Carslon, 2019)). GO IDs were assigned to each gene and the corresponding GO names were obtained with the “Term” function. Additionally, the same set of genes was subjected to KEGG pathways analysis (Kanehisa and Goto, 2000) with the KEGGREST package (version 1.26.1). Briefly, the KEGG orthology (KO) and KEGG pathway annotation were obtained from the KEGG database (Kanehisa, 2019) for *E. coli* K-12 MG1655 (eco). The code is available on Github (https://github.com/njacquie/TAT-RasGAP_project).

### *Pseudomonas aeruginosa* PA14 transposon library screening

The library of transposon (Tn) mutants in *P. aeruginosa* PA14 (Vitale et al., 2020) was grown in BM2 supplemented with 20 μM MgSO_4_ (Fernandez et al., 2012) and 0.2% L-rhamnose monohydrate (Sigma-Aldrich) in the absence or presence of 0.5 μM TAT-RasGAP_317-326_. Following growth for 12 generations, genomic DNA (gDNA) was extracted with the GenElute Bacterial Genomic DNA Kit (Sigma-Aldrich). The transposon sequencing (Tn-seq) circle method (Gallagher et al., 2011, Gallagher et al., 2013) was employed to sequence the transposon junctions. Briefly, the gDNA was sheared to an average size of 300 bp fragments with a focused-ultrasonicator. The DNA fragments were repaired and ligated to adapters with the NEBNext Ultra II DNA Library Prep Kit for Illumina (New England Biolabs). Following restriction of the Tn with BamHI (New England Biolabs), the fragments were circularized by ligation and exonuclease treatment was applied to remove undesired non-circularized DNAs (Gallagher et al., 2011). The Tn junctions were PCR amplified and amplicons were sequenced with the MiSeq Reagent Kit v2, 300-cycles (Illumina).

Following sequencing, the adapter sequences of the reads (.fastq) were trimmed with the command line “cutadapt -a adapter -q quality -o output.fastq.gz input.fastq.gz” (Martin, 2011). The software Tn-Seq Explorer (Solaimanpour et al., 2015) mapped the trimmed and paired reads onto the *P. aeruginosa* UCBPP-PA14 genome (Winsor et al., 2016), and determined the unique insertion density (UID, i.e. the number of unique Tn insertions divided per the length of the gene). The normalized UID between the treated and non-treated samples were compared and this ratio (log2-fold change, FC) was used to identify resistant determinants (log2-FC < − 1.0 and normalized UID > 0.0045).

### Selection of resistant mutants

Bacteria were grown in the corresponding medium, diluted 1:100 and cultured overnight with 0.5x MIC of TAT-RasGAP_317-326_. The subculture was diluted 1:100 and incubated with 0.5x or 1x MIC overnight. Cells that successfully grew were diluted 1:100 in medium containing the same concentration or twice the concentration of peptide. Each dilution in fresh medium containing peptide is considered one passage. This process was repeated for up to 20 passages.

## Notes

### Competing Interest Statement

The authors have declared no competing interest.

### Summary of Updates

This new version of the manuscript contains new versions of figures which were improved with new experiments.

## References

Akbari, R., Hakemi-Vala, M., Pashaie, F., Bevalian, P., Hashemi, A. & Pooshang Bagheri, K. 2019. Highly Synergistic Effects of Melittin with Conventional Antibiotics Against Multidrug-Resistant Isolates of Acinetobacter baumannii and Pseudomonas aeruginosa. Microb Drug Resist, 25, 193–202.

Anders, S., Pyl, P. T. & Huber, W. 2015. HTSeq--a Python framework to work with high-throughput sequencing data. Bioinformatics, 31, 166–9.

Annibaldi, A., Heulot, M., Martinou, J. C. & Widmann, C. 2014. TAT-RasGAP_317-326_-mediated tumor cell death sensitization can occur independently of Bax and Bak. Apoptosis, 19, 719–33.

Ashburner, M., Ball, C. A., Blake, J. A., Botstein, D., Butler, H., Cherry, J. M., Davis, A. P., Dolinski, K., Dwight, S. S., Eppig, J. T., Harris, M. A., Hill, D. P., Issel-Tarver, L., Kasarskis, A., Lewis, S., Matese, J. C., Richardson, J. E., Ringwald, M., Rubin, G. M. & Sherlock, G. 2000. Gene ontology: tool for the unification of biology. The Gene Ontology Consortium. Nat Genet, 25, 25–9.

Baba, T., Ara, T., Hasegawa, M., Takai, Y., Okumura, Y., Baba, M., Datsenko, K. A., Tomita, M., Wanner, B. L. & Mori, H. 2006. Construction of Escherichia coli K-12 in-frame, single-gene knockout mutants: the Keio collection. Mol Syst Biol, 2, 2006 0008.

Bader, M. W., Sanowar, S., Daley, M. E., Schneider, A. R., Cho, U., Xu, W., Klevit, R. E., Le Moual, H. & Miller, S. I. 2005. Recognition of antimicrobial peptides by a bacterial sensor kinase. Cell, 122, 461–72.

Baranova, N. & Nikaido, H. 2002. The baeSR two-component regulatory system activates transcription of the yegMNOB (mdtABCD) transporter gene cluster in Escherichia coli and increases its resistance to novobiocin and deoxycholate. J Bacteriol, 184, 4168–76.

Barras, D., Chevalier, N., Zoete, V., Dempsey, R., Lapouge, K., Olayioye, M. A., Michielin, O. & Widmann, C. 2014a. A WXW motif is required for the anticancer activity of the TAT-RasGAP_317-326_ peptide. J Biol Chem, 289, 23701–11.

Barras, D., Lorusso, G., Lhermitte, B., Viertl, D., Ruegg, C. & Widmann, C. 2014b. Fragment N2, a caspase-3-generated RasGAP fragment, inhibits breast cancer metastatic progression. Int J Cancer, 135, 242–7.

Barras, D., Lorusso, G., Ruegg, C. & Widmann, C. 2014c. Inhibition of cell migration and invasion mediated by the TAT-RasGAP_317-326_ peptide requires the DLC1 tumor suppressor. Oncogene, 33, 5163–72.

Bociek, K., Ferluga, S., Mardirossian, M., Benincasa, M., Tossi, A., Gennaro, R. & Scocchi, M. 2015. Lipopolysaccharide Phosphorylation by the WaaY Kinase Affects the Susceptibility of Escherichia coli to the Human Antimicrobial Peptide LL-37. J Biol Chem, 290, 19933–41.

Bolger, A. M., Lohse, M. & Usadel, B. 2014. Trimmomatic: a flexible trimmer for Illumina sequence data. Bioinformatics, 30, 2114–20.

Brogden, K. A. 2005. Antimicrobial peptides: pore formers or metabolic inhibitors in bacteria? Nat Rev Microbiol, 3, 238–50.

Cariss, S. J., Constantinidou, C., Patel, M. D., Takebayashi, Y., Hobman, J. L., Penn, C. W. & Avison, M. B. 2010. YieJ (CbrC) mediates CreBC-dependent colicin E2 tolerance in Escherichia coli. J Bacteriol, 192, 3329–36.

Carslon, M. 2019. GO.db: A set of annotation maps describing the entire Gene Ontology.

Clifton, L. A., Skoda, M. W., Le Brun, A. P., Ciesielski, F., Kuzmenko, I., Holt, S. A. & Lakey, J. H. 2015. Effect of divalent cation removal on the structure of gram-negative bacterial outer membrane models. Langmuir, 31, 404–12.

Deutscher, J., Francke, C. & Postma, P. W. 2006. How phosphotransferase system-related protein phosphorylation regulates carbohydrate metabolism in bacteria. Microbiol Mol Biol Rev, 70, 939–1031.

Di Somma, A., Moretta, A., Cane, C., Cirillo, A. & Duilio, A. 2020. Antimicrobial and Antibiofilm Peptides. Biomolecules, 10.

Domingues, M. M., Inacio, R. G., Raimundo, J. M., Martins, M., Castanho, M. A. & Santos, N. C. 2012. Biophysical characterization of polymyxin B interaction with LPS aggregates and membrane model systems. Biopolymers, 98, 338–44.

Eder, K., Vizler, C., Kusz, E., Karcagi, I., Glavinas, H., Balogh, G. E., Vigh, L., Duda, E. & Gyorfy, Z. 2009. The role of lipopolysaccharide moieties in macrophage response to Escherichia coli. Biochem Biophys Res Commun, 389, 46–51.

Fernandez, L., Gooderham, W. J., Bains, M., Mcphee, J. B., Wiegand, I. & Hancock, R. E. 2010. Adaptive resistance to the "last hope" antibiotics polymyxin B and colistin in Pseudomonas aeruginosa is mediated by the novel two-component regulatory system ParR-ParS. Antimicrob Agents Chemother, 54, 3372–82.

Fernandez, L., Jenssen, H., Bains, M., Wiegand, I., Gooderham, W. J. & Hancock, R. E. 2012. The two-component system CprRS senses cationic peptides and triggers adaptive resistance in Pseudomonas aeruginosa independently of ParRS. Antimicrob Agents Chemother, 56, 6212–22.

Franklin, M. J., Nivens, D. E., Weadge, J. T. & Howell, P. L. 2011. Biosynthesis of the Pseudomonas aeruginosa Extracellular Polysaccharides, Alginate, Pel, and Psl. Front Microbiol, 2, 167.

Gallagher, L. A., Ramage, E., Patrapuvich, R., Weiss, E., Brittnacher, M. & Manoil, C. 2013. Sequence-defined transposon mutant library of Burkholderia thailandensis. mBio, 4, e00604–13.

Gallagher, L. A., Shendure, J. & Manoil, C. 2011. Genome-scale identification of resistance functions in Pseudomonas aeruginosa using Tn-seq. mBio, 2, e00315–10.

Gennaro, R., Skerlavaj, B. & Romeo, D. 1989. Purification, composition, and activity of two bactenecins, antibacterial peptides of bovine neutrophils. Infect Immun, 57, 3142–6.

Hancock, R. E. 1997. The bacterial outer membrane as a drug barrier. Trends Microbiol, 5, 37–42.

Hancock, R. E. W. 1984. Alterations in Outer-Membrane Permeability. Annual Review of Microbiology, 38, 237–264.

Hassan, M., Kjos, M., Nes, I. F., Diep, D. B. & Lotfipour, F. 2012. Natural antimicrobial peptides from bacteria: characteristics and potential applications to fight against antibiotic resistance. J Appl Microbiol, 113, 723–36.

Heinonen, T., Hargraves, S., Georgieva, M., Widmann, C. & Jacquier, N. 2021. The antimicrobial peptide TAT-RasGAP_317-326_ inhibits the formation and the expansion of bacterial biofilms in vitro. J Glob Antimicrob Resist.

Heulot, M., Chevalier, N., Puyal, J., Margue, C., Michel, S., Kreis, S., Kulms, D., Barras, D., Nahimana, A. & Widmann, C. 2016. The TAT-RasGAP_317-326_ anti-cancer peptide can kill in a caspase-, apoptosis-, and necroptosis-independent manner. Oncotarget, 7, 64342–64359.

Heulot, M., Jacquier, N., Aeby, S., Le Roy, D., Roger, T., Trofimenko, E., Barras, D., Greub, G. & Widmann, C. 2017. The Anticancer Peptide TAT-RasGAP_317-326_ Exerts Broad Antimicrobial Activity. Front Microbiol, 8, 994.

Hong, J., Lu, X., Deng, Z., Xiao, S., Yuan, B. & Yang, K. 2019. How Melittin Inserts into Cell Membrane: Conformational Changes, Inter-Peptide Cooperation, and Disturbance on the Membrane. Molecules, 24.

Jevprasesphant, R., Penny, J., Attwood, D. & D’Emanuele, A. 2004. Transport of dendrimer nanocarriers through epithelial cells via the transcellular route. J Control Release, 97, 259–67.

Kanehisa, M. 2019. Toward understanding the origin and evolution of cellular organisms. Protein Sci, 28, 1947–1951.

Kanehisa, M. & Goto, S. 2000. KEGG: kyoto encyclopedia of genes and genomes. Nucleic Acids Res, 28, 27–30.

Kanehisa, M., Sato, Y., Furumichi, M., Morishima, K. & Tanabe, M. 2019. New approach for understanding genome variations in KEGG. Nucleic Acids Res, 47, D590–D595.

Kong, X. D., Moriya, J., Carle, V., Pojer, F., Abriata, L. A., Deyle, K. & Heinis, C. 2020. De novo development of proteolytically resistant therapeutic peptides for oral administration. Nat Biomed Eng, 4, 560–571.

Kremer, J. R., Mastronarde, D. N. & Mcintosh, J. R. 1996. Computer visualization of three-dimensional image data using IMOD. J Struct Biol, 116, 71–6.

Kumar, P., Kizhakkedathu, J. N. & Straus, S. K. 2018. Antimicrobial Peptides: Diversity, Mechanism of Action and Strategies to Improve the Activity and Biocompatibility In Vivo. Biomolecules, 8.

Laubacher, M. E. & Ades, S. E. 2008. The Rcs phosphorelay is a cell envelope stress response activated by peptidoglycan stress and contributes to intrinsic antibiotic resistance. J Bacteriol, 190, 2065–74.

Lazar, V., Martins, A., Spohn, R., Daruka, L., Grezal, G., Fekete, G., Szamel, M., Jangir, P. K., Kintses, B., Csorgo, B., Nyerges, A., Gyorkei, A., Kincses, A., Der, A., Walter, F. R., Deli, M. A., Urban, E., Hegedus, Z., Olajos, G., Mehi, O., Balint, B., Nagy, I., Martinek, T. A., Papp, B. & Pal, C. 2018. Antibiotic-resistant bacteria show widespread collateral sensitivity to antimicrobial peptides. Nat Microbiol, 3, 718–731.

Lazzaro, B. P., Zasloff, M. & Rolff, J. 2020. Antimicrobial peptides: Application informed by evolution. Science, 368.

Lee, M. T., Sun, T. L., Hung, W. C. & Huang, H. W. 2013. Process of inducing pores in membranes by melittin. Proc Natl Acad Sci U S A, 110, 14243–8.

Leive, L. 1965. Release of lipopolysaccharide by EDTA treatment of E. coli. Biochem Biophys Res Commun, 21, 290–6.

Leon-Buitimea, A., Garza-Cardenas, C. R., Garza-Cervantes, J. A., Lerma-Escalera, J. A. & Morones-Ramirez, J. R. 2020. The Demand for New Antibiotics: Antimicrobial Peptides, Nanoparticles, and Combinatorial Therapies as Future Strategies in Antibacterial Agent Design. Front Microbiol, 11, 1669.

Loike, J. D. & Silverstein, S. C. 1983. A fluorescence quenching technique using trypan blue to differentiate between attached and ingested glutaraldehyde-fixed red blood cells in phagocytosing murine macrophages. J Immunol Methods, 57, 373–9.

Macfarlane, E. L., Kwasnicka, A., Ochs, M. M. & Hancock, R. E. 1999. PhoP-PhoQ homologues in Pseudomonas aeruginosa regulate expression of the outer-membrane protein OprH and polymyxin B resistance. Mol Microbiol, 34, 305–16.

Martin, M. 2011. Cutadapt removes adapter sequences from high-throughput sequencing reads. EMBnet.journal, 17, 10–12.

Mcphee, J. B., Lewenza, S. & Hancock, R. E. 2003. Cationic antimicrobial peptides activate a two-component regulatory system, PmrA-PmrB, that regulates resistance to polymyxin B and cationic antimicrobial peptides in Pseudomonas aeruginosa. Mol Microbiol, 50, 205–17.

Mendez-Samperio, P. 2010. The human cathelicidin hCAP18/LL-37: a multifunctional peptide involved in mycobacterial infections. Peptides, 31, 1791–8.

Michod, D., Annibaldi, A., Schaefer, S., Dapples, C., Rochat, B. & Widmann, C. 2009. Effect of RasGAP N2 fragment-derived peptide on tumor growth in mice. J Natl Cancer Inst, 101, 828–32.

Michod, D., Yang, J. Y., Chen, J., Bonny, C. & Widmann, C. 2004. A RasGAP-derived cell permeable peptide potently enhances genotoxin-induced cytotoxicity in tumor cells. Oncogene, 23, 8971–8.

Missiakas, D. M. & Schneewind, O. 2013. Growth and laboratory maintenance of Staphylococcus aureus. Curr Protoc Microbiol, Chapter 9, Unit 9C 1.

O’Neill, J. 2016. Tackling Drug-resistant Infections Globally: Final Report and Recommendations of the Review on Antimicrobial Resistance. In: Government, H. (ed.). London.

Ogasawara, H., Hasegawa, A., Kanda, E., Miki, T., Yamamoto, K. & Ishihama, A. 2007. Genomic SELEX search for target promoters under the control of the PhoQP-RstBA signal relay cascade. J Bacteriol, 189, 4791–9.

Olaitan, A. O., Morand, S. & Rolain, J. M. 2014. Mechanisms of polymyxin resistance: acquired and intrinsic resistance in bacteria. Front Microbiol, 5, 643.

Pelletier, C., Bourlioux, P. & Van Heijenoort, J. 1994. Effects of sub-minimal inhibitory concentrations of EDTA on growth of Escherichia coli and the release of lipopolysaccharide. FEMS Microbiol Lett, 117, 203–6.

Robinson, M. D., Mccarthy, D. J. & Smyth, G. K. 2010. edgeR: a Bioconductor package for differential expression analysis of digital gene expression data. Bioinformatics, 26, 139–40.

Sahlin, S., Hed, J. & Rundquist, I. 1983. Differentiation between attached and ingested immune complexes by a fluorescence quenching cytofluorometric assay. J Immunol Methods, 60, 115–24.

Schneider, C. A., Rasband, W. S. & Eliceiri, K. W. 2012. NIH Image to ImageJ: 25 years of image analysis. Nat Methods, 9, 671–5.

Serulla, M., Ichim, G., Stojceski, F., Grasso, G., Afonin, S., Heulot, M., Schober, T., Roth, R., Godefroy, C., Milhiet, P. E., Das, K., Garcia-Saez, A. J., Danani, A. & Widmann, C. 2020. TAT-RasGAP_317-326_ kills cells by targeting inner-leaflet-enriched phospholipids. Proc Natl Acad Sci U S A.

Solaimanpour, S., Sarmiento, F. & Mrazek, J. 2015. Tn-seq explorer: a tool for analysis of high-throughput sequencing data of transposon mutant libraries. PLoS One, 10, e0126070.

Spohn, R., Daruka, L., Lazar, V., Martins, A., Vidovics, F., Grezal, G., Mehi, O., Kintses, B., Szamel, M., Jangir, P. K., Csorgo, B., Gyorkei, A., Bodi, Z., Farago, A., Bodai, L., Foldesi, I., Kata, D., Maroti, G., Pap, B., Wirth, R., Papp, B. & Pal, C. 2019. Integrated evolutionary analysis reveals antimicrobial peptides with limited resistance. Nat Commun, 10, 4538.

Srinivas, P. & Rivard, K. 2017. Polymyxin Resistance in Gram-negative Pathogens. Curr Infect Dis Rep, 19, 38.

Team, R. C. 2019. R: A language and environment for statistical computing.

The Gene Ontology, C. 2019. The Gene Ontology Resource: 20 years and still GOing strong. Nucleic Acids Res, 47, D330–D338.

Tsoutsou, P., Annibaldi, A., Viertl, D., Ollivier, J., Buchegger, F., Vozenin, M. C., Bourhis, J., Widmann, C. & Matzinger, O. 2017. TAT-RasGAP_317-326_ Enhances Radiosensitivity of Human Carcinoma Cell Lines In Vitro and In Vivo through Promotion of Delayed Mitotic Cell Death. Radiat Res, 187, 562–569.

Vitale, A., Pessi, G., Urfer, M., Locher, H. H., Zerbe, K., Obrecht, D., Robinson, J. A. & Eberl, L. 2020. Identification of Genes Required for Resistance to Peptidomimetic Antibiotics by Transposon Sequencing. Front Microbiol, 11, 1681.

Wan, C. P., Park, C. S. & Lau, B. H. 1993. A rapid and simple microfluorometric phagocytosis assay. J Immunol Methods, 162, 1–7.

Wang, G., Li, X. & Wang, Z. 2016. APD3: the antimicrobial peptide database as a tool for research and education. Nucleic Acids Res, 44, D1087–93.

Weatherspoon-Griffin, N., Yang, D., Kong, W., Hua, Z. & Shi, Y. 2014. The CpxR/CpxA two-component regulatory system up-regulates the multidrug resistance cascade to facilitate Escherichia coli resistance to a model antimicrobial peptide. J Biol Chem, 289, 32571–82.

Winsor, G. L., Griffiths, E. J., Lo, R., Dhillon, B. K., Shay, J. A. & Brinkman, F. S. 2016. Enhanced annotations and features for comparing thousands of Pseudomonas genomes in the Pseudomonas genome database. Nucleic Acids Res, 44, D646–53.

Xhindoli, D., Pacor, S., Benincasa, M., Scocchi, M., Gennaro, R. & Tossi, A. 2016. The human cathelicidin LL-37--A pore-forming antibacterial peptide and host-cell modulator. Biochim Biophys Acta, 1858, 546–66.

Yadavalli, S. S., Carey, J. N., Leibman, R. S., Chen, A. I., Stern, A. M., Roggiani, M., Lippa, A. M. & Goulian, M. 2016. Antimicrobial peptides trigger a division block in Escherichia coli through stimulation of a signalling system. Nat Commun, 7, 12340.

Yamamoto, N., Nakahigashi, K., Nakamichi, T., Yoshino, M., Takai, Y., Touda, Y., Furubayashi, A., Kinjyo, S., Dose, H., Hasegawa, M., Datsenko, K. A., Nakayashiki, T., Tomita, M., Wanner, B. L. & Mori, H. 2009. Update on the Keio collection of Escherichia coli single-gene deletion mutants. Mol Syst Biol, 5, 335.

Yethon, J. A., Heinrichs, D. E., Monteiro, M. A., Perry, M. B. & Whitfield, C. 1998. Involvement of waaY, waaQ, and waaP in the modification of Escherichia coli lipopolysaccharide and their role in the formation of a stable outer membrane. J Biol Chem, 273, 26310–6.

