## Supplementary figures and tables for "Bacterial surface properties influence the activity of the TAT-RasGAP_317-326_ antimicrobial peptide"

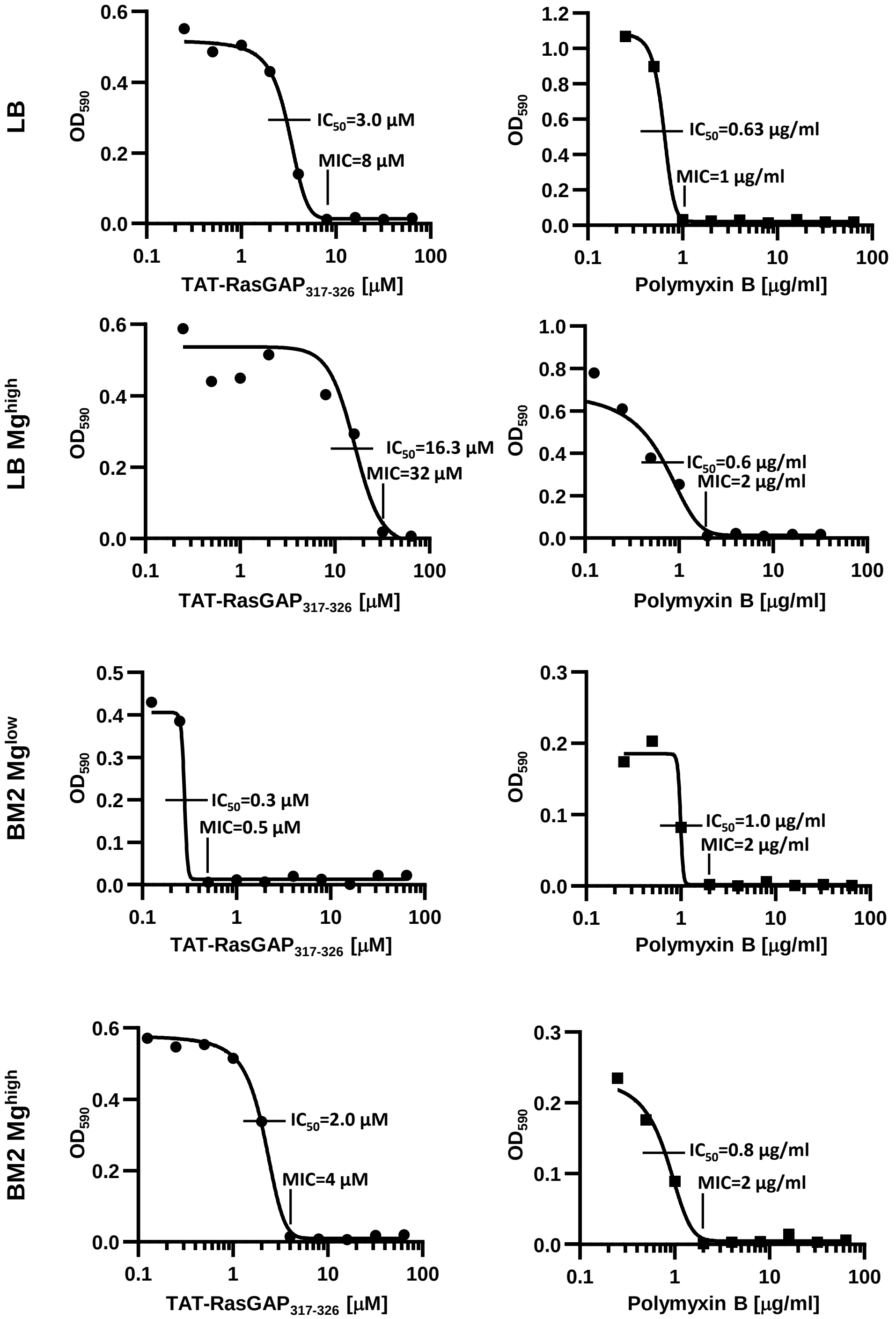

**Supplementary Figure 1. *E. coli* MG1655 sensitivity to TAT-RasGAP$_{317\text{-}326}$ and polymyxin B varies depending on the growth medium used.** *E. coli* MG1655 were grown overnight in the indicated medium, diluted to $OD_{600}$ = 0.1 and grown for an additional 1-hour period. Culture was then diluted 20 times and 10 µl was added per well of a 96-well plate containing serial dilutions of TAT-RasGAP$_{317\text{-}326}$ or polymyxin B. $OD_{590}$ measurements after 16 hours of incubation in the presence of the indicated concentrations of TAT-RasGAP$_{317\text{-}326}$ are shown and MIC is defined as the lowest concentration of TAT-RasGAP$_{317\text{-}326}$ that completely inhibits bacterial proliferation. $IC_{50}$ is defined as the concentration required to inhibit 50% of growth and was calculated using GraphPad Prism 8.

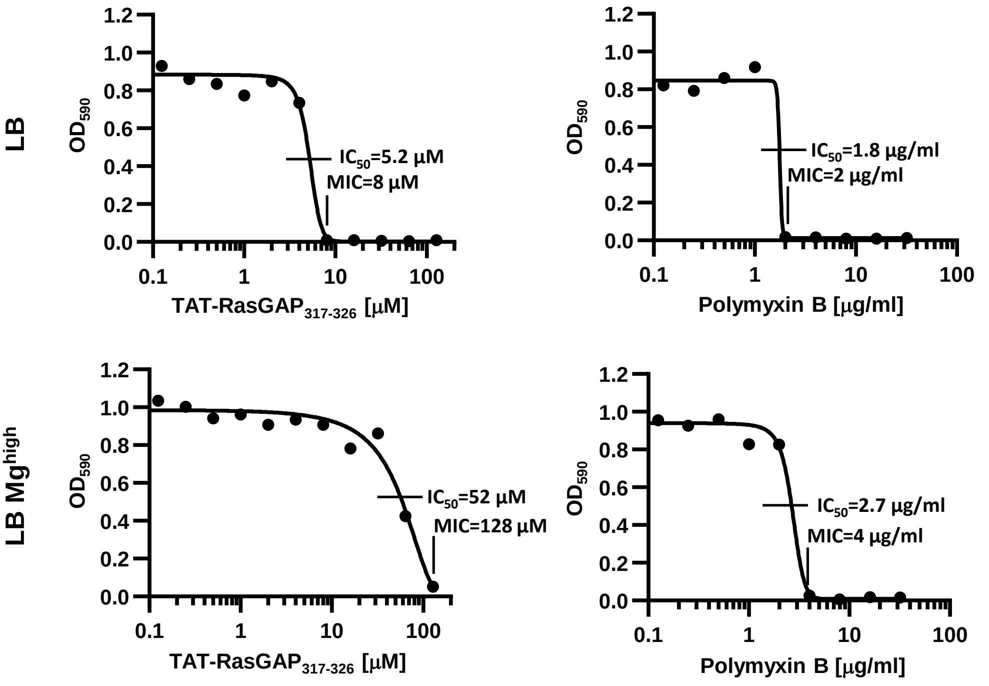

**Supplementary Figure 2. *E. coli* ATCC 25922 sensitivity to TAT-RasGAP$_{317-326}$ and polymyxin B varies depending on the growth medium used.** *E. coli* ATCC 25922 were treated and analyzed as indicated in Supplementary Figure 1.

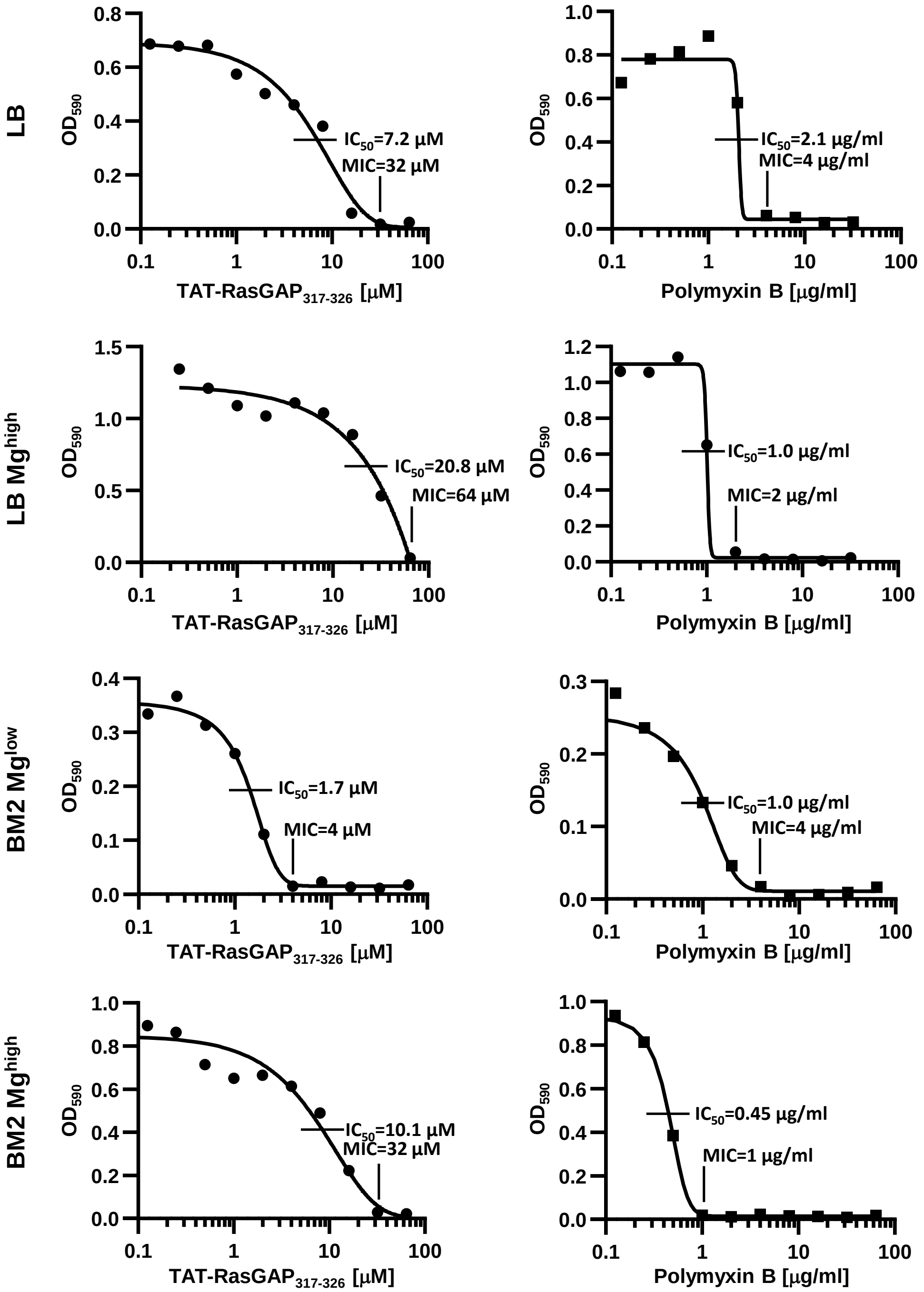

**Supplementary Figure 3. *P. aeruginosa* PA14 sensitivity to TAT-RasGAP$_{317-326}$ and polymyxin B varies depending on the growth medium used.** *P. aeruginosa* PA14 were grown overnight in the indicated medium, diluted to $OD_{600}$ = 0.1 and grown for 1 hour. Culture was then diluted 10 times and 10 µl was added per well of a 96-well plate containing serial dilutions of TAT-RasGAP$_{317-326}$ or polymyxin B. $OD_{590}$ measurements, MIC and IC50 calculations were performed similarly as in Supplementary Figure 1.

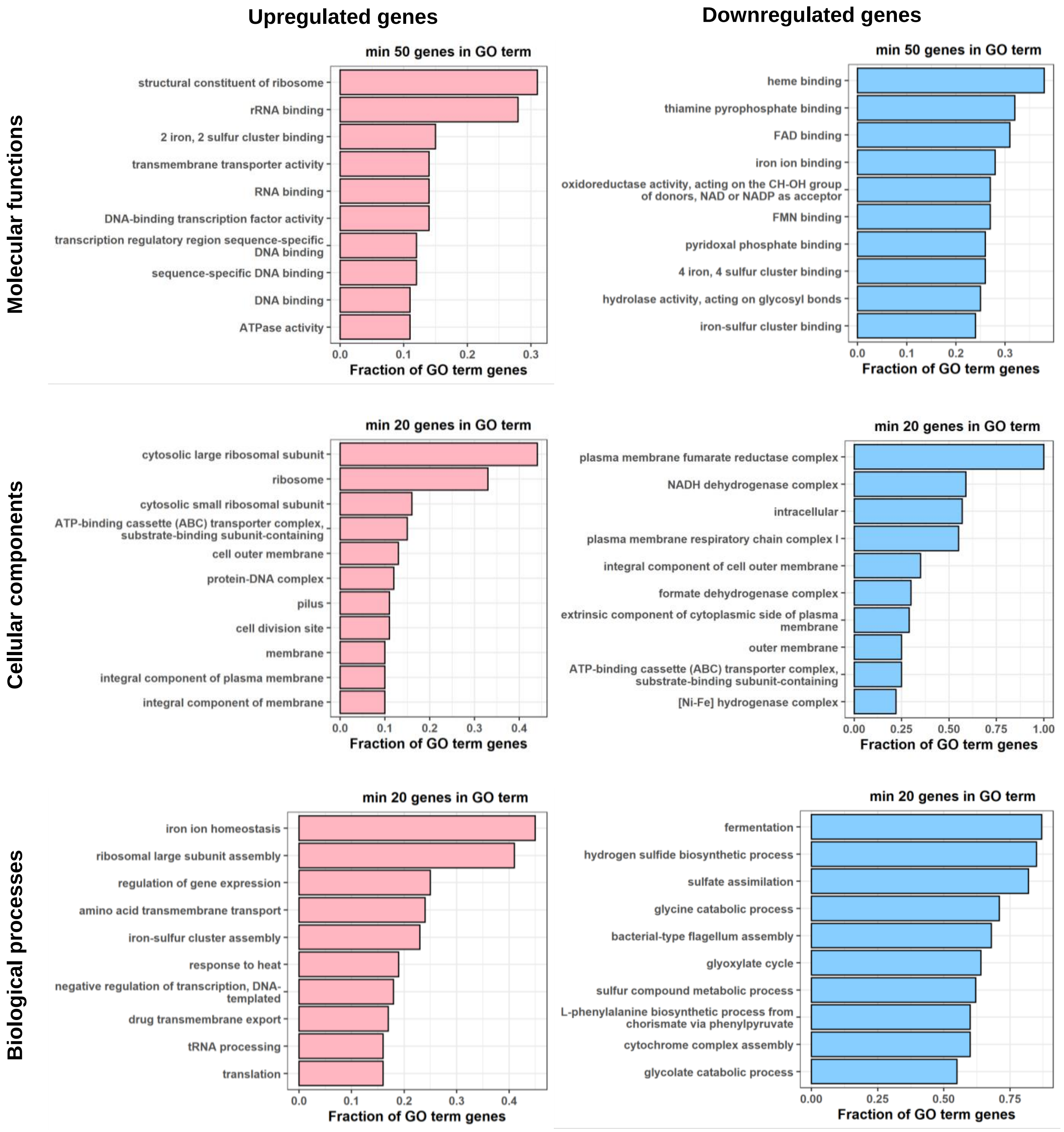

**Supplementary Figure 4. Enrichment analysis of GO terms.** GO terms in molecular functions, biological processes or cellular components associated with up- or downregulated genes upon treatment with TAT-RasGAP$_{317-326}$. The top 10 groups having the highest enrichment in up- or down-regulated genes are shown.

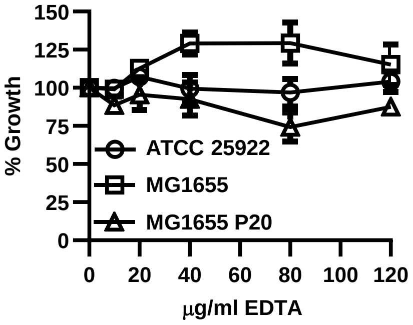

**Supplementary Figure 5.** Overnight growth of *E.coli* strains MG1655, ATCC 25922 or MG1655 P20 that were selected for resistance to TAT-RasGAP$_{317\text{-}326}$ for 20 passages (see Fig. 8A) in presence of indicated concentrations of EDTA was measured and expressed as percentage of a control grown in absence of EDTA. Error bars represent the range of two independent experiments.

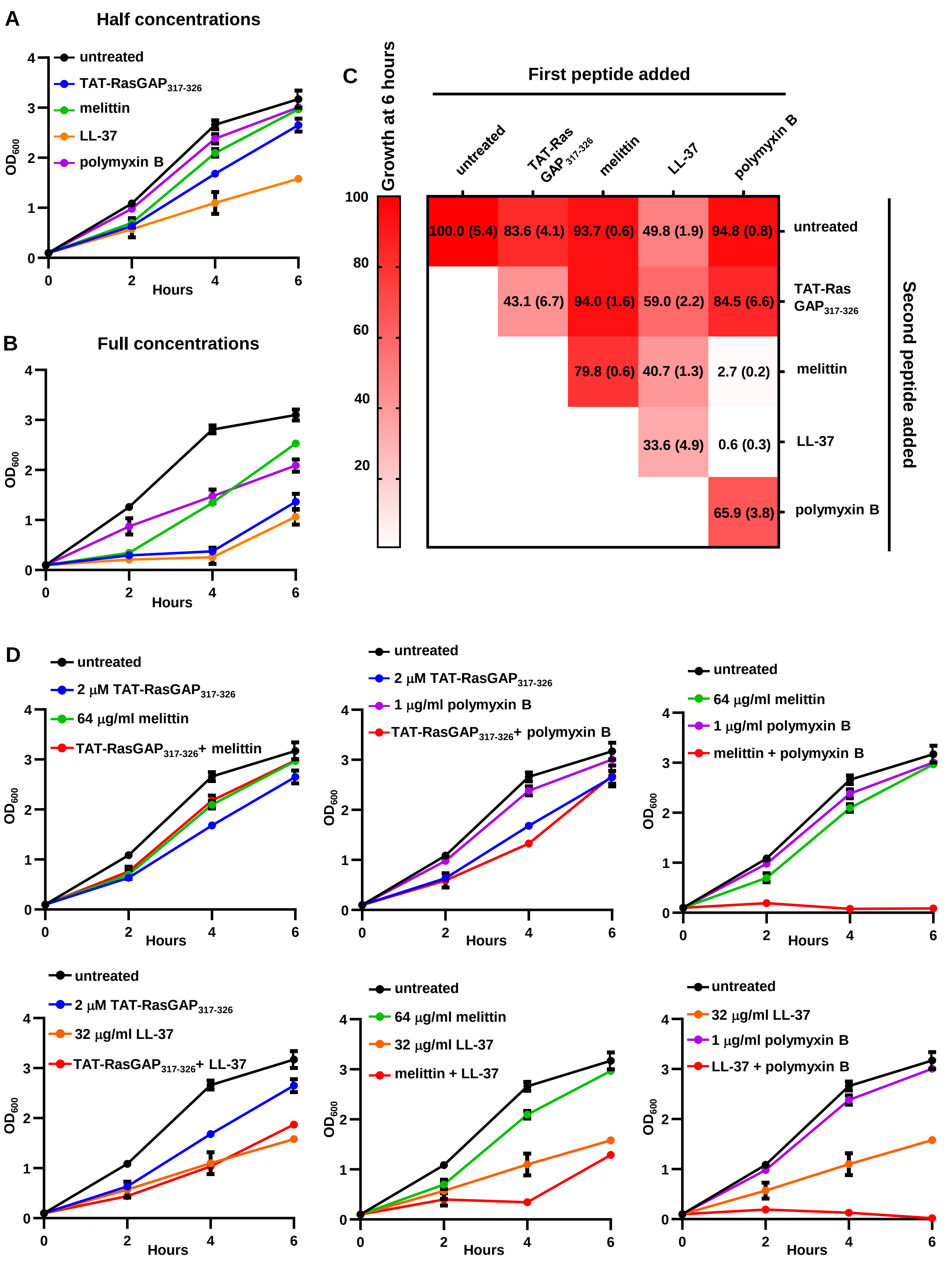
A
Half concentrations
untreated
TAT-RasGAP317-326
melittin
LL-37
polymyxin B
OD600
Hours
B
Full concentrations
OD600
Hours
C
Growth at 6 hours
First peptide added
untreated
TAT-Ras GAP 317-326
melittin
LL-37
polymyxin B
100.0 (5.4)
83.6 (4.1)
93.7 (0.6)
49.8 (1.9)
94.8 (0.8)
untreated
43.1 (6.7)
94.0 (1.6)
59.0 (2.2)
84.5 (6.6)
TAT-Ras GAP317-326
79.8 (0.6)
40.7 (1.3)
2.7 (0.2)
melittin
33.6 (4.9)
0.6 (0.3)
LL-37
65.9 (3.8)
polymyxin B
Second peptide added
D
untreated
2 μM TAT-RasGAP317-326
64 μg/ml melittin
TAT-RasGAP317-326+ melittin
1 μg/ml polymyxin B
TAT-RasGAP317-326+ polymyxin B
melittin + polymyxin B
32 μg/ml LL-37
TAT-RasGAP317-326+ LL-37
melittin + LL-37
LL-37 + polymyxin B
OD600
Hours

**Supplementary Figure 6. Effect of melittin, LL-37 and polymyxin B on TAT-RasGAP$_{317\text{-}326}$ activity. (A-B)** *E. coli* MG1655 were grown overnight at 37°C, diluted to $OD_{600}$ = 0.1, and grown further during 1 hour before addition of the indicated AMPs. $OD_{600}$ was measured at the indicated time points. For each AMP, two concentrations were tested: (A) a "half" concentration (2 μM TAT-RasGAP$_{317\text{-}326}$, 64 μg/ml melittin, 32 μg/ml LL-37 or 1 μg/ml polymyxin B) and (B) a "full" concentration corresponding to twice that of panel A (4 μM TAT-RasGAP$_{317\text{-}326}$, 128 μg/ml melittin, 64 μg/ml LL-37 or 2 μg/ml polymyxin B). Average and range of two independent experiments are presented. **(C)** Combinations of "half" concentrations of AMPs were then tested and growth at 6h was expressed as percentage of growth compared to an untreated control for each combination of AMPs. Average and range (in brackets) of two independent experiments are shown. **D)** Detailed growth curves of *E. coli* in the presence of combinations of AMPs used to produce panel C. Average and range of two independent experiments are shown.

**Table S1.** List of genes, extracted from Dataset 1, which were upregulated in the presence of TAT-RasGAP$_{317-326}$

| WT_TAT-RasGAP | WT_untreated | FoldChange | $Log_2$FoldChange | Mean_WT_untr_TAT | $Log_2$_mean_WT_untr_TAT | Locus_tag | Gene | Product |
|---|---|---|---|---|---|---|---|---|
| 88.81641406 | 0 | Inf | Inf | 44.408207 | 5.472754419 | b1574 | *dicF* | Qin prophage; small regulatory RNA DicF |
| 264.0399693 | 0 | Inf | Inf | 132.019985 | 7.044612525 | b1909 | *leuZ* | tRNA-Leu |
| 106.5329514 | 0 | Inf | Inf | 53.2664757 | 5.735155925 | b3969 | *gltT* | tRNA-Glu |
| 1491.176548 | 0 | Inf | Inf | 745.588274 | 9.54223536 | b4663 | *azuC* | uncharacterized protein AzuC |
| 33.79578422 | 0 | Inf | Inf | 16.8978921 | 4.078771387 | b1165 | *ymgA* | putative two-component system connector protein YmgA |
| 44.95230759 | 0 | Inf | Inf | 22.4761538 | 4.490323272 | b1352 | *kilR* | Rac prophage; inhibitor of FtsZ, killing protein |
| 66.02404599 | 0 | Inf | Inf | 33.012023 | 5.044919645 | b1666 | *valW* | tRNA-Val |
| 66.7576465 | 0 | Inf | Inf | 33.3788233 | 5.060861189 | b2646 | *ypjF* | CP4-57 prophage; toxin of the YpjF-YfjZ toxin-antitoxin system |
| 62.21304138 | 0 | Inf | Inf | 31.1065207 | 4.959145131 | b4222 | *ytfP* | gamma-glutamylamine cyclotransferase family protein YtfP |
| 44.83114234 | 0 | Inf | Inf | 22.4155712 | 4.486429356 | b4732 | *ykiD* | protein YkiD |
| 2534.24491 | 3.933658 | 644.2463533 | 9.3314687 | 1269.08928 | 10.30957786 | b3686 | *ibpB* | small heat shock protein IbpB |
| 19731.87503 | 32.83591 | 600.9236468 | 9.2310379 | 9882.35547 | 13.27063924 | b3687 | *ibpA* | small heat shock protein IbpA |
| 7482.81578 | 21.41254 | 349.4595737 | 8.4489818 | 3752.11416 | 11.87348801 | b4062 | *soxS* | DNA-binding transcriptional dual regulator SoxS |
| 531.3075121 | 3.532691 | 150.3973895 | 7.2326357 | 267.420102 | 8.062964105 | b0399 | *phoB* | DNA-binding transcriptional dual regulator PhoB |
| 4471.383418 | 34.89665 | 128.1321795 | 7.001489 | 2253.14003 | 11.13772126 | b1306 | *pspC* | phage shock protein C |
| 707.5817757 | 5.552815 | 127.4275705 | 6.9935336 | 356.567296 | 8.478030574 | b3275 | *rrlD* | 23S ribosomal RNA |
| 3580.002669 | 28.37237 | 126.1791814 | 6.9793301 | 1804.18752 | 10.81713358 | b0748 | *lysZ* | tRNA-Lys |
| 560.9135856 | 4.640384 | 120.8765416 | 6.9173905 | 282.776985 | 8.143520894 | b0805 | *fiu* | putative iron siderophore outer membrane transporter |
| 2107.173034 | 17.912 | 117.6402796 | 6.8782383 | 1062.54252 | 10.05330486 | b2673 | *nrdH* | glutaredoxin-like protein |
| 57.10458622 | 0.512307 | 111.4655537 | 6.8004541 | 28.8084466 | 4.848419966 | b1600 | *mdtJ* | multidrug/spermidine efflux pump membrane subunit MdtJ |
| 7172.338494 | 73.91639 | 97.03312246 | 6.6004054 | 3623.12744 | 11.82301984 | b4484 | *cpxP* | periplasmic protein CpxP |
| 735.1848713 | 7.672813 | 95.8168611 | 6.5822076 | 371.428842 | 8.536942038 | b2685 | *emrA* | multidrug efflux pump membrane fusion protein EmrA |
| 1401.900739 | 15.62536 | 89.71955564 | 6.4873506 | 708.763052 | 9.469159587 | b2606 | *rplS* | 50S ribosomal subunit protein L19 |
| 129.5966588 | 1.55864 | 83.14727787 | 6.3775971 | 65.5776493 | 6.035132283 | b0585 | *fes* | enterochelin esterase |
| 782.8486774 | 9.713064 | 80.5974979 | 6.3326631 | 396.280871 | 8.630379517 | b1307 | *pspD* | phage shock protein D |
| 6243.016765 | 78.45235 | 79.5771801 | 6.3142829 | 3160.73456 | 11.62604417 | b3343 | *tusB* | sulfurtransferase complex subunit TusB |
| 456.9014464 | 6.228292 | 73.35902728 | 6.1969026 | 231.564869 | 7.855272588 | b0595 | *entB* | enterobactin synthase component B |
| 805.1392969 | 11.96317 | 67.30150363 | 6.0725668 | 408.551233 | 8.674373197 | b1531 | *marA* | DNA-binding transcriptional dual regulator MarA |
| 37.65815956 | 0.568195 | 66.27681574 | 6.0504324 | 19.1131773 | 4.256495722 | b4742 | *ymjE* | protein YmjE |
| 316.3386228 | 5.229372 | 60.49265725 | 5.9186881 | 160.783998 | 7.328980015 | b0584 | *fepA* | ferric enterobactin outer membrane transporter |

| | | | | | | | | |
|---|---|---|---|---|---|---|---|---|
| 7105.676286 | 118.7528 | 59.83587904 | 5.9029389 | 3612.21453 | 11.81866786 | b1305 | *pspB* | phage shock protein B |
| 381.4435646 | 6.822342 | 55.91094034 | 5.8050587 | 194.132953 | 7.600901221 | b3005 | *exbD* | Ton complex subunit ExbD |
| 64.43284393 | 1.246705 | 51.68252699 | 5.6916047 | 32.8397743 | 5.037372305 | b1018 | *efeO* | ferrous iron transport system protein EfeO |
| 495.2866638 | 9.850773 | 50.27896365 | 5.651883 | 252.568718 | 7.980532157 | b1228 | *ychS* | putative uncharacterized protein YchS |
| 34.64426395 | 0.69618 | 49.76340035 | 5.6370132 | 17.6702218 | 4.143248242 | b0353 | *mhpT* | 3-hydroxyphenylpropionate/3- hydroxycinnamate:H(+) symporter |
| 53.69371498 | 1.08384 | 49.54024608 | 5.6305291 | 27.3887776 | 4.775512975 | b2539 | *hcaF* | putative 3-phenylpropionate/cinnamate dioxygenase subunit beta |
| 39.94047226 | 0.811707 | 49.20551467 | 5.6207481 | 20.3760897 | 4.348805314 | b1569 | *dicC* | Qin prophage; DNA-binding transcriptional regulator for DicB |
| 33.5145433 | 0.707945 | 47.34058262 | 5.5650056 | 17.1112443 | 4.096872769 | b3457 | *livH* | branched chain amino acid/phenylalanine ABC transporter membrane subunit LivH |
| 534.3387506 | 11.40229 | 46.86239497 | 5.5503588 | 272.870522 | 8.092072738 | b2864 | *glyU* | tRNA-Gly |
| 159.1212921 | 3.426973 | 46.432026 | 5.5370483 | 81.2741325 | 6.344724346 | b1505 | *ydeT* | fimbrial usher domain-containing protein YdeT |
| 406.0729254 | 9.08342 | 44.70485167 | 5.4823595 | 207.578172 | 7.697510937 | b2155 | *cirA* | ferric dihyroxybenzoylserine outer membrane transporter |
| 141.2180984 | 3.196097 | 44.18454383 | 5.4654699 | 72.2070978 | 6.174068753 | b0883 | *serW* | tRNA-Ser |
| 745.2072421 | 17.16589 | 43.41208675 | 5.4400249 | 381.186568 | 8.574353472 | b3556 | *cspA* | cold shock protein CspA |
| 34.60044575 | 0.801301 | 43.18034961 | 5.432303 | 17.7008732 | 4.14574863 | b1375 | *ynaE* | Rac prophage; uncharacterized protein YnaE |
| 358.323094 | 8.750204 | 40.95025607 | 5.3558006 | 183.536649 | 7.519924362 | b2106 | *rcnA* | Ni(2(+))/Co(2(+)) exporter |
| 903.9401136 | 22.27063 | 40.58888078 | 5.3430127 | 463.105374 | 8.855196688 | b0802 | *ybiJ* | DUF1471 domain-containing protein YbiJ |
| 1956.519607 | 49.38387 | 39.61859841 | 5.3081059 | 1002.95174 | 9.970036469 | b4050 | *pspG* | phage shock protein G |
| 203.7175411 | 5.261065 | 38.72172749 | 5.2750714 | 104.489303 | 6.707211449 | b3941 | *metF* | 5,10-methylenetetrahydrofolate reductase |
| 116.9125869 | 3.063797 | 38.15937875 | 5.2539658 | 59.9881919 | 5.906606643 | b4455 | *hokA* | small toxic polypeptide |
| 1285.398513 | 35.00082 | 36.72481563 | 5.1986833 | 660.199664 | 9.366758595 | b0665 | *glnV* | tRNA-Gln |
| 409.5008321 | 11.43759 | 35.8030806 | 5.1620118 | 210.469209 | 7.717465379 | b3728 | *pstS* | phosphate ABC transporter periplasmic binding protein |
| 467.4042158 | 13.2356 | 35.31416192 | 5.142175 | 240.319909 | 7.908812363 | b1230 | *tyrV* | tRNA-Tyr |
| 2523.585758 | 71.83609 | 35.12977598 | 5.1346225 | 1297.71092 | 10.34175333 | b0666 | *metU* | tRNA-Met |
| 3317.408637 | 99.0999 | 33.47539713 | 5.0650293 | 1708.25427 | 10.73830702 | b0150 | *fhuA* | ferrichrome outer membrane transporter/phage receptor |
| 868.7803478 | 26.3164 | 33.01288359 | 5.0449573 | 447.548375 | 8.805899821 | b0843 | *ybjH* | uncharacterized protein YbjH |
| 102.9167094 | 3.202714 | 32.13421369 | 5.0060383 | 53.0597119 | 5.729544937 | b3088 | *alx* | putative membrane-bound redox modulator Alx |
| 376.5815956 | 11.76975 | 31.99570414 | 4.9998063 | 194.175675 | 7.601218673 | b1436 | *yncJ* | protein YncJ |
| 132.4158863 | 4.192171 | 31.58647141 | 4.9812349 | 68.3040286 | 6.093898767 | b0327 | *yahM* | uncharacterized protein YahM |
| 5667.018257 | 181.058 | 31.29945783 | 4.9680658 | 2924.03814 | 11.51374642 | b1304 | *pspA* | phage shock protein A |
| 86.25611967 | 2.793361 | 30.87897095 | 4.9485528 | 44.5247404 | 5.476535296 | b2968 | *yghD* | putative type II secretion system M-type protein |
| 280.1164821 | 9.283689 | 30.17297059 | 4.9151848 | 144.700086 | 7.176921966 | b4354 | *btsT* | pyruvate:H(+) symporter |
| 111.4594953 | 3.77164 | 29.5520001 | 4.8851839 | 57.6155675 | 5.848386769 | b0527 | *ybcI* | conserved inner membrane protein YbcI |
| 166.1954491 | 5.73993 | 28.95426546 | 4.855704 | 85.9676894 | 6.425722626 | b0593 | *entC* | isochorismate synthase EntC |

| | | | | | | | | |
|---|---|---|---|---|---|---|---|---|
| 239.3774484 | 8.721134 | 27.44797419 | 4.7786278 | 124.049291 | 6.954769679 | b1112 | *bhsA* | DUF1471 domain-containing multiple stress resistance outer membrane protein BhsA |
| 97.10594604 | 3.655746 | 26.56255566 | 4.7313221 | 50.3808458 | 5.654803438 | b2863 | *ygeQ* | protein YgeQ |
| 39.22724954 | 1.48813 | 26.36009716 | 4.7202838 | 20.3576897 | 4.347501943 | b0021 | *insB-1* | IS1 protein InsB |
| 1249.756608 | 47.44966 | 26.33857871 | 4.7191056 | 648.603134 | 9.341192185 | b4242 | *mgtA* | Mg(2(+)) importing P-type ATPase |
| 1163.507275 | 44.61311 | 26.07994154 | 4.7048687 | 604.060192 | 9.238548505 | b1530 | *marR* | DNA-binding transcriptional repressor MarR |
| 478.159526 | 18.50372 | 25.8412635 | 4.6916047 | 248.331623 | 7.956124181 | b3273 | *thrV* | tRNA-Thr |
| 1683.205617 | 65.34243 | 25.75976369 | 4.6870475 | 874.274025 | 9.771941726 | b4736 | *yliM* | protein YliM |
| 465.9901191 | 18.18046 | 25.63137787 | 4.6798391 | 242.085287 | 7.919371592 | b3828 | *metR* | DNA-binding transcriptional dual regulator MetR |
| 76.76730556 | 3.024264 | 25.38379726 | 4.665836 | 39.8957848 | 5.318164421 | b1527 | *yneK* | protein YneK |
| 36.44338022 | 1.440126 | 25.30569327 | 4.6613901 | 18.941753 | 4.243497947 | b1040 | *csgD* | DNA-binding transcriptional dual regulator CsgD |
| 64.4865216 | 2.573589 | 25.05703566 | 4.6471438 | 33.5300555 | 5.067382966 | b4482 | *yigE* | DUF2233 domain-containing protein YigE |
| 162.0176632 | 6.54085 | 24.77012304 | 4.6305291 | 84.2792567 | 6.397105685 | b3185 | *rpmA* | 50S ribosomal subunit protein L27 |
| 1653.76163 | 67.51337 | 24.49532173 | 4.6144343 | 860.637498 | 9.74926189 | b3938 | *metJ* | DNA-binding transcriptional repressor MetJ |
| 140.7008159 | 5.838048 | 24.10066025 | 4.5910008 | 73.269432 | 6.195139527 | b2105 | *rcnR* | DNA-binding transcriptional repressor RcnR |
| 7479.869132 | 311.8715 | 23.98381659 | 4.5839894 | 3895.87032 | 11.92772994 | b0014 | *dnaK* | chaperone protein DnaK |
| 427.0117365 | 18.21822 | 23.43871953 | 4.5508219 | 222.614978 | 7.798406853 | b3960 | *argH* | argininosuccinate lyase |
| 97.39179197 | 4.310445 | 22.59436899 | 4.4978914 | 50.8511186 | 5.668207606 | b4517 | *gnsA* | putative phosphatidylethanolamine synthesis regulator GnsA |
| 1520.516414 | 68.20811 | 22.29231059 | 4.4784743 | 794.362263 | 9.633653277 | b0156 | *erpA* | iron-sulfur cluster insertion protein ErpA |
| 264.8108024 | 11.98658 | 22.0922719 | 4.4654699 | 138.398692 | 7.112686494 | b1573 | *ydfC* | Qin prophage; uncharacterized protein YdfC |
| 34.35963464 | 1.596753 | 21.51844666 | 4.427502 | 17.9781936 | 4.168176165 | b2674 | *nrdI* | dimanganese-tyrosyl radical cofactor maintenance flavodoxin NrdI |
| 239.0658444 | 11.23622 | 21.27636414 | 4.4111797 | 125.151031 | 6.967526362 | b0803 | *ybiI* | zinc finger domain-containing protein YbiI |
| 62.76359927 | 2.97626 | 21.08807773 | 4.3983557 | 32.8699296 | 5.03869646 | b4678 | *yoeI* | uncharacterized protein YoeI |
| 351.323074 | 16.76868 | 20.95114216 | 4.388957 | 184.045879 | 7.523921635 | b3909 | *kdgT* | 2-dehydro-3-deoxy-D-gluconate:H(+) symporter |
| 70.60904918 | 3.472303 | 20.3349321 | 4.3458883 | 37.0406762 | 5.21103853 | b1557 | *cspB* | Qin prophage; cold shock-like protein CspB |
| 87.2984608 | 4.403512 | 19.82473666 | 4.3092298 | 45.8509863 | 5.518880862 | b4183 | *yjfK* | conserved protein YjfK |
| 127.2248229 | 6.544782 | 19.43912418 | 4.2808913 | 66.8848023 | 6.06360653 | b0586 | *entF* | apo-serine activating enzyme |
| 155.6333153 | 8.130271 | 19.14245151 | 4.2587037 | 81.8817931 | 6.35547079 | b3058 | *folB* | dihydroneopterin aldolase |
| 144.4227451 | 7.564473 | 19.0922418 | 4.2549146 | 75.993609 | 6.247806189 | b0153 | *fhuB* | iron(III) hydroxamate ABC transporter membrane subunit |
| 1704.683041 | 89.62473 | 19.02023051 | 4.2494628 | 897.153886 | 9.809211657 | b0966 | *hspQ* | heat shock protein, hemimethylated DNA-binding protein |
| 32.07917296 | 1.736152 | 18.47717287 | 4.2076721 | 16.9076623 | 4.079605295 | b3466 | *yhhL* | DUF1145 domain-containing protein YhhL |
| 185.0583378 | 10.19337 | 18.15477369 | 4.182277 | 97.6258542 | 6.609191362 | b0066 | *thiQ* | thiamine ABC transporter ATP binding subunit |
| 292.100488 | 16.32824 | 17.88928701 | 4.161024 | 154.214362 | 7.268793316 | b0349 | *mhpC* | 2-hydroxy-6-ketonona-2,4-dienedioate hydrolase |
| 218.328498 | 12.27915 | 17.78042391 | 4.1522178 | 115.303824 | 6.849296552 | b0594 | *entE* | 2,3-dihydroxybenzoate-AMP ligase |

| | | | | | | | | |
|---|---|---|---|---|---|---|---|---|
| 121.4608245 | 6.866357 | 17.68926666 | 4.1448023 | 64.1635909 | 6.003682979 | b1020 | *phoH* | ATP-binding protein PhoH |
| 110.8652102 | 6.403838 | 17.31230762 | 4.1137261 | 58.634524 | 5.873678469 | b2128 | *yehW* | glycine betaine ABC transporter membrane subunit YehW |
| 1335.514055 | 78.32461 | 17.05101426 | 4.0917857 | 706.919333 | 9.465401787 | b4567 | *yjjZ* | protein YjjZ |
| 1679.752117 | 99.92009 | 16.81095437 | 4.0713297 | 889.836105 | 9.797395826 | b0747 | *lysY* | tRNA-Lys |
| 348.8382655 | 20.83382 | 16.7438464 | 4.0655591 | 184.836042 | 7.530102293 | b4372 | *holD* | DNA polymerase III subunit psi |
| 838.7982716 | 51.37832 | 16.32591977 | 4.0290924 | 445.088294 | 8.797947748 | b1452 | *yncE* | PQQ-like domain-containing protein YncE |
| 64.66552652 | 4.006504 | 16.14013914 | 4.0125811 | 34.3360151 | 5.10165071 | b0451 | *amtB* | ammonia/ammonium transporter |
| 67.80301235 | 4.219989 | 16.06710684 | 4.0060383 | 36.0115006 | 5.170385814 | b0252 | *yafZ* | CP4-6 prophage; conserved protein YafZ |
| 338.5747494 | 21.70189 | 15.60116074 | 3.9635815 | 180.138322 | 7.492961318 | b2963 | *mltC* | membrane-bound lytic murein transglycosylase C |
| 38.51402683 | 2.485853 | 15.4932816 | 3.9535708 | 20.4999401 | 4.35754779 | b1365 | *ynaK* | Rac prophage; ParB-like nuclease domain-containing protein YnaK |
| 162.7474725 | 10.53804 | 15.44381389 | 3.9489572 | 86.6427544 | 6.437007203 | b2527 | *hscB* | co-chaperone for [Fe-S] cluster biosynthesis |
| 74.08936155 | 4.816842 | 15.3813157 | 3.943107 | 39.4531016 | 5.302066816 | b0151 | *fhuC* | iron(III) hydroxamate ABC transporter ATP binding subunit |
| 41.84239951 | 2.741292 | 15.26375149 | 3.9320377 | 22.2918457 | 4.47844417 | b4347 | *symE* | toxic protein SymE |
| 106.0423797 | 6.996432 | 15.15663746 | 3.9218778 | 56.5194057 | 5.820674392 | b0590 | *fepD* | ferric enterobactin ABC transporter membrane subunit FebD |
| 94.14539891 | 6.250146 | 15.06291266 | 3.9129289 | 50.1977723 | 5.649551436 | b3272 | *rrfF* | 5S ribosomal RNA |
| 665.9267827 | 44.26496 | 15.04410753 | 3.9111266 | 355.09587 | 8.472064771 | b0015 | *dnaJ* | chaperone protein DnaJ |
| 584.973154 | 41.25913 | 14.17802868 | 3.825585 | 313.116143 | 8.290554079 | b3743 | *asnC* | DNA-binding transcriptional dual regulator AsnC |
| 146.8606387 | 10.36559 | 14.16808616 | 3.824573 | 78.6131167 | 6.296698143 | b2723 | *hycC* | formate hydrogenlyase subunit HycC |
| 9868.615318 | 709.6254 | 13.90679576 | 3.7977181 | 5289.12035 | 12.36881209 | b1829 | *htpX* | zinc dependent endoprotease |
| 182.245943 | 13.19191 | 13.81498203 | 3.7881618 | 97.7189245 | 6.610566079 | b3063 | *ttdT* | L-tartrate:succinate antiporter |
| 38.42390282 | 2.791459 | 13.764808 | 3.7829126 | 20.6076811 | 4.365110269 | b2852 | *ygeH* | putative transcriptional regulator YgeH |
| 66.88675543 | 5.018365 | 13.32839545 | 3.7364312 | 35.9525603 | 5.168022608 | b0282 | *yagP* | putative LysR family substrate binding domain-containing protein YagP |
| 273.184192 | 20.60556 | 13.2577902 | 3.7287684 | 146.894876 | 7.198640263 | b3426 | *glpD* | aerobic glycerol 3-phosphate dehydrogenase |
| 87.37066797 | 6.591345 | 13.25536314 | 3.7285043 | 46.9810063 | 5.554005712 | b3547 | *yhjX* | putative pyruvate transporter |
| 87.7486292 | 6.66982 | 13.15607203 | 3.7176569 | 47.2092245 | 5.560996878 | b0591 | *entS* | enterobactin exporter EntS |
| 35.3305029 | 2.69046 | 13.13177001 | 3.7149895 | 19.0104815 | 4.248723167 | b0785 | *moaE* | molybdopterin synthase catalytic subunit |
| 5406.698687 | 412.071 | 13.12079364 | 3.7137831 | 2909.38485 | 11.50649843 | b3461 | *rpoH* | RNA polymerase, sigma 32 (sigma H) factor |
| 48.20080336 | 3.676556 | 13.11031287 | 3.7126302 | 25.9386798 | 4.697033149 | b2755 | *cas1* | multifunctional nuclease Cas1 |
| 1238.046644 | 96.11817 | 12.88046398 | 3.6871127 | 667.082408 | 9.381721185 | b4171 | *miaA* | tRNA dimethylallyltransferase |
| 140.9144035 | 11.08897 | 12.70762086 | 3.667622 | 76.0016858 | 6.247959515 | b4063 | *soxR* | DNA-binding transcriptional dual regulator SoxR |
| 2505.349742 | 197.6339 | 12.67671952 | 3.6641095 | 1351.49183 | 10.40033708 | b3301 | *rplO* | 50S ribosomal subunit protein L15 |
| 322.376669 | 25.9239 | 12.4355005 | 3.6363927 | 174.150284 | 7.444189019 | b3238 | *yhcN* | DUF1471 domain-containing stress-induced protein YhcN |
| 256.5121014 | 20.65266 | 12.42029641 | 3.6346277 | 138.582378 | 7.114600011 | b2181 | *yejG* | protein YejG |

| | | | | | | | | |
|---|---|---|---|---|---|---|---|---|
| 52.07054162 | 4.243926 | 12.26942704 | 3.616996 | 28.1572339 | 4.815433707 | b1690 | *ydiM* | putative exporter YdiM |
| 848.9042749 | 69.38721 | 12.23430464 | 3.6128602 | 459.145743 | 8.842808359 | b4532 | *hicA* | toxin of the HicA-HicB toxin-antitoxin system |
| 4196.210329 | 344.3487 | 12.18593471 | 3.607145 | 2270.2795 | 11.1486542 | b3066 | *dnaG* | DNA primase |
| 893.9789589 | 76.72455 | 11.65179876 | 3.5424808 | 485.351752 | 8.922886889 | b1641 | *slyB* | outer membrane lipoprotein SlyB |
| 155.7678418 | 13.49463 | 11.5429478 | 3.5289398 | 84.6312373 | 6.403118354 | b1846 | *yebE* | conserved inner membrane protein YebE |
| 43.02661837 | 3.734364 | 11.52180688 | 3.5262951 | 23.3804911 | 4.547233327 | b3181 | *greA* | transcription elongation factor GreA |
| 209.9334183 | 18.31939 | 11.45962767 | 3.5184883 | 114.126405 | 6.834488817 | b4368 | *leuV* | tRNA-Leu |
| 53.84934894 | 4.710255 | 11.43236448 | 3.5150519 | 29.2798018 | 4.871833885 | b3907 | *rhaT* | rhamnose/lyxose:H(+) symporter |
| 32.61104064 | 2.880251 | 11.32228935 | 3.5010938 | 17.7456461 | 4.149393194 | b0532 | *sfmD* | putative fimbrial usher protein SfmD |
| 151.1608468 | 13.38212 | 11.29573335 | 3.497706 | 82.2714829 | 6.362320542 | b1907 | *tyrP* | tyrosine:H(+) symporter |
| 138.9765412 | 12.43651 | 11.17487879 | 3.4821873 | 75.7065278 | 6.242345797 | b0685 | *ybfE* | LexA-regulated protein |
| 215.2595319 | 19.28756 | 11.16053782 | 3.4803346 | 117.273545 | 6.873733797 | b4011 | *yjaA* | stress response protein |
| 128.7685112 | 11.59065 | 11.10969255 | 3.473747 | 70.1795785 | 6.132979378 | b1013 | *rutR* | DNA-binding transcriptional dual regulator RutR |
| 51.32113803 | 4.625559 | 11.09512103 | 3.4718535 | 27.9733486 | 4.805981058 | b3051 | *yqiK* | protein YqiK |
| 30.17901289 | 2.722189 | 11.08630372 | 3.4707065 | 16.4506009 | 4.040068377 | b3137 | *kbaY* | tagatose-1,6-bisphosphate aldolase 1 subunit KbaY |
| 84.18771248 | 7.662438 | 10.98706571 | 3.4577342 | 45.9250754 | 5.521210184 | b0157 | *yadS* | conserved inner membrane protein YadS |
| 278.00809 | 25.69078 | 10.82131636 | 3.4358041 | 151.849436 | 7.246497744 | b2531 | *iscR* | DNA-binding transcriptional dual regulator IscR |
| 65.34798277 | 6.066318 | 10.77226481 | 3.4292497 | 35.7071503 | 5.158141097 | b4416 | *rybA* | small RNA RybA |
| 120.1725418 | 11.27649 | 10.6569134 | 3.4137177 | 65.7245142 | 6.038359669 | b2675 | *nrdE* | ribonucleoside-diphosphate reductase 2 subunit alpha |
| 75.77229399 | 7.144932 | 10.60504131 | 3.4066783 | 41.4586128 | 5.373599938 | b0260 | *mmuP* | S-methyl-L-methionine transporter |
| 73.83952855 | 6.985457 | 10.57046503 | 3.4019669 | 40.4124928 | 5.336729439 | b2236 | *yfaE* | ferredoxin-like diferric-tyrosyl radical cofactor maintenance protein YfaE |
| 87.35140105 | 8.376484 | 10.4281703 | 3.3824141 | 47.8639425 | 5.58086733 | b2618 | *yfjF* | putative component of the Rsx system |
| 197.1536692 | 18.91254 | 10.42449194 | 3.3819052 | 108.033107 | 6.755329686 | b3151 | *yraQ* | permease family protein YraQ |
| 123.2026208 | 11.8637 | 10.38483735 | 3.3764067 | 67.5331617 | 6.077524196 | b1160 | *iraM* | anti-adaptor protein IraM, inhibitor of sigma(S) proteolysis |
| 3445.422726 | 331.853 | 10.38237711 | 3.3760649 | 1888.63785 | 10.88313037 | b3309 | *rplX* | 50S ribosomal subunit protein L24 |
| 194.227895 | 18.79267 | 10.33530187 | 3.3695086 | 106.510281 | 6.73484889 | b2811 | *csdE* | sulfur acceptor protein CsdE |
| 189.4977901 | 18.42992 | 10.28207516 | 3.3620596 | 103.963853 | 6.699938204 | b3820 | *yigI* | putative thioesterase YigI |
| 109.0293382 | 10.71454 | 10.17583433 | 3.3470752 | 59.8719369 | 5.903808037 | b2668 | *ygaP* | thiosulfate sulfurtransferase YgaP |
| 156.9089982 | 15.62536 | 10.04194177 | 3.3279664 | 86.2671812 | 6.430739912 | b4734 | *yldA* | protein YldA |
| 369.7796292 | 37.10062 | 9.966939721 | 3.3171506 | 203.440124 | 7.668460436 | b4043 | *lexA* | DNA-binding transcriptional repressor LexA |
| 55.58035472 | 5.588471 | 9.945539134 | 3.3140496 | 30.5844127 | 4.93472467 | b0984 | *gfcD* | putative lipoprotein GfcD |
| 267.2822478 | 26.97306 | 9.909228886 | 3.3087728 | 147.127655 | 7.20092464 | b4367 | *fhuF* | hydroxamate siderophore iron reductase |
| 520.681696 | 52.61602 | 9.895877168 | 3.3068276 | 286.648859 | 8.163140728 | b4698 | *mgrR* | small regulatory RNA MgrR |

| | | | | | | | | |
|---|---|---|---|---|---|---|---|---|
| 181.0834945 | 18.31681 | 9.886189207 | 3.3054145 | 99.7001545 | 6.639523836 | b2817 | *amiC* | N-acetylmuramoyl-L-alanine amidase C |
| 349.0326876 | 35.42544 | 9.852598766 | 3.3005043 | 192.229066 | 7.586682684 | b0217 | *yafT* | lipoprotein YafT |
| 14391.83375 | 1461.275 | 9.848816424 | 3.2999504 | 7926.55461 | 12.9524782 | b3854 | *rrlA* | 23S ribosomal RNA |
| 952.9496378 | 96.97594 | 9.826659985 | 3.2967011 | 524.962791 | 9.03607136 | b0161 | *degP* | periplasmic serine endoprotease DegP |
| 134.8569228 | 13.72498 | 9.825653799 | 3.2965534 | 74.2909525 | 6.215114618 | b4012 | *yjaB* | putative N-acetyltransferase YjaB |
| 572.7178434 | 58.59512 | 9.774156662 | 3.2889722 | 315.65648 | 8.302211556 | b2748 | *ftsB* | cell division protein FtsB |
| 128.9730432 | 13.2356 | 9.744402761 | 3.2845738 | 71.1043229 | 6.151865369 | b1298 | *puuD* | gamma-glutamyl-gamma-aminobutyrate hydrolase |
| 939.0400045 | 96.55674 | 9.725266433 | 3.2817378 | 517.798371 | 9.016246617 | b0817 | *mntR* | DNA-binding transcriptional dual regulator MntR |
| 173.2941856 | 17.97608 | 9.640264107 | 3.2690727 | 95.6351342 | 6.579468824 | b3335 | *gspO* | Type II secretion system prepilin peptidase |
| 2084.60142 | 219.8944 | 9.480009284 | 3.2448885 | 1152.24793 | 10.17023547 | b3414 | *nfuA* | iron-sulfur cluster carrier protein NfuA |
| 149.0176684 | 15.83858 | 9.408526986 | 3.2339689 | 82.4281221 | 6.365064723 | b2686 | *emrB* | multidrug efflux pump membrane subunit EmrB |
| 453.0747322 | 48.82926 | 9.2787542 | 3.2139311 | 250.951998 | 7.971267621 | b4599 | *mgtS* | small protein MgtS |
| 288.7125566 | 31.25073 | 9.238586432 | 3.2076721 | 159.981643 | 7.321762559 | b3195 | *mlaF* | intermembrane phospholipid transport system, ATP binding subunit MlaF |
| 29.85098014 | 3.239405 | 9.214958334 | 3.2039776 | 16.5451925 | 4.048340169 | b0078 | *ilvH* | acetolactate synthase/acetohydroxybutanoate synthase, regulatory subunit |
| 287.4287558 | 31.25073 | 9.197505765 | 3.2012427 | 159.339742 | 7.315962335 | b0631 | *ybeD* | DUF493 domain-containing protein YbeD |
| 52.39396113 | 5.706655 | 9.181203908 | 3.1986833 | 29.050308 | 4.860481552 | b1172 | *ymgG* | PF13488 family protein YmgG |
| 349.3289802 | 38.24102 | 9.134927681 | 3.1913933 | 193.785002 | 7.598313105 | b2401 | *valU* | tRNA-Val |
| 39.9165987 | 4.380332 | 9.112687462 | 3.1878766 | 22.1484655 | 4.469134842 | b0770 | *ybhI* | putative tricarboxylate transporter |
| 51.76851619 | 5.702688 | 9.077915366 | 3.182361 | 28.7356019 | 4.844767365 | b2971 | *yghG* | lipoprotein YghG |
| 1218.935165 | 134.4604 | 9.065385974 | 3.1803684 | 676.697768 | 9.40236782 | b4164 | *glyX* | tRNA-Gly |
| 195.8985068 | 21.67559 | 9.037747596 | 3.1759633 | 108.787048 | 6.765362994 | b1535 | *dgcZ* | diguanylate cyclase DgcZ |
| 190.8267008 | 21.14948 | 9.022759623 | 3.1735688 | 105.988092 | 6.727758372 | b3261 | *fis* | DNA-binding transcriptional dual regulator Fis |
| 36.65605168 | 4.070473 | 9.005354239 | 3.170783 | 20.3632624 | 4.347896806 | b2985 | *yghS* | putative ATP-binding protein YghS |
| 488.0657971 | 54.5104 | 8.953626621 | 3.1624722 | 271.2881 | 8.083681956 | b1838 | *pphA* | phosphoprotein phosphatase 1 |
| 63.01167674 | 7.040678 | 8.949660389 | 3.1618329 | 35.0261774 | 5.13036164 | b4097 | *phnK* | carbon-phosphorus lyase subunit PhnK |
| 96.25509972 | 10.76706 | 8.939777434 | 3.1602389 | 53.5110787 | 5.741765707 | b0787 | *ybhM* | Bax1-I family protein YbhM |
| 232.3354289 | 26.04227 | 8.921472482 | 3.1572818 | 129.188851 | 7.013337764 | b1839 | *yebY* | DUF2511 domain-containing protein YebY |
| 40.61174071 | 4.595695 | 8.836908764 | 3.1435418 | 22.603718 | 4.498488193 | b0588 | *fepC* | ferric enterobactin ABC transporter ATP binding subunit |
| 582.6278853 | 66.2022 | 8.800732823 | 3.1376237 | 324.415043 | 8.341696909 | b0180 | *fabZ* | 3-hydroxy-acyl-[acyl-carrier-protein] dehydratase |
| 121.9407072 | 13.87395 | 8.789184686 | 3.1357293 | 67.9073285 | 6.085495372 | b2740 | *ygbN* | putative transporter YgbN |
| 219.6725975 | 25.00058 | 8.786699054 | 3.1353213 | 122.33659 | 6.93471216 | b1564 | *relB* | Qin prophage; antitoxin/DNA-binding transcriptional repressor RelB |
| 68.80750142 | 7.870554 | 8.742396368 | 3.1280288 | 38.3390276 | 5.260741842 | b3635 | *mutM* | DNA-formamidopyrimidine glycosylase |
| 39.33002836 | 4.50338 | 8.733446331 | 3.1265511 | 21.9167042 | 4.453958961 | b0572 | *cusC* | copper/silver export system outer membrane channel |

| | | | | | | | | |
|---|---|---|---|---|---|---|---|---|
| 30.32057936 | 3.472303 | 8.732123284 | 3.1263325 | 16.8964413 | 4.078647512 | b0583 | *entD* | phosphopantetheinyl transferase EntD |
| 99.03980802 | 11.46816 | 8.636069926 | 3.1103749 | 55.2539826 | 5.788006551 | b1282 | *yciH* | putative translation factor |
| 31.81765796 | 3.689322 | 8.624255877 | 3.1084 | 17.75349 | 4.150030757 | b0319 | *yahE* | DUF2877 domain-containing protein YahE |
| 44.47814122 | 5.167443 | 8.607378665 | 3.1055739 | 24.8227923 | 4.633593505 | b1168 | *pdeG* | putative c-di-GMP phosphodiesterase PdeG |
| 168.2210422 | 19.62255 | 8.572842886 | 3.0997737 | 93.9217963 | 6.553388096 | b0199 | *metN* | L-methionine/D-methionine ABC transporter ATP binding subunit |
| 126.8485375 | 14.80298 | 8.569123652 | 3.0991477 | 70.8257571 | 6.146202213 | b3658 | *selC* | tRNA-Sec |
| 279.1013461 | 32.69831 | 8.53565051 | 3.0935011 | 155.899828 | 7.284475526 | b1216 | *chaA* | Na(+)/K(+):H(+) antiporter ChaA |
| 12111.89328 | 1420.569 | 8.526086886 | 3.0918838 | 6766.23106 | 12.72413673 | b3699 | *gyrB* | DNA gyrase subunit B |
| 1498.398349 | 176.9778 | 8.466588828 | 3.0817808 | 837.68808 | 9.710269334 | b2185 | *rplY* | 50S ribosomal subunit protein L25 |
| 1465.476853 | 173.8388 | 8.430090788 | 3.0755482 | 819.657826 | 9.678877958 | b3163 | *nlpI* | lipoprotein NlpI |
| 117.4934578 | 13.93782 | 8.42982735 | 3.0755031 | 65.7156414 | 6.038164891 | b1453 | *ansP* | L-asparagine transporter |
| 139.6169861 | 16.58202 | 8.419781949 | 3.0737829 | 78.0995027 | 6.287241456 | b2457 | *eutM* | putative structural protein, ethanolamine utilization microcompartment |
| 29.35518635 | 3.547049 | 8.275945119 | 3.0489241 | 16.4511179 | 4.040113716 | b4087 | *alsA* | D-allose ABC transporter ATP binding subunit |
| 72.26723575 | 8.773864 | 8.236648871 | 3.0420575 | 40.5205498 | 5.340581844 | b1987 | *cbl* | DNA-binding transcriptional activator Cbl |
| 30.44966697 | 3.712958 | 8.200919117 | 3.0357856 | 17.0813124 | 4.094346921 | b0137 | *yadL* | putative fimbrial protein YadL |
| 72.94148023 | 9.009219 | 8.096315556 | 3.0172655 | 40.9753496 | 5.356684354 | b3471 | *yhhQ* | putative queuosine precursor transporter |
| 215.1894832 | 26.78634 | 8.03355342 | 3.0060383 | 120.987911 | 6.918719091 | b2815 | *metW* | tRNA-Met |
| 249.1993153 | 31.09678 | 8.013668385 | 3.0024628 | 140.14805 | 7.130807858 | b3937 | *yiiX* | putative lipid binding hydrolase |
| 136.3485088 | 17.04585 | 7.998926035 | 2.9998063 | 76.6971804 | 6.261101635 | b0283 | *paoD* | molybdenum cofactor insertion chaperone for PaoABC |
| 496463.3736 | 62568.48 | 7.934719854 | 2.9881793 | 279515.928 | 18.09257097 | b0745 | *lysW* | tRNA-Lys |
| 53.61923381 | 6.760643 | 7.931084626 | 2.9875182 | 30.1899385 | 4.915995914 | b1025 | *dgcT* | putative diguanylate cyclase DgcT |
| 37.35199566 | 4.72572 | 7.903979978 | 2.9825793 | 21.0388578 | 4.394984477 | b2122 | *yehQ* | SWIM zinc finger domains-containing protein YehQ |
| 116.4849851 | 14.83085 | 7.854233031 | 2.9734704 | 65.6579197 | 6.036897134 | b0456 | *ybaA* | DUF1428 domain-containing protein YbaA |
| 70.30807284 | 8.951615 | 7.854233031 | 2.9734704 | 39.6298441 | 5.308515386 | b2981 | *yghO* | putative DNA-binding transcriptional regulator YghO |
| 2040.674403 | 260.5935 | 7.830872028 | 2.969173 | 1150.63395 | 10.16821323 | b2608 | *rimM* | ribosome maturation factor RimM |
| 1320.304149 | 170.7495 | 7.732405887 | 2.9509174 | 745.526806 | 9.542116416 | b4039 | *ubiC* | chorismate lyase |
| 217.5284993 | 28.14507 | 7.728832427 | 2.9502505 | 122.836783 | 6.940598821 | b3184 | *yhbE* | inner membrane protein YhbE |
| 474.0813437 | 61.38801 | 7.722702502 | 2.9491058 | 267.734677 | 8.0646602 | b0094 | *ftsA* | cell division protein FtsA |
| 211.3041175 | 27.60481 | 7.65461222 | 2.9363293 | 119.454464 | 6.900316956 | b1252 | *tonB* | Ton complex subunit TonB |
| 27146.0147 | 3549.751 | 7.647301894 | 2.9349508 | 15347.8827 | 13.90575202 | b1095 | *fabF* | beta-ketoacyl-[acyl carrier protein] synthase II |
| 38.66615714 | 5.058148 | 7.64433087 | 2.9343902 | 21.8621526 | 4.450363553 | b0124 | *gcd* | quinoprotein glucose dehydrogenase |
| 508.0713361 | 67.03281 | 7.579442301 | 2.9220917 | 287.552074 | 8.167679435 | b0998 | *torD* | trimethylamine-N-oxide reductase-specific chaperone |
| 1123.138092 | 148.3039 | 7.573220415 | 2.9209069 | 635.720994 | 9.312249922 | b2617 | *bamE* | outer membrane protein assembly factor BamE |

| | | | | | | | | |
|---|---|---|---|---|---|---|---|---|
| 42.07853044 | 5.572929 | 7.55052331 | 2.9165766 | 23.8257297 | 4.574448496 | b0834 | *dgcI* | putative diguanylate cyclase DgcI |
| 58.70663658 | 7.842388 | 7.485811141 | 2.9041587 | 33.2745124 | 5.056345619 | b2104 | *thiM* | hydroxyethylthiazole kinase |
| 69.10672899 | 9.308728 | 7.423864099 | 2.8921703 | 39.2077283 | 5.29306615 | b2825 | *ppdB* | conserved protein PpdB |
| 40.71152385 | 5.489993 | 7.415587773 | 2.890561 | 23.1007584 | 4.529868308 | b4504 | *ykfH* | DUF987 domain-containing protein YkfH |
| 52.47880779 | 7.090501 | 7.401283011 | 2.8877754 | 29.7846546 | 4.896497324 | b3454 | *livF* | branched chain amino acid/phenylalanine ABC transporter ATP binding subunit LivF |
| 28.69631556 | 3.89679 | 7.364090636 | 2.8805074 | 16.2965528 | 4.026494922 | b2452 | *eutH* | putative inner membrane protein |
| 84.98728617 | 11.55183 | 7.35704366 | 2.8791262 | 48.2695559 | 5.59304165 | b3991 | *thiG* | 1-deoxy-D-xylulose 5-phosphate:thiol sulfurtransferase |
| 345.199796 | 47.21577 | 7.311111566 | 2.8700908 | 196.207785 | 7.616238477 | b0357 | *frmR* | DNA-binding transcriptional repressor FrmR |
| 33.53545255 | 4.595695 | 7.297144357 | 2.867332 | 19.065574 | 4.252898058 | b0342 | *lacA* | galactoside O-acetyltransferase |
| 74.53177413 | 10.25415 | 7.268453094 | 2.8616484 | 42.3929597 | 5.405752788 | b1705 | *ydiE* | PF10636 family protein YdiE |
| 46.32551375 | 6.3777 | 7.263671217 | 2.8606989 | 26.3516067 | 4.719819024 | b2577 | *yfiE* | putative LysR-type DNA-binding transcriptional regulator YfiE |
| 31.10408459 | 4.286604 | 7.256112766 | 2.8591969 | 17.6953445 | 4.145297942 | b4366 | *bglJ* | DNA-binding transcriptional regulator BglJ |
| 64.68493394 | 8.92878 | 7.244543709 | 2.8568948 | 36.8068568 | 5.201902646 | b3191 | *mlaB* | intermembrane phospholipid transport system protein MlaB |
| 49.11933856 | 6.793637 | 7.230198078 | 2.8540352 | 27.9564876 | 4.805111209 | b2100 | *yegV* | putative sugar kinase YegV |
| 110.1268709 | 15.26367 | 7.21496859 | 2.8509931 | 62.6952685 | 5.970284663 | b0400 | *phoR* | sensory histidine kinase PhoR |
| 1312.143492 | 182.2747 | 7.198715997 | 2.8477396 | 747.209075 | 9.545368166 | b3006 | *exbB* | Ton complex subunit ExbB |
| 198.6703078 | 27.61692 | 7.193788699 | 2.8467518 | 113.143615 | 6.822011366 | b3939 | *metB* | O-succinylhomoserine(thiol)-lyase/O- succinylhomoserine lyase |
| 34.3027786 | 4.773381 | 7.186264583 | 2.8452421 | 19.5380798 | 4.28821678 | b2406 | *xapB* | xanthosine:H(+) symporter XapB |
| 182.2136874 | 25.39742 | 7.174496683 | 2.8428776 | 103.805552 | 6.697739804 | b3162 | *deaD* | ATP-dependent RNA helicase DeaD |
| 47.76349379 | 6.682546 | 7.147499735 | 2.8374387 | 27.2230198 | 4.766755204 | b2619 | *ratA* | ribosome association toxin RatA |
| 251.5019094 | 35.37293 | 7.110011059 | 2.8298518 | 143.437419 | 7.164277621 | b3647 | *ligB* | DNA ligase B |
| 17939.40576 | 2525.17 | 7.104235511 | 2.8286794 | 10232.2881 | 13.32084117 | b4480 | *hdfR* | DNA-binding transcriptional dual regulator HdfR |
| 70.12871551 | 9.885435 | 7.094145963 | 2.826629 | 40.007075 | 5.32218325 | b0293 | *ecpA* | common pilus major subunit |
| 254.25411 | 36.03025 | 7.056684255 | 2.8189905 | 145.142181 | 7.181323043 | b2980 | *glcC* | DNA-binding transcriptional dual regulator GlcC |
| 954.949234 | 136.202 | 7.011272737 | 2.8096764 | 545.575607 | 9.091635333 | b3401 | *hslO* | molecular chaperone Hsp33 |
| 37.59444017 | 5.393527 | 6.970288996 | 2.8012185 | 21.4939835 | 4.425860976 | b1008 | *rutE* | putative malonic semialdehyde reductase |
| 118.9205039 | 17.27014 | 6.885902933 | 2.7836458 | 68.0953217 | 6.08948378 | b2402 | *valX* | tRNA-Val |
| 865.881492 | 125.8001 | 6.882993805 | 2.7830362 | 495.840809 | 8.953733205 | b3601 | *mtlR* | transcriptional repressor MtlR |
| 95.05501629 | 13.81373 | 6.881199446 | 2.7826601 | 54.4343721 | 5.76644601 | b0597 | *entH* | proofreading thioesterase in enterobactin biosynthesis |
| 112.2771054 | 16.38927 | 6.850646723 | 2.7762402 | 64.3331882 | 6.007491281 | b4296 | *yjhF* | KpLE2 phage-like element; putative transporter YjhF |
| 382.7936925 | 56.10199 | 6.823174887 | 2.7704432 | 219.447841 | 7.77773427 | b4110 | *yjcZ* | uncharacterized protein YjcZ |
| 179.6981622 | 26.41431 | 6.803061963 | 2.7661842 | 103.056234 | 6.687287971 | b3156 | *yhbS* | putative acyltransferase with acyl-CoA N-acyltransferase domain |
| 1007.355768 | 148.441 | 6.786238546 | 2.7626121 | 577.898364 | 9.174671976 | b3302 | *rpmD* | 50S ribosomal subunit protein L30 |

| | | | | | | | | |
|---|---|---|---|---|---|---|---|---|
| 36.31322529 | 5.357268 | 6.778310698 | 2.7609258 | 20.8352465 | 4.380954265 | b3322 | *gspB* | putative general secretion pathway protein B |
| 173.6746172 | 25.68553 | 6.76157413 | 2.7573592 | 99.6800738 | 6.639233231 | b0460 | *hha* | hemolysin expression modulating protein |
| 87.97519306 | 13.01563 | 6.759195328 | 2.7568515 | 50.4954121 | 5.658080408 | b1297 | *puuA* | glutamate-putrescine ligase |
| 52.77848121 | 7.812682 | 6.755488104 | 2.75606 | 30.2955817 | 4.9210355 | b0467 | *priC* | primosomal replication protein N'' |
| 186.2530186 | 27.59805 | 6.748775574 | 2.7546258 | 106.925532 | 6.740462579 | b3334 | *gspM* | Type II secretion system protein GspM |
| 109.8870783 | 16.28274 | 6.74868385 | 2.7546062 | 63.0849102 | 5.979223051 | b0807 | *rlmF* | 23S rRNA m(6)A1618 methyltransferase |
| 46.85970081 | 6.999598 | 6.69462785 | 2.7430039 | 26.9296492 | 4.751123531 | b1022 | *pgaC* | poly-N-acetyl-D-glucosamine synthase subunit PgaC |
| 244.7780372 | 36.87586 | 6.637893717 | 2.7307255 | 140.826948 | 7.137779622 | b1712 | *ihfA* | integration host factor subunit alpha |
| 413.6173063 | 63.06965 | 6.558103499 | 2.7132787 | 238.343479 | 7.896898345 | b0403 | *malZ* | maltodextrin glucosidase |
| 70.01768361 | 10.67864 | 6.556797274 | 2.7129913 | 40.3481623 | 5.334431059 | b3641 | *slmA* | nucleoid occlusion factor SlmA |
| 223.3216439 | 34.15777 | 6.53794507 | 2.7088373 | 128.739708 | 7.008313297 | b0986 | *gfcB* | lipoprotein GfcB |
| 203.4090371 | 31.25073 | 6.508937445 | 2.702422 | 117.329883 | 6.874426691 | b0842 | *mdfA* | multidrug efflux pump MdfA/Na(+):H(+) antiporter/K(+):H(+) antiporter |
| 289.1608681 | 44.6439 | 6.477052446 | 2.6953374 | 166.902383 | 7.382860743 | b3636 | *rpmG* | 50S ribosomal subunit protein L33 |
| 1258.108758 | 195.2729 | 6.442822678 | 2.6876929 | 726.690836 | 9.505197903 | b2684 | *mprA* | DNA-binding transcriptional repressor MprA |
| 346.4777878 | 54.29991 | 6.380817169 | 2.6737412 | 200.388848 | 7.646658413 | b2664 | *csiR* | DNA-binding transcriptional repressor CsiR |
| 4730.483879 | 742.3118 | 6.372637098 | 2.6718905 | 2736.39785 | 11.41806229 | b2082 | *ogrK* | prophage P2 late control protein OgrK |
| 37.27672332 | 5.870303 | 6.350051416 | 2.6667683 | 21.573513 | 4.431189214 | b3376 | *yhfS* | putative aminotransferase YhfS |
| 259.5778488 | 40.89602 | 6.347265083 | 2.6661351 | 150.236932 | 7.231095696 | b0783 | *moaC* | cyclic pyranopterin monophosphate synthase |
| 94.53931271 | 14.90621 | 6.342279017 | 2.6650013 | 54.722759 | 5.774069064 | b3219 | *yhcF* | DUF1120 domain-containing protein YhcF |
| 144.4118213 | 22.8158 | 6.32946633 | 2.6620839 | 83.6138093 | 6.385669327 | b0804 | *ybiX* | PKHD-type hydroxylase YbiX |
| 103.9107323 | 16.42788 | 6.325269071 | 2.6611269 | 60.1693039 | 5.91095576 | b0589 | *fepG* | ferric enterobactin ABC transporter membrane subunit FepG |
| 110.3378343 | 17.48043 | 6.312077687 | 2.658115 | 63.9091323 | 5.997950195 | b4304 | *sgcC* | putative PTS enzyme IIC component SgcC |
| 1718.31812 | 273.3892 | 6.285244133 | 2.6519688 | 995.85368 | 9.959789974 | b3983 | *rplK* | 50S ribosomal subunit protein L11 |
| 908.8169175 | 145.0034 | 6.267556763 | 2.6479032 | 526.910149 | 9.041413158 | b2348 | *argW* | tRNA-Arg |
| 40.82680729 | 6.52321 | 6.258698594 | 2.6458627 | 23.6750088 | 4.565293057 | b3944 | *yijF* | conserved protein YijF |
| 69.79520583 | 11.15171 | 6.258698593 | 2.6458627 | 40.473459 | 5.338904248 | b0730 | *mngR* | DNA-binding transcriptional repressor MngR |
| 3917.084908 | 627.1748 | 6.245603558 | 2.642841 | 2272.12983 | 11.14982956 | b2572 | *rseA* | anti-sigma factor |
| 331.63476 | 53.30769 | 6.221142446 | 2.6371795 | 192.471227 | 7.588498981 | b4291 | *fecA* | ferric citrate outer membrane transporter |
| 70.74509744 | 11.38032 | 6.216440145 | 2.6360887 | 41.0627103 | 5.359756948 | b4049 | *dusA* | tRNA-dihydrouridine synthase A |
| 53.93194834 | 8.677501 | 6.215147747 | 2.6357887 | 31.3047245 | 4.968308498 | b3422 | *rtcR* | DNA-binding transcriptional activator RtcR |
| 178.0576023 | 28.70311 | 6.203424388 | 2.6330648 | 103.380359 | 6.691818301 | b3460 | *livJ* | branched chain amino acid/phenylalanine ABC transporter periplasmic binding protein |
| 35.44679959 | 5.748191 | 6.166600865 | 2.6244755 | 20.5974953 | 4.364397008 | b2349 | *intS* | CPS-53 (KpLE1) prophage; prophage CPS-53 integrase |
| 28.78330667 | 4.677657 | 6.153360065 | 2.6213744 | 16.7304817 | 4.064407082 | b2162 | *rihB* | pyrimidine-specific ribonucleoside hydrolase RihB |

| | | | | | | | | |
|---|---|---|---|---|---|---|---|---|
| 184.9582173 | 30.14451 | 6.135718553 | 2.6172323 | 107.551363 | 6.748881994 | b2107 | *rcnB* | periplasmic protein involved in nickel/cobalt export |
| 137.9339565 | 22.52959 | 6.122345145 | 2.6140844 | 80.2317757 | 6.326101823 | b0849 | *grxA* | reduced glutaredoxin 1 |
| 2681.088511 | 439.0078 | 6.107154082 | 2.6105002 | 1560.04817 | 10.60737487 | b3722 | *bglF* | beta-glucoside specific PTS enzyme II/BglG kinase/BglG phosphatase |
| 99.95684328 | 16.39699 | 6.096049361 | 2.6078746 | 58.1769152 | 5.862374896 | b1871 | *cmoB* | tRNA U34 carboxymethyltransferase |
| 272.2322987 | 44.67129 | 6.094122605 | 2.6074185 | 158.451793 | 7.307900172 | b3842 | *rfaH* | transcription antiterminator RfaH |
| 70.4737828 | 11.58432 | 6.083548449 | 2.6049131 | 41.0290523 | 5.358573925 | b3666 | *uhpT* | hexose-6-phosphate:phosphate antiporter |
| 4517.573992 | 743.4875 | 6.076193753 | 2.6031679 | 2630.53074 | 11.36113819 | b3400 | *hslR* | heat shock protein Hsp15 |
| 117.2202516 | 19.30192 | 6.072983838 | 2.6024055 | 68.2610861 | 6.092991464 | b1193 | *emtA* | lytic murein transglycosylase E |
| 42.83501118 | 7.071333 | 6.057558425 | 2.5987364 | 24.953172 | 4.641151312 | b2633 | *yfjQ* | CP4-57 prophage; protein YfjQ |
| 107.5947416 | 17.85756 | 6.025165065 | 2.5910008 | 62.7261504 | 5.970995119 | b3535 | *yhjR* | PF10945 family protein YhjR |
| 588.782336 | 97.84454 | 6.017528607 | 2.5891711 | 343.313439 | 8.423382524 | b4537 | *yecJ* | DUF2766 domain-containing protein YecJ |
| 195.2310035 | 32.45268 | 6.015866971 | 2.5887727 | 113.841842 | 6.830887095 | b3740 | *rsmG* | 16S rRNA m(7)G527 methyltransferase |
| 3178.076205 | 529.8089 | 5.998533487 | 2.5846098 | 1853.94253 | 10.85638081 | b2529 | *iscU* | scaffold protein for iron-sulfur cluster assembly |
| 6203.135728 | 1034.939 | 5.993719658 | 2.5834516 | 3619.03749 | 11.82139034 | b2340 | *sixA* | phosphohistidine phosphatase |
| 1389.625315 | 232.4273 | 5.978752728 | 2.5798445 | 811.026304 | 9.663604896 | b2607 | *trmD* | tRNA m(1)G37 methyltransferase |
| 59.17026344 | 9.90004 | 5.976770165 | 2.5793661 | 34.5351517 | 5.109993656 | b1670 | *ydhU* | putative cytochrome YdhU |
| 144.8856099 | 24.24755 | 5.975267219 | 2.5790032 | 84.5665816 | 6.402015757 | b3161 | *mtr* | tryptophan:H(+) symporter Mtr |
| 1117.751742 | 187.1063 | 5.973887064 | 2.57867 | 652.429007 | 9.349677116 | b0458 | *ylaC* | putative inner membrane protein |
| 144.7335459 | 24.2492 | 5.968590747 | 2.5773903 | 84.4913725 | 6.40073213 | b2210 | *mqo* | malate:quinone oxidoreductase |
| 273.0484789 | 45.85221 | 5.954968968 | 2.574094 | 159.450344 | 7.316963397 | b2437 | *eutR* | putative AraC-type transcriptional regulator EutR |
| 214.4422975 | 36.13365 | 5.93469712 | 2.5691744 | 125.287976 | 6.969104156 | b1825 | *yebO* | uncharacterized protein YebO |
| 127.2318889 | 21.45112 | 5.931247625 | 2.5683356 | 74.3415031 | 6.216095954 | b1581 | *rspA* | mandelate racemase/muconate lactonizing enzyme family protein RspA |
| 374.9078996 | 63.33481 | 5.919460414 | 2.5654657 | 219.121355 | 7.77558628 | b4203 | *rplI* | 50S ribosomal subunit protein L9 |
| 58.96932627 | 9.997084 | 5.898652413 | 2.5603854 | 34.4832054 | 5.10782198 | b1528 | *ydeA* | L-arabinose exporter |
| 458.1742747 | 77.84272 | 5.88589726 | 2.5572624 | 268.008499 | 8.066134943 | b0429 | *cyoD* | cytochrome bo3 ubiquinol oxidase subunit 4 |
| 1184.563627 | 201.3496 | 5.883117952 | 2.556581 | 692.956629 | 9.436621249 | b3637 | *rpmB* | 50S ribosomal subunit protein L28 |
| 38.29643345 | 6.532638 | 5.862322765 | 2.5514724 | 22.4145358 | 4.486362717 | b1974 | *yodB* | putative cytochrome |
| 1998.836038 | 344.861 | 5.796063196 | 2.5350733 | 1171.84851 | 10.19457036 | b1060 | *bssS* | regulator of biofilm formation |
| 257.2211263 | 44.4197 | 5.790698836 | 2.5337375 | 150.820415 | 7.236687914 | b1828 | *yebQ* | putative transporter YebQ |
| 27.38611985 | 4.739023 | 5.778853285 | 2.5307832 | 16.0625715 | 4.005630974 | b4029 | *yjbH* | YjbH family protein |
| 247.9676627 | 43.03379 | 5.762161828 | 2.5266102 | 145.500726 | 7.184882546 | b1957 | *yodC* | protein YodC |
| 282.9450908 | 49.4944 | 5.716709516 | 2.515185 | 166.219744 | 7.376947948 | b0356 | *frmA* | S-(hydroxymethyl)glutathione dehydrogenase |
| 375.6034616 | 65.82946 | 5.705705018 | 2.5124052 | 220.716459 | 7.786050407 | b2942 | *metK* | methionine adenosyltransferase |

| | | | | | | | | |
|---|---|---|---|---|---|---|---|---|
| 18660.49987 | 3283.393 | 5.683298284 | 2.5067284 | 10971.9463 | 13.42153184 | b3758 | *rrlC* | 23S ribosomal RNA |
| 128.718568 | 22.70029 | 5.670349789 | 2.5034377 | 75.7094275 | 6.242401054 | b2392 | *mntH* | Mn(2(+))/Fe(2(+)): H(+) symporter MntH |
| 162.4975378 | 28.68218 | 5.665453717 | 2.5021915 | 95.5898567 | 6.578785632 | b1532 | *marB* | multiple antibiotic resistance protein |
| 156.4751484 | 27.65041 | 5.659052603 | 2.5005605 | 92.0627813 | 6.524546123 | b2237 | *inaA* | putative lipopolysaccharide kinase InaA |
| 50.7351945 | 8.983625 | 5.647519392 | 2.4976173 | 29.8594096 | 4.900113734 | b0377 | *sbmA* | peptide antibiotic/peptide nucleic acid transporter |
| 111.5610357 | 19.77258 | 5.642209659 | 2.4962603 | 65.6668073 | 6.037092409 | b0818 | *ybiR* | putative transporter YbiR |
| 128.7982656 | 22.90363 | 5.623487395 | 2.4914651 | 75.850947 | 6.245095288 | b0831 | *gsiC* | glutathione ABC transporter membrane subunit GsiC |
| 18661.30785 | 3356.248 | 5.560169337 | 2.4751288 | 11008.778 | 13.42636671 | b0441 | *ppiD* | periplasmic folding chaperone |
| 73.37153154 | 13.20453 | 5.556541117 | 2.4741871 | 43.2880324 | 5.43589632 | b0989 | *cspH* | CspA family protein CspH |
| 65.55309257 | 11.80583 | 5.552603099 | 2.4731643 | 38.6794617 | 5.273495811 | b1949 | *fliQ* | flagellar biosynthesis protein FliQ |
| 118.3011263 | 21.58767 | 5.480031084 | 2.4541841 | 69.9444003 | 6.128136656 | b4440 | *ryfA* | small regulatory RNA RyfA |
| 890.7524496 | 162.6667 | 5.475937462 | 2.453106 | 526.709554 | 9.040863819 | b1054 | *lpxL* | lauroyl acyltransferase |
| 36.98920584 | 6.759562 | 5.472130591 | 2.4521027 | 21.8743839 | 4.451170479 | b0592 | *fepB* | ferric enterobactin ABC transporter periplasmic binding protein |
| 2998.795866 | 558.4546 | 5.369811597 | 2.4248715 | 1778.62522 | 10.79654683 | b1842 | *holE* | DNA polymerase III subunit theta |
| 1243.678709 | 232.6885 | 5.344821361 | 2.4181417 | 738.183627 | 9.527835929 | b3255 | *accB* | biotin carboxyl carrier protein |
| 56.48723934 | 10.59052 | 5.33375268 | 2.4151509 | 33.538882 | 5.067762693 | b2506 | *yfgI* | nalidixic acid resistance protein YfgI |
| 174.5134224 | 32.77515 | 5.324564478 | 2.4126635 | 103.644288 | 6.695496804 | b1684 | *sufA* | iron-sulfur cluster insertion protein SufA |
| 36.71078448 | 6.928228 | 5.298726724 | 2.4056457 | 21.819506 | 4.447546535 | b1972 | *msrQ* | periplasmic protein-L-methionine sulfoxide reductase heme binding subunit |
| 68.35880386 | 12.96057 | 5.274365681 | 2.3989976 | 40.6596887 | 5.345527263 | b0358 | *yaiO* | outer membrane protein YaiO |
| 28.00222121 | 5.332292 | 5.251441682 | 2.3927135 | 16.6672567 | 4.05894476 | b2917 | *scpA* | methylmalonyl-CoA mutase |
| 1235.505059 | 235.6377 | 5.243240698 | 2.3904588 | 735.571368 | 9.522721514 | b1109 | *ndh* | NADH:quinone oxidoreductase II |
| 36.42530315 | 6.975609 | 5.221809724 | 2.3845499 | 21.7004561 | 4.43965346 | b2462 | *eutS* | putative structural protein, ethanolamine utilization microcompartment |
| 72.51864549 | 13.92994 | 5.205954025 | 2.3801626 | 43.2242945 | 5.433770512 | b1486 | *ddpB* | putative D,D-dipeptide ABC transporter membrane subunit DdpB |
| 296.1791131 | 56.95904 | 5.199861767 | 2.3784733 | 176.569075 | 7.464088872 | b3469 | *zntA* | Zn(2(+))/Cd(2(+))/Pb(2(+)) exporting P-type ATPase |
| 33.22778785 | 6.433974 | 5.164427199 | 2.3686083 | 19.8308807 | 4.309676844 | b3329 | *gspH* | Type II secretion system protein GspH |
| 51.81645986 | 10.05353 | 5.154056861 | 2.3657085 | 30.9349944 | 4.951167868 | b2306 | *hisP* | lysine/arginine/ornithine ABC transporter/histidine ABC transporter, ATP binding subunit |
| 167.7212149 | 32.56378 | 5.150544332 | 2.3647249 | 100.1425 | 6.645910561 | b1716 | *rplT* | 50S ribosomal subunit protein L20 |
| 74.47714008 | 14.53725 | 5.123194246 | 2.3570436 | 44.5071935 | 5.475966627 | b2239 | *glpQ* | glycerophosphoryl diester phosphodiesterase |
| 125.1115456 | 24.4211 | 5.12309233 | 2.3570149 | 74.7663224 | 6.224316666 | b0956 | *matP* | macrodomain Ter protein |
| 60.65832131 | 11.85824 | 5.115287469 | 2.3548153 | 36.2582821 | 5.180238668 | b3861 | *yihF* | uncharacterized protein YihF |
| 2164.123093 | 423.4048 | 5.111238772 | 2.353673 | 1293.76395 | 10.33735871 | b3756 | *rrsC* | 16S ribosomal RNA |
| 1103.756734 | 217.5344 | 5.073941849 | 2.343107 | 660.645551 | 9.367732636 | b3294 | *rplQ* | 50S ribosomal subunit protein L17 |
| 160.9582626 | 31.75477 | 5.068789657 | 2.3416413 | 96.3565176 | 6.590310349 | b4594 | *ymgJ* | uncharacterized protein YmgJ |

| | | | | | | | | |
|---|---|---|---|---|---|---|---|---|
| 240.2669035 | 47.46204 | 5.062295752 | 2.3397918 | 143.864474 | 7.168566563 | b3727 | *pstC* | phosphate ABC transporter membrane subunit PstC |
| 721.6543398 | 142.5894 | 5.061067372 | 2.3394417 | 432.121848 | 8.755294365 | b2532 | *trmJ* | tRNA Cm32/Um32 methyltransferase |
| 73.58490949 | 14.54775 | 5.058163264 | 2.3386136 | 44.0663312 | 5.461604884 | b4293 | *fecI* | RNA polymerase sigma factor FecI |
| 52.33996264 | 10.3801 | 5.042336722 | 2.3340925 | 31.3600316 | 4.97085511 | b2492 | *focB* | putative formate transporter |
| 57.25802039 | 11.37636 | 5.033069614 | 2.3314386 | 34.317191 | 5.100859564 | b4330 | *yjiH* | Gate family protein YjiH |
| 502.1087942 | 100.0023 | 5.020970888 | 2.3279664 | 301.055563 | 8.233885965 | b1145 | *ymfK* | e14 prophage; putative repressor protein YmfK |
| 51.04772741 | 10.20857 | 5.000477129 | 2.3220658 | 30.6281494 | 4.936786293 | b4090 | *rpiB* | allose-6-phosphate isomerase/ribose-5-phosphate isomerase B |
| 185.6330558 | 37.15477 | 4.996209934 | 2.3208341 | 111.393915 | 6.799526621 | b0819 | *ldtB* | L,D-transpeptidase YbiS |
| 11375.90237 | 2277.885 | 4.994063221 | 2.3202141 | 6826.89375 | 12.73701358 | b3312 | *rpmC* | 50S ribosomal subunit protein L29 |
| 120.9082324 | 24.21593 | 4.992920772 | 2.319884 | 72.5620825 | 6.181143955 | b2840 | *ygeA* | amino acid racemase |
| 56.78611363 | 11.409 | 4.977310272 | 2.3153663 | 34.0975549 | 5.091596383 | b1952 | *dsrB* | protein DsrB |
| 162.6317817 | 32.74063 | 4.967276774 | 2.3124551 | 97.6862069 | 6.610082966 | b3144 | *yraJ* | putative fimbrial usher protein YraJ |
| 863.2032679 | 174.1112 | 4.957769856 | 2.3096913 | 518.657235 | 9.018637609 | b2814 | *metZ* | tRNA-Met |
| 124.6185941 | 25.26507 | 4.932446882 | 2.3023035 | 74.9418299 | 6.227699299 | b3338 | *chiA* | endochitinase |
| 113.9043098 | 23.14869 | 4.920551471 | 2.29882 | 68.5264988 | 6.098590073 | b4685 | *yrbN* | uncharacterized protein YrbN |
| 1067.557 | 217.3216 | 4.912337742 | 2.2964098 | 642.43929 | 9.327416317 | b2390 | *ypeC* | DUF2502 domain-containing protein YpeC |
| 121.1986745 | 24.78506 | 4.88998904 | 2.2898312 | 72.9918675 | 6.189663828 | b0075 | *leuL* | leu operon leader peptide |
| 522.3601033 | 107.2223 | 4.871747489 | 2.2844394 | 314.791215 | 8.29825147 | b0038 | *caiB* | gamma-butyrobetainyl-CoA:carnitine CoA transferase |
| 59.60568233 | 12.28409 | 4.852266268 | 2.2786587 | 35.9448869 | 5.167714658 | b0629 | *ybeF* | putative LysR-type DNA-binding transcriptional regulator YbeF |
| 298.9944327 | 61.71626 | 4.844661986 | 2.276396 | 180.355348 | 7.494698391 | b0841 | *ybjG* | undecaprenyl pyrophosphate phosphatase |
| 135.3939652 | 28.11201 | 4.816232268 | 2.267905 | 81.7529871 | 6.35319954 | b0495 | *ybbA* | putative ABC transporter ATP-binding protein YbbA |
| 32.0730287 | 6.676918 | 4.803568024 | 2.2641064 | 19.3749732 | 4.276122412 | b2538 | *hcaE* | putative 3-phenylpropionate/cinnamate dioxygenase subunit alpha |
| 1246.18778 | 259.8745 | 4.795344979 | 2.2616346 | 753.03113 | 9.556565696 | b4202 | *rpsR* | 30S ribosomal subunit protein S18 |
| 1000.847731 | 208.7926 | 4.793501489 | 2.2610799 | 604.820175 | 9.240362454 | b0254 | *perR* | putative transcriptional regulator PerR |
| 154.6674411 | 32.36683 | 4.778579191 | 2.2565817 | 93.5171335 | 6.547158804 | b2941 | *yqgD* | DUF2684 domain-containing protein YqgD |
| 23632.47416 | 4946.624 | 4.777495237 | 2.2562544 | 14289.5492 | 13.80267279 | b3315 | *rplV* | 50S ribosomal subunit protein L22 |
| 49.17560355 | 10.30952 | 4.769922343 | 2.2539658 | 29.7425611 | 4.894456978 | b1568 | *ydfX* | Qin prophage; uncharacterized protein YdfX |
| 76.05632663 | 15.96653 | 4.763485203 | 2.2520175 | 46.011428 | 5.523920328 | b0402 | *proY* | putative transporter ProY |
| 87.75956759 | 18.44156 | 4.758792112 | 2.2505954 | 53.1005656 | 5.730655323 | b1330 | *ynaI* | small conductance mechanosensitive channel YnaI |
| 675.6230803 | 141.9759 | 4.75871597 | 2.2505723 | 408.7995 | 8.675249623 | b2592 | *clpB* | ClpB chaperone |
| 43.01774782 | 9.041643 | 4.757735521 | 2.2502751 | 26.0296956 | 4.702086535 | b0522 | *purK* | 5-(carboxyamino)imidazole ribonucleotide synthase |
| 60.4766665 | 12.71538 | 4.756183367 | 2.2498043 | 36.596022 | 5.193614929 | b0709 | *dtpD* | dipeptide:H(+) symporter DtpD |
| 249.1582158 | 52.39926 | 4.754995132 | 2.2494439 | 150.778737 | 7.23628918 | b3744 | *asnA* | asparagine synthetase A |

| | | | | | | | | |
|---|---|---|---|---|---|---|---|---|
| 84.33858652 | 17.7561 | 4.749838459 | 2.2478784 | 51.0473411 | 5.673763914 | b0231 | *dinB* | DNA polymerase IV |
| 50.73119941 | 10.68678 | 4.747099749 | 2.2470464 | 30.7089885 | 4.940589089 | b1628 | *rsxB* | SoxR [2Fe-2S] reducing system protein RsxB |
| 171.7571642 | 36.36594 | 4.723023164 | 2.2397106 | 104.061551 | 6.701293302 | b3183 | *obgE* | GTPase ObgE |
| 1697.791993 | 361.9369 | 4.690850276 | 2.2298495 | 1029.86447 | 10.00823877 | b3851 | *rrsA* | 16S ribosomal RNA |
| 125.1879358 | 26.68981 | 4.690476599 | 2.2297345 | 75.9388736 | 6.246766695 | b1035 | *ycdY* | chaperone protein YcdY |
| 1031.714996 | 221.2999 | 4.662066162 | 2.2209695 | 626.507472 | 9.291187905 | b3067 | *rpoD* | RNA polymerase, sigma 70 (sigma D) factor |
| 38.57510887 | 8.344023 | 4.623082629 | 2.2088551 | 23.4595661 | 4.552104422 | b3662 | *nepI* | purine ribonucleoside exporter |
| 30.78969021 | 6.682546 | 4.607479168 | 2.2039776 | 18.736118 | 4.227750159 | b0247 | *ykfG* | CP4-6 prophage; RadC-like JAB domain-containing protein YkfG |
| 420.9035447 | 91.6688 | 4.591567989 | 2.1989869 | 256.286174 | 8.001611842 | b2530 | *iscS* | cysteine desulfurase |
| 87.01620026 | 19.04238 | 4.569608349 | 2.1920705 | 53.0292881 | 5.728717476 | b4208 | *cycA* | serine/alanine/glycine/cycloserine:H(+)symporter |
| 113.6994078 | 24.98159 | 4.551328744 | 2.1862878 | 69.3404966 | 6.115626264 | b4460 | *araH* | arabinose ABC transporter membrane subunit |
| 1321.024328 | 290.9294 | 4.540704109 | 2.182916 | 805.976865 | 9.654594617 | b3303 | *rpsE* | 30S ribosomal subunit protein S5 |
| 445.837981 | 98.3021 | 4.535386127 | 2.1812254 | 272.070041 | 8.08783429 | b3194 | *mlaE* | intermembrane phospholipid transport system, integral membrane subunit MlaE |
| 109.5345507 | 24.18927 | 4.528229646 | 2.1789471 | 66.8619083 | 6.063112626 | b3071 | *nfeR* | DNA-binding transcriptional repressor NfeR |
| 232.8154024 | 51.6839 | 4.504602305 | 2.1713997 | 142.24965 | 7.152281291 | b1611 | *fumC* | fumarase C |
| 48.56707086 | 10.78894 | 4.501560107 | 2.1704251 | 29.6780064 | 4.891322279 | b4256 | *yjgM* | putative acetyltransferase YjgM |
| 75.31631913 | 16.74146 | 4.498789917 | 2.169537 | 46.0288904 | 5.52446776 | b4459 | *ryjA* | small RNA RyjA |
| 472.5202402 | 105.1612 | 4.493295269 | 2.1677739 | 288.840711 | 8.174130288 | b3741 | *mnmG* | 5-carboxymethylaminomethyluridine-tRNA synthase subunit MnmG |
| 634.5657277 | 142.0577 | 4.466956752 | 2.1592923 | 388.311728 | 8.60107147 | b3816 | *corA* | Ni(2(+))/Co(2(+))/Mg(2(+)) transporter |
| 246.9387512 | 55.32916 | 4.463085233 | 2.1580414 | 151.133955 | 7.239684014 | b2326 | *epmC* | EF-P-Lys34 hydroxylase |
| 654.2936505 | 146.6768 | 4.460784674 | 2.1572975 | 400.485229 | 8.645605221 | b3472 | *dcrB* | periplasmic bacteriophage sensitivity protein DcrB |
| 276.4500731 | 62.14018 | 4.44881374 | 2.1534207 | 169.295125 | 7.40339662 | b2603 | *yfiR* | DUF4154 domain-containing protein YfiR |
| 129.5983509 | 29.22365 | 4.434707234 | 2.1488389 | 79.4110026 | 6.311267005 | b3458 | *livK* | L-leucine/L-phenylalanine ABC transporter periplasmic binding protein |
| 55.50522385 | 12.54281 | 4.425262478 | 2.145763 | 34.0240166 | 5.08848156 | b2354 | *yfdK* | CPS-53 (KpLE1) prophage; putative tail fiber assembly protein YfdK |
| 409.0455263 | 92.45905 | 4.424072251 | 2.1453749 | 250.752289 | 7.970119061 | b4447 | *sibD* | small RNA SibD |
| 354.9535986 | 80.23836 | 4.423739614 | 2.1452665 | 217.595978 | 7.765508079 | b3931 | *hslU* | ATPase component of the HslVU protease |
| 43.16547934 | 9.758133 | 4.423538908 | 2.145201 | 26.461806 | 4.725839622 | b4092 | *phnP* | 5-phospho-alpha-D-ribosyl 1,2-cyclic phosphate phosphodiesterase |
| 1713.470125 | 388.1982 | 4.413905232 | 2.1420557 | 1050.83417 | 10.0373193 | b1288 | *fabI* | enoyl-[acyl-carrier-protein] reductase |
| 59.46025194 | 13.5694 | 4.38193823 | 2.1315691 | 36.5148236 | 5.190410356 | b0749 | *lysQ* | tRNA-Lys |
| 63.86748167 | 14.60461 | 4.373103675 | 2.1286576 | 39.2360468 | 5.294107784 | b2835 | *lplT* | lysophospholipid transporter |
| 139.6490084 | 32.032 | 4.359672283 | 2.1242197 | 85.8405026 | 6.423586618 | b3759 | *rrfC* | 5S ribosomal RNA |
| 431.1881928 | 98.94184 | 4.357996624 | 2.1236651 | 265.065015 | 8.050202456 | b0093 | *ftsQ* | cell division protein FtsQ |
| 50.14139942 | 11.53163 | 4.348160791 | 2.1204053 | 30.8365168 | 4.946567906 | b3173 | *yhbX* | putative hydrolase, inner membrane |

| | | | | | | | | |
|---|---|---|---|---|---|---|---|---|
| 85.776919 | 19.79213 | 4.333890661 | 2.1156628 | 52.7845235 | 5.722043087 | b1533 | *eamA* | cysteine/O-acetylserine exporter EamA |
| 63.24826027 | 14.59976 | 4.332143476 | 2.115081 | 38.9240107 | 5.282588468 | b3959 | *argB* | acetylglutamate kinase |
| 150.9545029 | 34.89665 | 4.325759535 | 2.1129535 | 92.9255749 | 6.538003803 | b3532 | *bcsB* | cellulose synthase periplasmic subunit |
| 28.05902085 | 6.495249 | 4.319929669 | 2.1110078 | 17.2771352 | 4.110792109 | b2408 | *yfeN* | conserved outer membrane protein YfeN |
| 462.6990923 | 107.1454 | 4.318424185 | 2.110505 | 284.922224 | 8.154424345 | b4320 | *fimH* | type 1 fimbriae D-mannose specific adhesin |
| 112.1177649 | 25.965 | 4.318034965 | 2.1103749 | 69.041381 | 6.109389416 | b0945 | *pyrD* | dihydroorotate dehydrogenase, type 2 |
| 997.9412284 | 233.3388 | 4.276791275 | 2.0965288 | 615.640001 | 9.265943162 | b4618 | *tisB* | membrane-depolarizing toxin TisB |
| 104.6059988 | 24.47974 | 4.273166714 | 2.0953056 | 64.5428681 | 6.012185783 | b2960 | *trmI* | tRNA m(7)G46 methyltransferase |
| 196.4773542 | 46.01557 | 4.269802014 | 2.0941692 | 121.24646 | 6.921798814 | b1601 | *tqsA* | autoinducer 2 exporter |
| 28.90242495 | 6.772167 | 4.267825255 | 2.0935011 | 17.8372959 | 4.156825019 | b1006 | *rutG* | pyrimidine:H(+) symporter |
| 76.77333125 | 18.13658 | 4.233064687 | 2.0817025 | 47.4549574 | 5.568486901 | b3494 | *uspB* | putative universal stress (ethanol tolerance) protein B |
| 4957.720846 | 1171.902 | 4.23048983 | 2.0808247 | 3064.81158 | 11.58158267 | b3321 | *rpsJ* | 30S ribosomal subunit protein S10 |
| 131.7163868 | 31.20732 | 4.220688185 | 2.0774782 | 81.4618558 | 6.348052775 | b2836 | *aas* | acyltransferase |
| 227.1011394 | 53.89618 | 4.213677529 | 2.0750799 | 140.498662 | 7.134412578 | b3055 | *ygiM* | putative signal transduction protein (SH3 domain) |
| 139.6282715 | 33.17809 | 4.208448628 | 2.0732885 | 86.4031789 | 6.433012487 | b4356 | *lgoT* | galactonate:H(+) symporter |
| 40.71587337 | 9.695739 | 4.199357469 | 2.0701686 | 25.2058061 | 4.655684189 | b1657 | *ydhP* | putative transporter YdhP |
| 293.3890601 | 69.97712 | 4.19264259 | 2.0678599 | 181.683091 | 7.505280344 | b3070 | *nfeF* | NADPH-dependent ferric chelate reductase |
| 306.8442631 | 73.20773 | 4.191419176 | 2.0674388 | 190.025994 | 7.570052972 | b2586 | *yfiM* | protein YfiM |
| 35.31285309 | 8.426417 | 4.190731606 | 2.0672021 | 21.8696352 | 4.450857248 | b4340 | *yjiR* | fused putative DNA-binding transcriptional regulator/putative aminotransferase YjiR |
| 44.9330313 | 10.72468 | 4.189684316 | 2.0668415 | 27.8288566 | 4.798509727 | b3667 | *uhpC* | inner membrane protein sensing glucose-6-phosphate |
| 93.92594576 | 22.50927 | 4.172768038 | 2.0610047 | 58.2176059 | 5.863383607 | b0738 | *tolR* | Tol-Pal system protein TolR |
| 491.9309132 | 117.9645 | 4.170161977 | 2.0601034 | 304.947688 | 8.252417965 | b3175 | *secG* | Sec translocon subunit SecG |
| 33.98661304 | 8.152364 | 4.168927343 | 2.0596762 | 21.0694885 | 4.397083386 | b1722 | *ydiY* | acid-inducible putative outer membrane protein YdiY |
| 122.7252521 | 29.57658 | 4.149406128 | 2.0529049 | 76.1509172 | 6.250789509 | b1607 | *ydgC* | GlpM family protein |
| 7979.122002 | 1931.343 | 4.131385829 | 2.0466258 | 4955.23237 | 12.274737 | b3306 | *rpsH* | 30S ribosomal subunit protein S8 |
| 82.58368325 | 20.06626 | 4.115549907 | 2.0410852 | 51.3249703 | 5.681588981 | b1477 | *yddM* | putative DNA-binding transcriptional regulator YddM |
| 26.0489448 | 6.331847 | 4.113956792 | 2.0405266 | 16.1903959 | 4.017066357 | b1988 | *nac* | DNA-binding transcriptional dual regulator Nac |
| 476.1510093 | 115.7434 | 4.113848814 | 2.0404888 | 295.947224 | 8.209196115 | b1743 | *spy* | ATP-independent periplasmic chaperone |
| 196.5324826 | 47.98092 | 4.096055199 | 2.0342352 | 122.2567 | 6.933769716 | b3022 | *mqsR* | mRNA interferase/toxin of the MqsR-MqsA toxin-antitoxin system |
| 226.100195 | 55.34768 | 4.08508924 | 2.0303676 | 140.723935 | 7.136723924 | b2783 | *mazE* | antitoxin of the MazF-MazE toxin-antitoxin system MazE |
| 255.0336455 | 62.50146 | 4.0804432 | 2.0287259 | 158.767551 | 7.310772276 | b0775 | *bioB* | biotin synthase |
| 79.17238339 | 19.40405 | 4.0801995 | 2.0286397 | 49.2882153 | 5.623170836 | b0248 | *yafX* | CP4-6 prophage; protein YafX |
| 604.099643 | 148.2549 | 4.074735219 | 2.0267063 | 376.177294 | 8.555268959 | b4319 | *fimG* | type 1 fimbriae minor subunit FimG |

| | | | | | | | | |
|---|---|---|---|---|---|---|---|---|
| 49.83617136 | 12.31372 | 4.047206836 | 2.0169266 | 31.0749456 | 4.957679962 | b1971 | *msrP* | periplasmic protein-L-methionine sulfoxide reductase catalytic subunit |
| 110.4900862 | 27.34439 | 4.040686094 | 2.0146003 | 68.9172368 | 6.106792954 | b0253 | *ykfA* | CP4-6 prophage; putative GTP-binding protein YkfA |
| 387.7893812 | 96.35641 | 4.024531105 | 2.0088207 | 242.072897 | 7.919297751 | b2017 | *yefM* | YefM antitoxin of the YoeB-YefM toxin-antitoxin pair and DNA binding transcriptional repressor |
| 87.62290722 | 21.81423 | 4.016776711 | 2.0060383 | 54.7185706 | 5.773958639 | b0065 | *yabI* | DedA family protein YabI |
| 66.17466445 | 16.47457 | 4.016776709 | 2.0060383 | 41.3246167 | 5.368929531 | b3603 | *lldP* | lactate/glycolate:H(+) symporter LldP |

**Table S2.** List of genes, extracted from Dataset 1, which were downregulated in the presence of TAT-RasGAP$_{317\text{-}326}$

| WT_TAT-RasGAP | WT_untreated | FoldChange | $Log_2$Fold Change | Mean_WT_untr_TAT | $Log_2$_mean_WT_untr_TAT | Locus_tag | Gene | Product |
|---|---|---|---|---|---|---|---|---|
| 0 | 49.9555 | 0 | | 24.9777721 | 4.642572895 | b0031 | *dapB* | 4-hydroxy-tetrahydrodipicolinate reductase |
| 0 | 44.0351 | 0 | | 22.0175587 | 4.46058261 | b0101 | *yacG* | DNA gyrase inhibitor YacG |
| 0 | 269.628 | 0 | | 134.814189 | 7.07482854 | b0163 | *yaeH* | DUF3461 domain-containing protein YaeH |
| 0 | 55.0013 | 0 | | 27.5006411 | 4.781393347 | b0668 | *glnW* | tRNA-Gln |
| 0 | 61.6791 | 0 | | 30.8395347 | 4.946709095 | b0746 | *valZ* | tRNA-Val |
| 0 | 200.703 | 0 | | 100.351718 | 6.64892151 | b0798 | *ybiA* | N-glycosidase YbiA |
| 0 | 3307.45 | 0 | | 1653.72508 | 10.6915037 | b0836 | *bssR* | regulator of biofilm formation |
| 0 | 60.8213 | 0 | | 30.4106552 | 4.926504994 | b0968 | *yccX* | acylphosphatase |
| 0 | 77.1151 | 0 | | 38.5575536 | 5.268941611 | b1073 | *flgB* | flagellar basal-body rod protein FlgB |
| 0 | 90.3964 | 0 | | 45.1982013 | 5.498193457 | b1139 | *lit* | e14 prophage; cell death peptidase Lit |
| 0 | 88.4167 | 0 | | 44.2083477 | 5.466246908 | b1295 | *ymjA* | DUF2543 domain-containing protein YmjA |
| 0 | 37.7228 | 0 | | 18.8613865 | 4.237363823 | b1441 | *ydcT* | putative ABC transporter ATP-binding protein YdcT |
| 0 | 44.7013 | 0 | | 22.3506281 | 4.482243471 | b1493 | *gadB* | glutamate decarboxylase B |
| 0 | 35.7674 | 0 | | 17.8837177 | 4.160574772 | b1513 | *lsrA* | Autoinducer-2 ABC transporter ATP binding subunit |
| 0 | 73.4948 | 0 | | 36.7473982 | 5.199570201 | b1519 | *tam* | trans-aconitate 2-methyltransferase |
| 0 | 43.9937 | 0 | | 21.996872 | 4.459226481 | b1597 | *asr* | acid shock protein |
| 0 | 186.267 | 0 | | 93.1333593 | 6.541226112 | b1635 | *gstA* | glutathione S-transferase GstA |
| 0 | 120.539 | 0 | | 60.2692622 | 5.913350498 | b1665 | *valV* | tRNA-Val |
| 0 | 41.2689 | 0 | | 20.6344523 | 4.366983243 | b1674 | *ydhY* | putative 4Fe-4S ferredoxin-type protein |
| 0 | 37.9473 | 0 | | 18.9736566 | 4.245925837 | b1675 | *fumD* | fumarase D |
| 0 | 99.5991 | 0 | | 49.7995481 | 5.638060744 | b1708 | *nlpC* | NlpC/P60 family lipoprotein NlpC |
| 0 | 118.753 | 0 | | 59.3763842 | 5.891817336 | b1715 | *pheM* | pheST-ihfA operon leader peptide |
| 0 | 371.787 | 0 | | 185.893509 | 7.538332584 | b1724 | *ydiZ* | protein YdiZ |
| 0 | 225.4 | 0 | | 112.699765 | 6.816340699 | b1784 | *yeaH* | DUF444 domain-containing protein YeaH |
| 0 | 71.4302 | 0 | | 35.7151183 | 5.158462996 | b1788 | *yoaI* | protein YoaI |
| 0 | 64.4794 | 0 | | 32.2396756 | 5.010765324 | b1836 | *yebV* | protein YebV |
| 0 | 130.029 | 0 | | 65.0146276 | 6.022692439 | b1895 | *uspC* | universal stress protein C |
| 0 | 51.5637 | 0 | | 25.781851 | 4.688283942 | b1906 | *yecH* | DUF2492 domain-containing protein YecH |
| 0 | 149.263 | 0 | | 74.6316741 | 6.221716142 | b1953 | *yodD* | stress-induced protein |
| 0 | 135.779 | 0 | | 67.8895137 | 6.085116846 | b1954 | *dsrA* | small regulatory RNA DsrA |
| 0 | 65.1424 | 0 | | 32.5711819 | 5.025524164 | b1967 | *hchA* | protein/nucleic acid deglycase 1 |
| 0 | 42.2804 | 0 | | 21.1401987 | 4.401917033 | b2018 | *hisL* | his operon leader peptide |
| 0 | 42.1281 | 0 | | 21.064027 | 4.396709368 | b2111 | *yehD* | putative fimbrial protein YehD |
| 0 | 42.5542 | 0 | | 21.2770918 | 4.411229066 | b2112 | *yehE* | DUF2574 domain-containing protein YehE |
| 0 | 32.4226 | 0 | | 16.2113154 | 4.018929254 | b2145 | *yeiS* | DUF2542 domain-containing protein YeiS |
| 0 | 32.3283 | 0 | | 16.1641699 | 4.014727518 | b2248 | *yfaX* | putative DNA-binding transcriptional regulator YfaX |
| 0 | 91.7759 | 0 | | 45.8879275 | 5.520042744 | b2263 | *menH* | 2-succinyl-6-hydroxy-2, 4-cyclohexadiene-1-carboxylate synthase |

| | | | | | | | | |
|---|---|---|---|---|---|---|---|---|
| 0 | 157.479 | 0 | | 78.7395807 | 6.299017125 | b2266 | *elaB* | tail anchored inner membrane protein |
| 0 | 45.2138 | 0 | | 22.60691 | 4.498691907 | b2543 | *yphA* | putative inner membrane protein |
| 0 | 62.7576 | 0 | | 31.3788053 | 4.971718519 | b2602 | *yfiL* | DUF2799 domain-containing lipoprotein YfiL |
| 0 | 161.936 | 0 | | 80.9677967 | 6.339276313 | b2694 | *argV* | tRNA-Arg |
| 0 | 287.095 | 0 | | 143.547303 | 7.16538241 | b2728 | *hypC* | hydrogenase 3 maturation protein HypC |
| 0 | 45.9731 | 0 | | 22.9865254 | 4.5227165 | b2763 | *cysI* | sulfite reductase, hemoprotein subunit |
| 0 | 39.1676 | 0 | | 19.5837899 | 4.291588079 | b2790 | *yqcA* | putative flavodoxin YqcA |
| 0 | 67.3421 | 0 | | 33.6710583 | 5.073437162 | b2801 | *fucP* | L-fucose:H(+) symporter |
| 0 | 65.2846 | 0 | | 32.6423004 | 5.028670827 | b2995 | *hybB* | hydrogenase 2 membrane subunit |
| 0 | 35.7553 | 0 | | 17.877669 | 4.160086738 | b3074 | *ygjH* | putative tRNA-binding protein YgjH |
| 0 | 1523 | 0 | | 761.499382 | 9.572699056 | b3118 | *tdcA* | DNA-binding transcriptional activator TdcA |
| 0 | 323.709 | 0 | | 161.8543 | 7.338551887 | b3158 | *yhbU* | putative peptidase YhbU |
| 0 | 96.8429 | 0 | | 48.4214585 | 5.59757463 | b3239 | *yhcO* | putative barnase inhibitor |
| 0 | 89.2878 | 0 | | 44.6438979 | 5.480391091 | b3362 | *yhfG* | DUF2559 domain-containing protein YhfG |
| 0 | 60.0343 | 0 | | 30.0171471 | 4.907714963 | b3408 | *feoA* | ferrous iron transport protein A |
| 0 | 58.7743 | 0 | | 29.387153 | 4.877113693 | b3509 | *hdeB* | periplasmic acid stress chaperone |
| 0 | 51.4927 | 0 | | 25.7463389 | 4.686295389 | b3512 | *gadE* | DNA-binding transcriptional activator GadE |
| 0 | 34.9454 | 0 | | 17.472698 | 4.127030488 | b3592 | *yibF* | glutathione transferase-like protein YibF |
| 0 | 67.847 | 0 | | 33.9234882 | 5.084212619 | b3761 | *trpT* | tRNA-Trp |
| 0 | 67.9364 | 0 | | 33.9681832 | 5.086112152 | b3811 | *xerC* | site-specific tyrosine recombinase |
| 0 | 268.756 | 0 | | 134.378133 | 7.070154578 | b3866 | *yihI* | Der GTPase-activating protein YihI |
| 0 | 251.723 | 0 | | 125.861451 | 6.975692666 | b4023 | *yjbD* | conserved protein YjbD |
| 0 | 36.9056 | 0 | | 18.4528111 | 4.205768711 | b4068 | *yjcH* | conserved inner membrane protein YjcH |
| 0 | 215.283 | 0 | | 107.641398 | 6.750089227 | b4128 | *ghoS* | antitoxin of the GhoTS toxin-antitoxin system |
| 0 | 74.1168 | 0 | | 37.0583861 | 5.211728146 | b4137 | *cutA* | copper binding protein CutA |
| 0 | 65.4683 | 0 | | 32.7341492 | 5.032724576 | b4397 | *creA* | PF05981 family protein CreA |
| 0 | 46.132 | 0 | | 23.0660139 | 4.527696805 | b4409 | *blr* | beta-lactam resistance protein |
| 0 | 41.6676 | 0 | | 20.833819 | 4.380855417 | b4410 | *ecnA* | entericidin A lipoprotein, antidote to entericidin B |
| 0 | 172.437 | 0 | | 86.2185279 | 6.429926024 | b4429 | *sokB* | putative small regulatory RNA SokB |
| 0 | 49.8587 | 0 | | 24.9293561 | 4.639773715 | b4468 | *glcE* | glycolate dehydrogenase, putative FAD-binding subunit |
| 0 | 310.863 | 0 | | 155.431255 | 7.280132828 | b4515 | *cydX* | cytochrome bd-I ubiquinol oxidase subunit CydX |
| 0 | 130.685 | 0 | | 65.3424324 | 6.029948256 | b4522 | *ymiA* | uncharacterized protein YmiA |
| 0 | 38.423 | 0 | | 19.2115134 | 4.263899271 | b4525 | *ymjC* | putative uncharacterized protein YmjC |
| 0 | 46.008 | 0 | | 23.0040085 | 4.523813371 | b4535 | *yniD* | uncharacterized protein YniD |
| 0 | 299.914 | 0 | | 149.956854 | 7.228403655 | b4551 | *yheV* | DUF2387 domain-containing protein YheV |
| 0 | 64.7337 | 0 | | 32.366826 | 5.016443991 | b4555 | *yicS* | uncharacterized protein YicS |
| 0 | 56.324 | 0 | | 28.1619937 | 4.815677568 | b4566 | *topAI* | KpLE2 phage-like element; toxin of the TopAI-YjhQ toxin-antitoxin system, TopA inhibitor |
| 0 | 331.062 | 0 | | 165.531203 | 7.370959381 | b4597 | *rydC* | small regulatory RNA RydC |
| 0 | 93.7522 | 0 | | 46.8760928 | 5.550780419 | b4613 | *dinQ* | UV inducible membrane toxin DinQ |
| 0 | 247.922 | 0 | | 123.961223 | 6.953745086 | b4624 | *ryjB* | small RNA RyjB |
| 0 | 227.684 | 0 | | 113.84194 | 6.830888338 | b4672 | *ymiB* | putative protein YmiB |
| 0 | 67.5016 | 0 | | 33.7507868 | 5.07684923 | b4675 | *yoaJ* | uncharacterized protein YoaJ |

| | | | | | | | | |
|---|---|---|---|---|---|---|---|---|
| 0 | 50.2244 | 0 | | 25.1121926 | 4.650316092 | b4679 | *yohP* | putative membrane protein YohP |
| 0 | 121.326 | 0 | | 60.6631789 | 5.922749196 | b4699 | *fnrS* | small regulatory RNA FnrS |
| 0 | 55.6157 | 0 | | 27.8078517 | 4.797420386 | b4700 | *sokE* | small RNA SokE |
| 0 | 59.784 | 0 | | 29.8920012 | 4.90168758 | b4704 | *arrS* | small regulatory RNA ArrS |
| 0 | 312.507 | 0 | | 156.253643 | 7.287746013 | b4727 | *yacM* | protein YacM |
| 0 | 669.197 | 0 | | 334.598318 | 8.386286381 | b4743 | *ynaL* | protein YnaL |
| 0 | 47.4548 | 0 | | 23.727405 | 4.56848242 | b1074 | *flgC* | flagellar basal-body rod protein FlgC |
| 0.270533 | 375.278 | 0.00072089 | -10.4379 | 187.774339 | 7.552856109 | b1593 | *ynfK* | putative dethiobiotin synthetase |
| 2.56178 | 3116.78 | 0.00082193 | -10.2487 | 1559.67181 | 10.60702677 | b4411 | *ecnB* | bacteriolytic entericidin B lipoprotein |
| 4.955021 | 2280.89 | 0.0021724 | -8.84649 | 1142.9235 | 10.15851313 | b0744 | *valT* | tRNA-Val |
| 0.164303 | 64.383 | 0.00255195 | -8.61418 | 32.273674 | 5.012285918 | b1440 | *ydcS* | putative ABC transporter periplasmic binding protein/polyhydroxybutyrate synthase |
| 2.197947 | 518.981 | 0.00423512 | -7.88338 | 260.589459 | 8.025634915 | b2876 | *yqeC* | uncharacterized protein YqeC |
| 9.509636 | 2108.48 | 0.00451019 | -7.7926 | 1058.99341 | 10.0484779 | b3117 | *tdcB* | catabolic threonine dehydratase |
| 2.164262 | 381.474 | 0.00567341 | -7.46157 | 191.819336 | 7.583604347 | b4370 | *leuQ* | tRNA-Leu |
| 21.01523 | 3342.28 | 0.00628769 | -7.31325 | 1681.6491 | 10.71566098 | b3127 | *garP* | galactarate/glucarate/glycerate transporter GarP |
| 5.162374 | 781.471 | 0.00660597 | -7.24201 | 393.316757 | 8.619547843 | b0621 | *dcuC* | anaerobic C4-dicarboxylate transporter DcuC |
| 5.318949 | 765.529 | 0.00694807 | -7.16917 | 385.424148 | 8.590303155 | b2146 | *preT* | NAD-dependent dihydropyrimidine dehydrogenase subunit PreT |
| 18.26922 | 2419.36 | 0.00755125 | -7.04907 | 1218.81656 | 10.2512653 | b1256 | *ompW* | outer membrane protein W |
| 1.316719 | 151.446 | 0.00869432 | -6.84571 | 76.3812786 | 6.255147165 | b4609 | *ryfD* | small regulatory RNA RyfD |
| 18.80466 | 2080.91 | 0.00903675 | -6.78998 | 1049.85703 | 10.03597716 | b3126 | *garL* | alpha-dehydro-beta-deoxy-D-glucarate aldolase |
| 27.54151 | 2942.59 | 0.00935961 | -6.73934 | 1485.06587 | 10.53631121 | b1587 | *ynfE* | putative selenate reductase YnfE |
| 5.341583 | 525.112 | 0.01017227 | -6.61921 | 265.226779 | 8.051082638 | b2702 | *srlA* | sorbitol-specific PTS enzyme IIC2 component |
| 9.222406 | 881.908 | 0.01045733 | -6.57934 | 445.565361 | 8.799493268 | b2762 | *cysH* | phosphoadenosine phosphosulfate reductase |
| 24.72505 | 2321.08 | 0.01065241 | -6.55268 | 1172.90095 | 10.19586546 | b3672 | *ivbL* | ilvBN operon leader peptide |
| 0.874754 | 81.9923 | 0.01066873 | -6.55047 | 41.4335418 | 5.372727243 | b1406 | *pdxI* | pyridoxine 4-dehydrogenase |
| 34.4712 | 3061.92 | 0.01125802 | -6.4729 | 1548.19746 | 10.59637377 | b2877 | *mocA* | molybdenum cofactor cytidylyltransferase |
| 12.21734 | 1044.57 | 0.01169601 | -6.41784 | 528.395417 | 9.045474144 | b3128 | *garD* | galactarate dehydratase |
| 4.70727 | 400.009 | 0.0117679 | -6.409 | 202.358298 | 7.660768197 | b2706 | *gutM* | DNA-binding transcriptional activator GutM |
| 8.48803 | 708.886 | 0.01197377 | -6.38398 | 358.686802 | 8.48658085 | b3476 | *nikA* | Ni(2(+)) ABC transporter periplasmic binding protein |
| 9.142729 | 757.03 | 0.0120771 | -6.37158 | 383.086586 | 8.581526701 | b4116 | *adiY* | DNA-binding transcriptional activator AdiY |
| 6.341966 | 442.083 | 0.01434563 | -6.12324 | 224.212722 | 7.808724328 | b1725 | *yniA* | putative kinase YniA |
| 75.71133 | 4777.21 | 0.01584844 | -5.97951 | 2426.4603 | 11.24463754 | b0904 | *focA* | formate channel FocA |
| 16.1879 | 1011.9 | 0.0159975 | -5.96601 | 514.044657 | 9.005749888 | b2287 | *nuoB* | NADH:quinone oxidoreductase subunit B |
| 269.734 | 15429.3 | 0.01748193 | -5.83799 | 7849.51859 | 12.93838846 | b0812 | *dps* | stationary phase nucleoid protein that sequesters iron and protects DNA from damage |
| 10.38846 | 578.462 | 0.01795876 | -5.79917 | 294.42511 | 8.201756904 | b1376 | *uspF* | nucleotide binding filament protein |
| 14.25067 | 776.513 | 0.01835214 | -5.76791 | 395.38167 | 8.627102179 | b1002 | *agp* | glucose-1-phosphatase |
| 1265.314 | 67710.3 | 0.01868717 | -5.74181 | 34487.8213 | 15.07379937 | b3707 | *tnaC* | tnaAB operon leader peptide |
| 6.350448 | 331.374 | 0.01916401 | -5.70546 | 168.862055 | 7.399701365 | b2869 | *ygeV* | putative sigma(54)-dependent transcriptional regulator YgeV |
| 57.10922 | 2913.92 | 0.01959877 | -5.67309 | 1485.51425 | 10.53674673 | b3510 | *hdeA* | HdeA monomer, chaperone active form |
| 12.18352 | 606.448 | 0.02008997 | -5.63738 | 309.315742 | 8.272936447 | b0707 | *ybgA* | DUF1722 domain-containing protein YbgA |
| 24.59308 | 1209.47 | 0.02033382 | -5.61997 | 617.030027 | 9.269196889 | b4153 | *frdB* | fumarate reductase iron-sulfur protein |
| 4.037425 | 195.241 | 0.02067919 | -5.59568 | 99.6391654 | 6.638641032 | b3588 | *aldB* | aldehyde dehydrogenase B |

| | | | | | | | | |
|---|---|---|---|---|---|---|---|---|
| 1.931188 | 92.3098 | 0.02092071 | -5.57892 | 47.120516 | 5.558283432 | b4435 | *isrC* | small RNA IsrC |
| 168.1641 | 7866.41 | 0.02137747 | -5.54776 | 4017.2895 | 11.97200672 | b4240 | *treB* | trehalose-specific PTS enzyme IIBC component |
| 114.2494 | 5273.56 | 0.02166456 | -5.52852 | 2693.9049 | 11.39548321 | b2579 | *grcA* | stress-induced alternate pyruvate formate-lyase subunit |
| 50.21088 | 2256.3 | 0.02225361 | -5.48982 | 1153.25674 | 10.17149801 | b1975 | *serU* | tRNA-Ser |
| 66.55106 | 2984.98 | 0.02229529 | -5.48712 | 1525.76722 | 10.57531916 | b1654 | *grxD* | glutaredoxin 4 |
| 0.794476 | 34.8109 | 0.02282259 | -5.45339 | 17.802707 | 4.154024725 | b1643 | *ydhI* | DUF1656 domain-containing protein YdhI |
| 182.0144 | 7713.82 | 0.0235959 | -5.40532 | 3947.91531 | 11.94687533 | b4189 | *bsmA* | DUF1471 domain-containing putative lipoprotein BsmA |
| 6.150833 | 260.319 | 0.0236281 | -5.40335 | 133.234701 | 7.057826068 | b2979 | *glcD* | glycolate dehydrogenase, putative FAD-linked subunit |
| 2.282313 | 93.7522 | 0.0243441 | -5.36028 | 48.0172492 | 5.585480851 | b2208 | *napF* | ferredoxin-type protein |
| 8.89151 | 353.784 | 0.02513257 | -5.3143 | 181.3379 | 7.502536669 | b0306 | *ykgE* | putative lactate utilization oxidoreductase YkgE |
| 91.10845 | 3596.1 | 0.02533534 | -5.30271 | 1843.60521 | 10.84831404 | b4035 | *malK* | maltose ABC transporter ATP binding subunit |
| 11.98052 | 430.833 | 0.02780782 | -5.16837 | 221.406605 | 7.790554449 | b0754 | *aroG* | 3-deoxy-7-phosphoheptulonate synthase, Phe-sensitive |
| 6.53601 | 230.391 | 0.02836926 | -5.13953 | 118.463272 | 6.888296033 | b1673 | *ydhV* | putative oxidoreductase |
| 93.0525 | 3254.48 | 0.02859212 | -5.12824 | 1673.76671 | 10.70888275 | b4154 | *frdA* | fumarate reductase flavoprotein subunit |
| 7.214207 | 249.647 | 0.02889767 | -5.1129 | 128.430416 | 7.0048431 | b2219 | *atoS* | sensory histidine kinase AtoS |
| 76.27729 | 2530.41 | 0.03014422 | -5.05197 | 1303.34458 | 10.34800284 | b0897 | *ycaC* | putative hydrolase |
| 3.79109 | 121.997 | 0.03107535 | -5.00809 | 62.8938909 | 5.974847985 | b2788 | *gudX* | glucarate dehydratase-related protein |
| 751.9664 | 22895.2 | 0.03284385 | -4.92823 | 11823.5819 | 13.52937954 | b3708 | *tnaA* | tryptophanase |
| 4.553826 | 137.129 | 0.03320843 | -4.91231 | 70.8412085 | 6.146516918 | b2464 | *talA* | transaldolase A |
| 1086.743 | 31986 | 0.03397556 | -4.87936 | 16536.3845 | 14.01335622 | b4034 | *malE* | maltose ABC transporter periplasmic binding protein |
| 45.785 | 1345.87 | 0.03401886 | -4.87752 | 695.828019 | 9.442586963 | b3087 | *ygjR* | putative oxidoreductase YgjR |
| 21.66177 | 631.928 | 0.03427884 | -4.86654 | 326.795111 | 8.35224259 | b1739 | *osmE* | osmotically-inducible lipoprotein OsmE |
| 17.2334 | 498.529 | 0.03456852 | -4.8544 | 257.880983 | 8.010561578 | b3125 | *garR* | tartronate semialdehyde reductase |
| 67.08428 | 1889.25 | 0.03550835 | -4.8157 | 978.169008 | 9.933939946 | b4239 | *treC* | trehalose-6-phosphate hydrolase |
| 33.05869 | 911.519 | 0.03626768 | -4.78517 | 472.289016 | 8.883526172 | b3024 | *ygiW* | BOF family protein YgiW |
| 21.12591 | 578.291 | 0.0365316 | -4.77471 | 299.708638 | 8.227416855 | b1188 | *ycgB* | PF04293 family protein YcgB |
| 7.82464 | 205.575 | 0.0380622 | -4.7155 | 106.699862 | 6.737414499 | b1732 | *katE* | catalase II |
| 34.47578 | 901.429 | 0.03824568 | -4.70856 | 467.952623 | 8.870218663 | b0380 | *yaiZ* | DUF2754 domain-containing protein YaiZ |
| 12.55272 | 322.23 | 0.03895581 | -4.68202 | 167.391227 | 7.387080108 | b2977 | *glcG* | putative heme-binding protein GlcG |
| 1.321339 | 33.8824 | 0.03899783 | -4.68046 | 17.6018539 | 4.137655481 | b1781 | *yeaE* | methylglyoxal reductase YeaE |
| 9.733392 | 248 | 0.03924748 | -4.67126 | 128.866913 | 7.009738086 | b1498 | *ydeN* | putative sulfatase |
| 14.48391 | 366.556 | 0.03951346 | -4.66151 | 190.52011 | 7.573799475 | b3115 | *tdcD* | propionate kinase |
| 26.07657 | 615.254 | 0.0423834 | -4.56036 | 320.665441 | 8.324925068 | b2957 | *ansB* | L-asparaginase 2 |
| 2.356736 | 53.855 | 0.04376076 | -4.51422 | 28.1058732 | 4.812799732 | b4342 | *yjiT* | putative uncharacterized protein YjiT |
| 26.29286 | 599.747 | 0.04383995 | -4.51161 | 313.019693 | 8.290109613 | b3116 | *tdcC* | threonine/serine:H(+) symporter |
| 95.50982 | 2175.32 | 0.04390605 | -4.50944 | 1135.41614 | 10.14900544 | b1480 | *sra* | 30S ribosomal subunit protein S22 |
| 63.56826 | 1441.14 | 0.04410972 | -4.50276 | 752.353813 | 9.555267475 | b1461 | *pptA* | putative 4-oxalocrotonate tautomerase (4-OT) |
| 54.56333 | 1230.87 | 0.04432906 | -4.4956 | 642.716867 | 9.328039523 | b1004 | *wrbA* | NAD(P)H:quinone oxidoreductase |
| 226.8481 | 5092.06 | 0.04454936 | -4.48845 | 2659.45445 | 11.37691461 | b0929 | *ompF* | outer membrane porin F |
| 70.84208 | 1561.51 | 0.04536782 | -4.46219 | 816.173566 | 9.672732175 | b4118 | *melR* | DNA-binding transcriptional dual regulator MelR |
| 43.94666 | 945.682 | 0.04647086 | -4.42753 | 494.814382 | 8.950743623 | b4380 | *yjjI* | DUF3029 domain-containing protein YjjI |
| 1.859662 | 39.8157 | 0.04670671 | -4.42023 | 20.8377026 | 4.381124322 | b0736 | *ybgC* | esterase/thioesterase |

| | | | | | | | | |
|---|---|---|---|---|---|---|---|---|
| 45.59749 | 968.238 | 0.04709324 | -4.40834 | 506.917935 | 8.985608399 | b2726 | *hypA* | hydrogenase 3 nickel incorporation protein HypA |
| 17.19752 | 364.689 | 0.04715662 | -4.4064 | 190.943468 | 7.577001755 | b2535 | *csiE* | stationary phase inducible protein CsiE |
| 67.66701 | 1390.66 | 0.04865829 | -4.36117 | 729.162213 | 9.510095989 | b2504 | *yfgG* | protein YfgG |
| 16.32874 | 335.12 | 0.04872512 | -4.35919 | 175.724171 | 7.45716884 | b0579 | *ybdF* | PF04237 family protein YbdF |
| 44.03741 | 894.795 | 0.04921506 | -4.34476 | 469.41643 | 8.874724529 | b0030 | *rihC* | ribonucleoside hydrolase RihC |
| 18.18592 | 367.997 | 0.04941871 | -4.3388 | 193.091286 | 7.593139249 | b1197 | *treA* | periplasmic trehalase |
| 9.795475 | 194.369 | 0.05039636 | -4.31054 | 102.082084 | 6.673585878 | b3153 | *yhbO* | protein/nucleic acid deglycase 2 |
| 42.40329 | 839.916 | 0.05048517 | -4.308 | 441.159491 | 8.785156514 | b2997 | *hybO* | hydrogenase 2 small subunit |
| 4.738341 | 91.6636 | 0.0516927 | -4.2739 | 48.2009939 | 5.590990991 | b1588 | *ynfF* | putative selenate reductase YnfF |
| 23.87327 | 461.015 | 0.05178411 | -4.27135 | 242.44431 | 7.921509583 | b0894 | *dmsA* | dimethyl sulfoxide reductase subunit A |
| 16.16638 | 312.034 | 0.05180971 | -4.27063 | 164.100086 | 7.35843218 | b4152 | *frdC* | fumarate reductase membrane protein FrdC |
| 20.25799 | 384.555 | 0.05267904 | -4.24663 | 202.406533 | 7.661112047 | b3001 | *gpr* | L-glyceraldehyde 3-phosphate reductase |
| 8.008897 | 151.452 | 0.05288073 | -4.24111 | 79.7304976 | 6.317059768 | b4335 | *yjiM* | putative dehydratase subunit |
| 4.161123 | 76.4866 | 0.0544033 | -4.20016 | 40.3238562 | 5.333561705 | b0946 | *zapC* | cell division protein ZapC |
| 57.84838 | 1063.28 | 0.0544057 | -4.2001 | 560.563089 | 9.130732941 | b1905 | *ftnA* | ferritin iron storage protein |
| 12.63423 | 232.148 | 0.05442311 | -4.19964 | 122.39125 | 6.935356611 | b1799 | *dmlR* | DNA-binding transcriptional regulator DmlR |
| 215.8084 | 3946.57 | 0.05468252 | -4.19278 | 2081.18948 | 11.02319261 | b3544 | *dppA* | dipeptide ABC transporter periplasmic binding protein |
| 260.4221 | 4736.58 | 0.05498099 | -4.18492 | 2498.5032 | 11.28684835 | b3049 | *glgS* | surface composition regulator |
| 7.049535 | 124.776 | 0.05649731 | -4.14567 | 65.9129972 | 6.042491068 | b3012 | *dkgA* | methylglyoxal reductase DkgA |
| 32.46393 | 566.285 | 0.05732793 | -4.12462 | 299.374342 | 8.225806769 | b0722 | *sdhD* | succinate:quinone oxidoreductase, membrane protein SdhD |
| 357.7952 | 6094.53 | 0.0587076 | -4.09031 | 3226.16252 | 11.6556034 | b0280 | *yagN* | CP4-6 prophage; protein YagN |
| 12.27983 | 209.074 | 0.05873435 | -4.08965 | 110.677001 | 6.790211648 | b0110 | *ampD* | 1,6-anhydro-N-acetylmuramoyl-L-alanine amidase |
| 1716.639 | 29203.4 | 0.05878213 | -4.08848 | 15460.0308 | 13.91625557 | b2096 | *gatY* | tagatose-1,6-bisphosphate aldolase 2 subunit GatY |
| 30.39598 | 514.901 | 0.05903271 | -4.08234 | 272.648366 | 8.090897697 | b3124 | *garK* | glycerate 2-kinase 1 |
| 63.25586 | 1039.73 | 0.06083873 | -4.03887 | 551.492992 | 9.107198743 | b2277 | *nuoM* | NADH:quinone oxidoreductase subunit M |
| 2.085799 | 33.9887 | 0.06136742 | -4.02638 | 18.0372536 | 4.172907779 | b1515 | *lsrD* | Autoinducer-2 ABC transporter membrane subunit LsrD |
| 10.18386 | 164.686 | 0.06183793 | -4.01536 | 87.4351217 | 6.450141005 | b0445 | *ybaE* | putative protein YbaE |
| 1.895545 | 30.5167 | 0.0621151 | -4.00891 | 16.206098 | 4.018464863 | b1464 | *yddE* | PF02567 family protein YddE |
| 3.351454 | 53.3993 | 0.06276214 | -3.99396 | 28.3753787 | 4.826567743 | b0895 | *dmsB* | dimethyl sulfoxide reductase subunit B |
| 7.563478 | 120.378 | 0.06283103 | -3.99238 | 63.9707744 | 5.999341042 | b0708 | *phr* | deoxyribodipyrimidine photolyase (photoreactivation) |
| 8.289532 | 130.31 | 0.06361411 | -3.97451 | 69.2995868 | 6.114774845 | b0853 | *ybjN* | protein YbjN |
| 12.55272 | 196.956 | 0.06373369 | -3.9718 | 104.754265 | 6.710865179 | b2369 | *evgA* | DNA-binding transcriptional activator EvgA |
| 11.4963 | 178.377 | 0.06444958 | -3.95569 | 94.936493 | 6.568890852 | b1412 | *azoR* | FMN dependent NADH:quinone oxidoreductase |
| 22.18151 | 343.381 | 0.06459729 | -3.95238 | 182.781506 | 7.513976295 | b1539 | *ydfG* | 3-hydroxy acid dehydrogenase |
| 7.230121 | 109.301 | 0.06614874 | -3.91814 | 58.2655377 | 5.86457092 | b4467 | *glcF* | glycolate dehydrogenase, putative iron-sulfur subunit |
| 25.90578 | 385.391 | 0.06721953 | -3.89498 | 205.648238 | 7.684034903 | b0919 | *ycbJ* | putative phosphotransferase YcbJ |
| 56.00444 | 825.981 | 0.06780356 | -3.8825 | 440.992618 | 8.784610697 | b2904 | *gcvH* | glycine cleavage system H protein |
| 110.5185 | 1585.63 | 0.06969985 | -3.8427 | 848.076651 | 9.728050855 | b0329 | *yahO* | DUF1471 domain-containing protein YahO |
| 9.889223 | 141.81 | 0.06973571 | -3.84196 | 75.8496256 | 6.245070154 | b0307 | *ykgF* | amino acid dehydrogenase with NAD(P)-binding domain and ferridoxin-like domain |
| 86.29995 | 1234.18 | 0.06992502 | -3.83805 | 660.23918 | 9.366844943 | b3709 | *tnaB* | tryptophan:H(+) symporter TnaB |
| 127.8095 | 1821.82 | 0.07015474 | -3.83332 | 974.816143 | 9.928986332 | b0525 | *ppiB* | peptidyl-prolyl cis-trans isomerase B |
| 71.01138 | 992.01 | 0.07158331 | -3.80423 | 531.510843 | 9.053955313 | b0651 | *rihA* | pyrimidine-specific ribonucleoside hydrolase RihA |

| | | | | | | | | |
|---|---|---|---|---|---|---|---|---|
| 41.41103 | 577.226 | 0.0717415 | -3.80105 | 309.318345 | 8.27294859 | b2727 | *hypB* | hydrogenase isoenzymes nickel incorporation protein HypB |
| 7.90855 | 108.079 | 0.07317384 | -3.77253 | 57.9937386 | 5.85782524 | b0581 | *ybdK* | carboxylate-amine ligase |
| 26.18481 | 356.497 | 0.07345027 | -3.76709 | 191.340989 | 7.58000215 | b3805 | *hemC* | hydroxymethylbilane synthase |
| 41.79549 | 560.411 | 0.07458006 | -3.74507 | 301.103269 | 8.23411456 | b2284 | *nuoF* | NADH:quinone oxidoreductase subunit F |
| 33.51773 | 448.963 | 0.07465584 | -3.7436 | 241.24054 | 7.914328558 | b0453 | *ybaY* | PF09619 family lipoprotein YbaY |
| 29.21107 | 388.726 | 0.07514566 | -3.73417 | 208.96852 | 7.707141813 | b4350 | *hsdR* | type I restriction enzyme EcoKI endonuclease component |
| 214.8197 | 2841.08 | 0.07561206 | -3.72524 | 1527.94879 | 10.57738047 | b1656 | *sodB* | superoxide dismutase (Fe) |
| 7.075169 | 91.7067 | 0.07714999 | -3.69619 | 49.3909264 | 5.626174123 | b3516 | *gadX* | DNA-binding transcriptional dual regulator GadX |
| 5.192123 | 67.1396 | 0.07733319 | -3.69277 | 36.1658828 | 5.176557463 | b1341 | *dgcM* | diguanylate cyclase DgcM |
| 339.5959 | 4285.54 | 0.07924236 | -3.65758 | 2312.56554 | 11.17527854 | b4108 | *yjdM* | conserved protein YjdM |
| 18.36983 | 228.664 | 0.08033553 | -3.63782 | 123.516851 | 6.948564063 | b1061 | *dinI* | DNA damage-inducible protein I |
| 20.82738 | 259.01 | 0.0804116 | -3.63645 | 139.918532 | 7.12844325 | b1276 | *acnA* | aconitate hydratase 1 |
| 97.20704 | 1179.97 | 0.0823806 | -3.60155 | 638.590942 | 9.31874828 | b3418 | *malT* | DNA-binding transcriptional activator MalT |
| 10.05304 | 121.756 | 0.08256708 | -3.59829 | 65.9045646 | 6.042306486 | b0906 | *ycaP* | DUF421 domain-containing protein YcaP |
| 6.258132 | 74.8142 | 0.08364901 | -3.57951 | 40.536158 | 5.341137451 | b2878 | *ygfK* | putative oxidoreductase, Fe-S subunit |
| 20.05604 | 236.73 | 0.0847211 | -3.56113 | 128.39309 | 7.004423751 | b1976 | *mtfA* | Mlc titration factor |
| 7.05209 | 82.9841 | 0.08498117 | -3.55671 | 45.0181171 | 5.49243381 | b0678 | *nagB* | glucosamine-6-phosphate deaminase |
| 5.777402 | 67.383 | 0.08573974 | -3.54389 | 36.5802119 | 5.192991527 | b0799 | *dinG* | ATP-dependent DNA helicase DinG |
| 143.4597 | 1673.03 | 0.0857484 | -3.54375 | 908.244865 | 9.826937493 | b4722 | *idlP* | iraD leader peptide |
| 10.26507 | 119.162 | 0.08614411 | -3.5371 | 64.7133656 | 6.015991805 | b1662 | *ribC* | riboflavin synthase |
| 39.31999 | 439.8 | 0.08940415 | -3.48351 | 239.560214 | 7.904244514 | b2660 | *lhgO* | L-2-hydroxyglutarate oxidase |
| 6.537875 | 72.7882 | 0.08982059 | -3.47681 | 39.6630151 | 5.309722446 | b4124 | *dcuR* | DNA-binding transcriptional activator DcuR |
| 43.66163 | 485.745 | 0.08988591 | -3.47576 | 264.703327 | 8.048232517 | b4188 | *yjfN* | protease activator |
| 99.51765 | 1097.4 | 0.09068523 | -3.46299 | 598.457028 | 9.225103849 | b2903 | *gcvP* | glycine decarboxylase |
| 8.724985 | 95.9944 | 0.09089057 | -3.45973 | 52.3596875 | 5.710384582 | b3638 | *yicR* | RadC-like JAB domain-containing protein YicR |
| 9.260203 | 101.847 | 0.09092301 | -3.45921 | 55.5534199 | 5.795803822 | b4357 | *lgoR* | putative DNA-binding transcriptional regulator LgoR |
| 50.67868 | 554.846 | 0.09133828 | -3.45264 | 302.762346 | 8.242041981 | b3750 | *rbsC* | ribose ABC transporter membrane subunit |
| 59.2834 | 636.069 | 0.0932028 | -3.42348 | 347.676122 | 8.441600176 | b0733 | *cydA* | cytochrome bd-I ubiquinol oxidase subunit I |
| 107.2211 | 1136.31 | 0.09435896 | -3.4057 | 621.76618 | 9.280228336 | b4449 | *sraG* | small regulatory RNA SraG |
| 5.811444 | 61.4598 | 0.09455689 | -3.40267 | 33.6356052 | 5.071917313 | b2327 | *yfcA* | conserved inner membrane protein YfcA |
| 20.37216 | 214.05 | 0.09517469 | -3.39328 | 117.211194 | 6.872966545 | b2976 | *glcB* | malate synthase G |
| 4.531668 | 47.6094 | 0.09518428 | -3.39313 | 26.0705405 | 4.704348587 | b0867 | *amiD* | N-acetylmuramoyl-L-alanine amidase D |
| 39.30837 | 409.814 | 0.09591762 | -3.38206 | 224.561103 | 7.810964248 | b3571 | *malS* | alpha-amylase |
| 28.88701 | 297.209 | 0.09719433 | -3.36298 | 163.047912 | 7.349152155 | b3812 | *yigB* | 5-amino-6-(5-phospho-D-ribitylamino)uracil phosphatase |
| 43.29894 | 445.422 | 0.09720885 | -3.36277 | 244.360358 | 7.932866446 | b2009 | *sbmC* | DNA gyrase inhibitor |
| 8.302063 | 85.3194 | 0.09730564 | -3.36133 | 46.8107564 | 5.548768171 | b3506 | *slp* | starvation lipoprotein |
| 7.627985 | 78.0935 | 0.09767759 | -3.35583 | 42.8607452 | 5.421585029 | b0681 | *chiP* | chitobiose outer membrane channel |
| 5.079135 | 51.9641 | 0.09774312 | -3.35486 | 28.521628 | 4.833984429 | b2176 | *pdeN* | putative c-di-GMP phosphodiesterase PdeN |
| 102.6507 | 1043.25 | 0.09839529 | -3.34527 | 572.949683 | 9.162264636 | b0881 | *clpS* | specificity factor for ClpA-ClpP chaperone-protease complex |
| 38.43504 | 387.733 | 0.0991276 | -3.33457 | 213.084009 | 7.73527852 | b1244 | *oppB* | murein tripeptide ABC transporter/oligopeptide ABC transporter inner membrane subunit |
| 60.9374 | 612.237 | 0.09953242 | -3.32869 | 336.587054 | 8.394835876 | b4069 | *acs* | acetyl-CoA synthetase (AMP-forming) |
| 6.084226 | 60.5881 | 0.10041942 | -3.31589 | 33.3361868 | 5.059017185 | b2578 | *eamB* | cysteine/O-acetylserine exporter EamB |

| | | | | | | | | |
|---|---|---|---|---|---|---|---|---|
| 6.109731 | 60.6578 | 0.10072464 | -3.31151 | 33.3837436 | 5.06107384 | b2425 | *cysP* | thiosulfate/sulfate ABC transporter periplasmic binding protein CysP |
| 1349.417 | 13207.8 | 0.10216807 | -3.29098 | 7278.61813 | 12.82944886 | b2597 | *raiA* | stationary phase translation inhibitor and ribosome stability factor |
| 15.6909 | 153.35 | 0.10232096 | -3.28883 | 84.5203569 | 6.401226953 | b4119 | *melA* | alpha-galactosidase |
| 14.24251 | 138.705 | 0.1026819 | -3.28375 | 76.4738328 | 6.256894278 | b2705 | *srlD* | sorbitol-6-phosphate 2-dehydrogenase |
| 10.88756 | 105.87 | 0.10283916 | -3.28154 | 58.3786891 | 5.867369909 | b1071 | *flgM* | anti-sigma factor for FliA (sigma(28)) |
| 18.06311 | 173.786 | 0.10393887 | -3.26619 | 95.9245199 | 6.583827735 | b4122 | *fumB* | fumarase B |
| 118.1432 | 1132.38 | 0.10433186 | -3.26075 | 625.261293 | 9.288315399 | b0963 | *mgsA* | methylglyoxal synthase |
| 30.76038 | 291.467 | 0.10553633 | -3.24419 | 161.113784 | 7.331936121 | b4376 | *osmY* | periplasmic chaperone OsmY |
| 5.316043 | 48.9253 | 0.10865626 | -3.20216 | 27.1206817 | 4.761321539 | b2422 | *cysA* | sulfate/thiosulfate ABC transporter ATP binding subunit |
| 19.63091 | 180.07 | 0.10901802 | -3.19736 | 99.8505914 | 6.641699067 | b0620 | *dpiA* | DNA-binding transcriptional dual regulator DpiA |
| 10.88386 | 99.1714 | 0.109748 | -3.18773 | 55.0276241 | 5.782084135 | b1448 | *mnaT* | L-amino acid N-acyltransferase |
| 46.09202 | 416.514 | 0.11066149 | -3.17577 | 231.302817 | 7.853639028 | b0389 | *yaiA* | protein YaiA |
| 12.84272 | 115.617 | 0.11107946 | -3.17034 | 64.2300492 | 6.005176497 | b2752 | *cysD* | sulfate adenylyltransferase subunit 2 |
| 39.75028 | 356.258 | 0.11157713 | -3.16389 | 198.004292 | 7.629387896 | b4151 | *frdD* | fumarate reductase membrane protein FrdD |
| 133.1527 | 1184.9 | 0.1123747 | -3.15361 | 659.0259 | 9.364191355 | b1085 | *yceQ* | DUF2655 domain-containing protein YceQ |
| 9.098865 | 80.6232 | 0.11285669 | -3.14744 | 44.8610231 | 5.487390619 | b1235 | *rssB* | regulator of RpoS |
| 14.29079 | 125.868 | 0.11353762 | -3.13876 | 70.0795538 | 6.130921685 | b0419 | *yajO* | 1-deoxyxylulose-5-phosphate synthase YajO |
| 79.32338 | 693.318 | 0.1144112 | -3.1277 | 386.320828 | 8.593655651 | b1000 | *cbpA* | curved DNA-binding protein |
| 52.96506 | 454.125 | 0.11663115 | -3.09997 | 253.544787 | 7.986096801 | b1750 | *ydjX* | DedA family protein YdjX |
| 37.29804 | 317.887 | 0.11733131 | -3.09134 | 177.592275 | 7.472425016 | b2946 | *rsmE* | 16S rRNA m(3)U1498 methyltransferase |
| 18.68159 | 159.191 | 0.11735329 | -3.09107 | 88.9363208 | 6.474700817 | b0677 | *nagA* | N-acetylglucosamine-6-phosphate deacetylase |
| 76.56728 | 644.774 | 0.11875048 | -3.07399 | 360.670879 | 8.494539135 | b3886 | *yihY* | putative inner membrane protein |
| 4.916482 | 40.8343 | 0.12040083 | -3.05408 | 22.8753836 | 4.515724032 | b2764 | *cysJ* | sulfite reductase, flavoprotein subunit |
| 70.96517 | 583.748 | 0.12156813 | -3.04016 | 327.356652 | 8.354719485 | b3942 | *katG* | hydroperoxidase I |
| 491.9992 | 4035.89 | 0.12190606 | -3.03616 | 2263.94375 | 11.1446224 | b4036 | *lamB* | maltose outer membrane channel/phage lambda receptor protein |
| 8.604687 | 70.3141 | 0.12237492 | -3.03062 | 39.4594131 | 5.302297593 | b2704 | *srlB* | sorbitol-specific PTS enzyme IIA component |
| 43.66774 | 353.266 | 0.12361147 | -3.01612 | 198.466902 | 7.632754623 | b4123 | *dcuB* | anaerobic C4-dicarboxylate transporter DcuB |
| 3.950157 | 31.9063 | 0.12380476 | -3.01386 | 17.9282474 | 4.164162558 | b2382 | *ypdC* | putative AraC-type DNA-binding transcriptional regulator YpdC |
| 146.5615 | 1183.3 | 0.12385778 | -3.01324 | 664.933049 | 9.377065276 | b2280 | *nuoJ* | NADH:quinone oxidoreductase subunit J |
| 94.1454 | 758.577 | 0.12410782 | -3.01033 | 426.361433 | 8.735933133 | b4021 | *pepE* | peptidase E |
| 25.05378 | 200.879 | 0.12472066 | -3.00323 | 112.966476 | 6.819750885 | b2565 | *recO* | DNA repair protein RecO |
| 8.448946 | 67.5096 | 0.1251518 | -2.99825 | 37.9792663 | 5.247140132 | b0829 | *gsiA* | glutathione ABC transporter ATP binding subunit GsiA |
| 6.747087 | 53.7513 | 0.12552427 | -2.99396 | 30.24917 | 4.918823652 | b1116 | *lolC* | lipoprotein release complex - inner membrane subunit |
| 11.34788 | 90.3341 | 0.1256212 | -2.99285 | 50.84101 | 5.667920787 | b1423 | *ydcJ* | DUF1338 domain-containing protein YdcJ |
| 529.3256 | 4212.28 | 0.12566255 | -2.99237 | 2370.80175 | 11.21115931 | b0903 | *pflB* | pyruvate formate-lyase |
| 13.04986 | 103.138 | 0.12652847 | -2.98247 | 58.0937876 | 5.860311988 | b0268 | *yagE* | CP4-6 prophage; putative 2-keto-3-deoxygluconate aldolase |
| 134.6802 | 1063.83 | 0.12659976 | -2.98165 | 599.253554 | 9.227022749 | b1824 | *yobF* | DUF2527 domain-containing protein YobF |
| 47.36435 | 373.382 | 0.12685216 | -2.97878 | 210.373331 | 7.716808017 | b3480 | *nikE* | Ni(2(+)) ABC transporter ATP binding subunit NikE |
| 29.92218 | 235.107 | 0.12727036 | -2.97403 | 132.514703 | 7.050008632 | b0865 | *ybjP* | DUF3828 domain-containing lipoprotein YbjP |
| 76.30085 | 596.521 | 0.12790968 | -2.9668 | 336.411053 | 8.394081296 | b3098 | *yqjD* | ribosome- and membrane-associated DUF883 domain-containing protein YqjD |
| 171.1324 | 1330.97 | 0.12857757 | -2.95929 | 751.049342 | 9.552763882 | b4056 | *yjbQ* | UPF0047 protein YjbQ |
| 10.57071 | 82.0414 | 0.12884609 | -2.95628 | 46.3060489 | 5.533128757 | b1896 | *otsA* | trehalose-6-phosphate synthase |

| | | | | | | | | |
|---|---|---|---|---|---|---|---|---|
| 6.17752 | 47.2452 | 0.13075445 | -2.93507 | 26.7113578 | 4.739381406 | b2842 | *kduD* | putative 2-keto-3-deoxy-D-gluconate dehydrogenase |
| 286.0113 | 2166.98 | 0.13198609 | -2.92154 | 1226.49612 | 10.26032695 | b1677 | *lpp* | murein lipoprotein |
| 39.00728 | 295.276 | 0.13210467 | -2.92025 | 167.141429 | 7.384925569 | b1132 | *hflD* | lysogenization regulator |
| 4.55825 | 34.4806 | 0.13219771 | -2.91923 | 19.5194013 | 4.2868369 | b4377 | *yjjU* | putative patatin-like phospholipase YjjU |
| 11.01116 | 82.056 | 0.13419076 | -2.89764 | 46.5335821 | 5.540200344 | b0782 | *moaB* | MoaB protein |
| 45.22671 | 335.486 | 0.13480963 | -2.891 | 190.356237 | 7.572558028 | b0627 | *tatE* | twin arginine protein translocation system - TatE protein |
| 135.6746 | 1004.51 | 0.13506487 | -2.88828 | 570.094521 | 9.155057327 | b2285 | *nuoE* | NADH:quinone oxidoreductase subunit E |
| 164.5986 | 1217.12 | 0.13523581 | -2.88645 | 690.860662 | 9.432250955 | b2094 | *gatA* | galactitol-specific PTS enzyme IIA component |
| 31.89626 | 234.637 | 0.13593895 | -2.87897 | 133.266436 | 7.058169669 | b1912 | *pgsA* | phosphatidylglycerophosphate synthase |
| 101.8084 | 747.474 | 0.13620329 | -2.87617 | 424.641109 | 8.730100232 | b3838 | *tatB* | twin arginine protein translocation system - TatB protein |
| 15.03602 | 110.212 | 0.1364278 | -2.87379 | 62.6241574 | 5.968647382 | b2370 | *evgS* | sensory histidine kinase EvgS |
| 38.91077 | 283.778 | 0.137117 | -2.86652 | 161.344307 | 7.333998861 | b3092 | *uxaC* | D-glucoronate/D-galacturonate isomerase |
| 25.25852 | 183.312 | 0.13778964 | -2.85946 | 104.285361 | 6.704392844 | b3470 | *tusA* | sulfur transfer protein TusA |
| 20.13172 | 145.64 | 0.13822916 | -2.85487 | 82.8859541 | 6.373055738 | b0773 | *ybhB* | putative kinase inhibitor |
| 12.67059 | 91.4047 | 0.13862071 | -2.85079 | 52.0376494 | 5.70148389 | b4219 | *msrA* | methionine sulfoxide reductase A |
| 92.76559 | 668.491 | 0.13876869 | -2.84925 | 380.628186 | 8.572238588 | b2584 | *patZ* | peptidyl-lysine acetyltransferase |
| 34.447 | 248.116 | 0.13883411 | -2.84857 | 141.281624 | 7.142430021 | b1783 | *yeaG* | protein kinase YeaG |
| 110.005 | 790.677 | 0.13912763 | -2.84552 | 450.341026 | 8.814874104 | b1205 | *ychH* | stress-induced protein |
| 12.17301 | 87.3019 | 0.13943588 | -2.84233 | 49.7374343 | 5.636260182 | b2126 | *btsS* | high-affinity pyruvate receptor |
| 948.3118 | 6771.91 | 0.14003615 | -2.83613 | 3860.10923 | 11.91442596 | b2095 | *gatZ* | tagatose-1,6-bisphosphate aldolase 2 subunit GatZ |
| 23.23773 | 165.553 | 0.14036418 | -2.83275 | 94.3954507 | 6.560645427 | b2989 | *yghU* | disulfide reductase/organic hydroperoxide reductase |
| 14.254 | 100.506 | 0.14182241 | -2.81784 | 57.3800051 | 5.842476192 | b3661 | *nlpA* | lipoprotein-28 |
| 547.0522 | 3840.51 | 0.14244252 | -2.81155 | 2193.78227 | 11.09920463 | b4139 | *aspA* | aspartate ammonia-lyase |
| 6.627212 | 45.6144 | 0.14528767 | -2.78302 | 26.1208149 | 4.707128 | b0505 | *allA* | ureidoglycolate lyase |
| 1455.247 | 9980.12 | 0.14581463 | -2.77779 | 5717.68356 | 12.48121506 | b0605 | *ahpC* | alkyl hydroperoxide reductase, AhpC component |
| 106.9306 | 731.064 | 0.14626695 | -2.77332 | 418.997537 | 8.710797952 | b3430 | *glgC* | glucose-1-phosphate adenylyltransferase |
| 21.42619 | 145.585 | 0.1471728 | -2.76442 | 83.5057424 | 6.383803504 | b3821 | *pldA* | outer membrane phospholipase A |
| 2473.398 | 16792.5 | 0.14729198 | -2.76325 | 9632.94074 | 13.23376058 | b0557 | *borD* | DLP12 prophage; prophage lipoprotein BorD |
| 6.671721 | 44.5384 | 0.14979692 | -2.73892 | 25.6050805 | 4.678358191 | b0790 | *ybhP* | endonuclease/exonuclease/phosphatase domain-containing protein YbhP |
| 123.9482 | 821.757 | 0.15083328 | -2.72897 | 472.852409 | 8.885246138 | b3336 | *bfr* | bacterioferritin |
| 64.89842 | 427.731 | 0.15172715 | -2.72045 | 246.314738 | 7.944359141 | b3448 | *yhhA* | DUF2756 domain-containing protein YhhA |
| 746.8274 | 4920.17 | 0.15178907 | -2.71986 | 2833.4967 | 11.46836781 | b1333 | *uspE* | universal stress protein with a role cellular motility |
| 7.600868 | 49.9112 | 0.15228771 | -2.71513 | 28.7560515 | 4.845793688 | b3947 | *ptsA* | putative PTS multiphosphoryl transfer protein PtsA |
| 9.34777 | 61.3616 | 0.15233905 | -2.71464 | 35.3546915 | 5.143829765 | b2996 | *hybA* | hydrogenase 2 iron-sulfur protein |
| 23.01843 | 150.523 | 0.15292297 | -2.70912 | 86.770752 | 6.439136927 | b1127 | *pepT* | peptidase T |
| 93.21694 | 609.297 | 0.15299104 | -2.70848 | 351.256846 | 8.456382533 | b1246 | *oppD* | murein tripeptide ABC transporter/oligopeptide ABC transporter ATP binding subunit |
| 85.79723 | 558.933 | 0.15350183 | -2.70367 | 322.365082 | 8.332551672 | b4033 | *malF* | maltose ABC transporter membrane subunit MalF |
| 371.1239 | 2408.48 | 0.1540905 | -2.69815 | 1389.80198 | 10.44066362 | b2398 | *yfeC* | putative DNA-binding transcriptional regulator YfeC |
| 13.58602 | 88.0356 | 0.15432421 | -2.69596 | 50.810806 | 5.667063443 | b1342 | *zntB* | Zn(2(+)):H(+) symporter |
| 8.790993 | 56.801 | 0.15476828 | -2.69182 | 32.7959956 | 5.035447766 | b2351 | *yfdH* | CPS-53 (KpLE1) prophage; bactoprenol glucosyl transferase |
| 89.37537 | 577.263 | 0.15482595 | -2.69128 | 333.319411 | 8.380761527 | b1051 | *msyB* | acidic protein that suppresses heat sensitivity of a secY mutant |
| 81.3341 | 523.303 | 0.15542437 | -2.68572 | 302.31875 | 8.239926649 | b1414 | *ydcF* | DUF218 domain-containing protein YdcF |

| | | | | | | | | |
|---|---|---|---|---|---|---|---|---|
| 62.30142 | 400.644 | 0.15550302 | -2.68499 | 231.472944 | 7.854699761 | b3147 | *lpoA* | outer membrane lipoprotein - activator of MrcA activity |
| 119.8214 | 769.904 | 0.15563157 | -2.68379 | 444.862864 | 8.797216862 | b2294 | *yfbU* | UPF0304 family protein YfbU |
| 47.55303 | 303.897 | 0.15647727 | -2.67598 | 175.725212 | 7.457177384 | b1613 | *manA* | mannose-6-phosphate isomerase |
| 56.63152 | 360.281 | 0.15718693 | -2.66945 | 208.456456 | 7.70360224 | b1646 | *sodC* | superoxide dismutase (Cu-Zn) |
| 14.87313 | 94.4138 | 0.15753127 | -2.66629 | 54.6434895 | 5.771977711 | b4019 | *metH* | cobalamin-dependent methionine synthase |
| 159.912 | 1012.43 | 0.15794813 | -2.66248 | 586.172965 | 9.195182621 | b2551 | *glyA* | serine hydroxymethyltransferase |
| 825.056 | 5173.7 | 0.15947117 | -2.64863 | 2999.3781 | 11.55044768 | b2007 | *yeeX* | DUF496 domain-containing protein YeeX |
| 225.7205 | 1413.61 | 0.1596766 | -2.64678 | 819.665396 | 9.678891282 | b0726 | *sucA* | subunit of E1(0) component of 2-oxoglutarate dehydrogenase |
| 55.40244 | 346.652 | 0.15982167 | -2.64547 | 201.027018 | 7.651245603 | b2414 | *cysK* | O-acetylserine sulfhydrylase A |
| 21.22886 | 131.667 | 0.16123187 | -2.63279 | 76.4477684 | 6.256402484 | b1655 | *mepH* | peptidoglycan DD-endopeptidase MepH |
| 66.94784 | 414.94 | 0.16134333 | -2.63179 | 240.944034 | 7.912554269 | b3858 | *yihD* | conserved protein YihD |
| 70.86213 | 438.966 | 0.16142953 | -2.63102 | 254.914227 | 7.993868086 | b2154 | *yeiG* | S-formylglutathione hydrolase/S-lactoylglutathione hydrolase |
| 41.06754 | 253.864 | 0.16176988 | -2.62799 | 147.465742 | 7.204236024 | b0864 | *artP* | L-arginine ABC transporter ATP binding subunit |
| 110.0401 | 678.181 | 0.16225759 | -2.62364 | 394.110736 | 8.622457239 | b0673 | *metT* | tRNA-Met |
| 131.9439 | 812.12 | 0.16246859 | -2.62177 | 472.031748 | 8.882740085 | b3236 | *mdh* | malate dehydrogenase |
| 55.49124 | 340.924 | 0.1627671 | -2.61912 | 198.207719 | 7.630869335 | b0928 | *aspC* | aspartate aminotransferase |
| 42.08978 | 257.607 | 0.16338733 | -2.61363 | 149.848568 | 7.227361484 | b2802 | *fucI* | L-fucose isomerase |
| 18.91221 | 115.483 | 0.16376643 | -2.61029 | 67.1975175 | 6.070336031 | b1402 | *insD-2* | IS2 insertion element protein InsB |
| 14.66199 | 88.8853 | 0.16495409 | -2.59986 | 51.7736286 | 5.694145529 | b2805 | *fucR* | DNA-binding transcriptional activator FucR |
| 25.82864 | 156.511 | 0.1650278 | -2.59922 | 91.169746 | 6.510483251 | b3515 | *gadW* | DNA-binding transcriptional dual regulator GadW |
| 2300.977 | 13890.3 | 0.16565317 | -2.59376 | 8095.65413 | 12.98293194 | b4226 | *ppa* | inorganic pyrophosphatase |
| 76.91879 | 459.452 | 0.16741414 | -2.57851 | 268.185497 | 8.067087412 | b1138 | *ymfE* | e14 prophage; uncharacterized protein YmfE |
| 140.0111 | 833.553 | 0.16796903 | -2.57373 | 486.782096 | 8.927132296 | b3609 | *secB* | SecB chaperone |
| 31.97391 | 189.863 | 0.16840523 | -2.56999 | 110.918413 | 6.793355068 | b0435 | *bolA* | DNA-binding transcriptional dual regulator BolA |
| 8.704441 | 51.5523 | 0.16884681 | -2.56621 | 30.1283687 | 4.913050657 | b1686 | *menI* | 1,4-dihydroxy-2-naphthoyl-CoA hydrolase |
| 243.7672 | 1441.75 | 0.16907767 | -2.56424 | 842.757166 | 9.718973179 | b4014 | *aceB* | malate synthase A |
| 20.15152 | 118.418 | 0.17017236 | -2.55493 | 69.284918 | 6.114469434 | b1647 | *ydhF* | putative oxidoreductase |
| 33.53124 | 196.922 | 0.1702764 | -2.55405 | 115.226818 | 6.848332726 | b1283 | *osmB* | osmotically-inducible lipoprotein OsmB |
| 16.02162 | 93.8986 | 0.17062691 | -2.55108 | 54.9600902 | 5.780312467 | b2662 | *gabT* | 4-aminobutyrate aminotransferase GabT |
| 464.4506 | 2719.86 | 0.17076301 | -2.54993 | 1592.15285 | 10.63676313 | b4602 | *ynhF* | stress response membrane protein YnhF |
| 80.63931 | 470.343 | 0.17144779 | -2.54416 | 275.491276 | 8.105862822 | b3284 | *smg* | DUF494 domain-containing protein Smg |
| 56.12859 | 326.704 | 0.17180256 | -2.54118 | 191.416318 | 7.580570009 | b0325 | *yahK* | aldehyde reductase, NADPH-dependent |
| 9.749685 | 56.7368 | 0.17184071 | -2.54086 | 33.2432224 | 5.054988332 | b3059 | *plsY* | putative glycerol-3-phosphate acyltransferase |
| 64.26243 | 371.744 | 0.17286755 | -2.53226 | 218.003086 | 7.768204749 | b1247 | *oppF* | murein tripeptide ABC transporter/oligopeptide ABC transporter ATP binding subunit |
| 66.50638 | 384.327 | 0.17304652 | -2.53077 | 225.416501 | 7.816449321 | b1757 | *ynjE* | molybdopterin synthase sulfurtransferase |
| 5.533315 | 31.9592 | 0.17313693 | -2.53001 | 18.7462491 | 4.228530051 | b2870 | *ygeW* | putative carbamoyltransferase YgeW |
| 52.33957 | 300.652 | 0.17408709 | -2.52212 | 176.495626 | 7.463488619 | b2133 | *dld* | D-lactate dehydrogenase |
| 24.0496 | 137.854 | 0.17445746 | -2.51905 | 80.9516422 | 6.338988443 | b0384 | *psiF* | PsiF family protein |
| 21.43777 | 122.688 | 0.174734 | -2.51677 | 72.062909 | 6.171184986 | b1326 | *mpaA* | murein tripeptide amidase A |
| 48.4145 | 276.73 | 0.17495224 | -2.51497 | 162.572171 | 7.344936505 | b0871 | *poxB* | pyruvate oxidase |
| 127.1843 | 722.686 | 0.17598836 | -2.50645 | 424.935156 | 8.731098898 | b1245 | *oppC* | murein tripeptide ABC transporter/oligopeptide ABC transporter inner membrane subunit |
| 8.621374 | 48.7649 | 0.17679475 | -2.49985 | 28.6931233 | 4.842633112 | b4307 | *yjhQ* | KpLE2 phage-like element; putative acetyltransferase TopAI antitoxin YjhQ |

| | | | | | | | | |
|---|---|---|---|---|---|---|---|---|
| 56.69821 | 320.386 | 0.17696865 | -2.49843 | 188.541915 | 7.558741475 | b4055 | *aphA* | acid phosphatase/phosphotransferase |
| 11.56819 | 65.1976 | 0.17743281 | -2.49466 | 38.3828956 | 5.262391646 | b3479 | *nikD* | Ni(2(+)) ABC transporter ATP binding subunit NikD |
| 53.79737 | 301.594 | 0.1783766 | -2.487 | 177.695852 | 7.473266191 | b1104 | *ycfL* | DUF1425 domain-containing protein YcfL |
| 9.479171 | 53.0619 | 0.17864371 | -2.48484 | 31.2705242 | 4.966731497 | b2342 | *fadI* | 3-ketoacyl-CoA thiolase FadI |
| 57.15228 | 319.544 | 0.17885583 | -2.48313 | 188.348079 | 7.557257506 | b1234 | *rssA* | putative patatin-like phospholipase RssA |
| 137.4236 | 764.111 | 0.17984766 | -2.47515 | 450.767259 | 8.81623892 | b0238 | *gpt* | xanthine-guanine phsophoribosyltransferase |
| 28.01946 | 155.696 | 0.17996311 | -2.47423 | 91.8575289 | 6.521326069 | b0953 | *rmf* | ribosome modulation factor |
| 37.07817 | 205.966 | 0.18002079 | -2.47376 | 121.522094 | 6.925074828 | b0655 | *gltI* | glutamate/aspartate ABC transporter periplasmic binding protein |
| 24.02064 | 132.269 | 0.18160439 | -2.46113 | 78.1448334 | 6.288078586 | b2276 | *nuoN* | NADH:quinone oxidoreductase subunit N |
| 77.05272 | 423.281 | 0.1820369 | -2.4577 | 250.16674 | 7.966746182 | b2091 | *gatD* | galactitol-1-phosphate 5-dehydrogenase |
| 133.2845 | 730.003 | 0.18258076 | -2.45339 | 431.643735 | 8.75369724 | b2897 | *sdhE* | FAD assembly factor |
| 13.02905 | 71.3605 | 0.18258076 | -2.45339 | 42.1947657 | 5.398992136 | b0892 | *rarA* | recombination factor |
| 14.33185 | 78.2499 | 0.18315491 | -2.44886 | 46.2908506 | 5.532655167 | b0753 | *ybgS* | PF13985 family protein YbgS |
| 11.28334 | 61.331 | 0.18397451 | -2.44242 | 36.3071807 | 5.182183004 | b1920 | *tcyJ* | cystine ABC transporter periplasmic binding protein |
| 13.78192 | 74.8348 | 0.18416448 | -2.44093 | 44.3083743 | 5.469507489 | b2714 | *ascG* | DNA-binding transcriptional repressor AscG |
| 23.93144 | 129.275 | 0.18512101 | -2.43346 | 76.6030204 | 6.259329372 | b3478 | *nikC* | Ni(2(+)) ABC transporter membrane subunit NikC |
| 6.864769 | 37.0405 | 0.18533151 | -2.43182 | 21.9526264 | 4.456321645 | b1745 | *astB* | N-succinylarginine dihydrolase |
| 35.86491 | 192.862 | 0.18596188 | -2.42692 | 114.363276 | 6.837480047 | b4751 | *yoaL* | protein YoaL |
| 26.54654 | 142.658 | 0.18608496 | -2.42597 | 84.6023485 | 6.402625807 | b3753 | *rbsR* | DNA-binding transcriptional dual regulator RbsR |
| 337.3543 | 1806.37 | 0.18675772 | -2.42076 | 1071.86435 | 10.06590662 | b3560 | *glyQ* | glycine--tRNA ligase subunit alpha |
| 16.31854 | 87.3458 | 0.18682682 | -2.42023 | 51.832161 | 5.69577564 | b2291 | *yfbR* | dCMP phosphohydrolase |
| 449.2181 | 2404.2 | 0.18684729 | -2.42007 | 1426.70871 | 10.4784751 | b1508 | *hipB* | antitoxin/DNA-binding transcriptional repressor HipB |
| 24.9542 | 132.91 | 0.18775302 | -2.41309 | 78.9319636 | 6.302537734 | b0228 | *rayT* | REP-associated tyrosine transposase |
| 132.02 | 702.495 | 0.18793019 | -2.41173 | 417.257405 | 8.704793843 | b1853 | *yebK* | DNA-binding transcriptional repressor YebK |
| 75.83935 | 401.051 | 0.1891015 | -2.40277 | 238.445183 | 7.897513826 | b4442 | *micA* | small regulatory RNA MicA |
| 46.14971 | 243.572 | 0.1894706 | -2.39995 | 144.86078 | 7.178523239 | b3830 | *ysgA* | putative dienelactone hydrolase |
| 18.58364 | 97.9422 | 0.18974079 | -2.3979 | 58.2629315 | 5.864506387 | b1726 | *yniB* | uncharacterized protein YniB |
| 29.63837 | 155.386 | 0.19074079 | -2.39031 | 92.5119666 | 6.531568088 | b1481 | *bdm* | biofilm-dependent modulation protein |
| 20.48075 | 106.252 | 0.19275554 | -2.37516 | 63.3666152 | 5.98565105 | b1590 | *ynfH* | putative menaquinol dehydrogenase |
| 46.03976 | 237.76 | 0.19363942 | -2.36856 | 141.900001 | 7.148730785 | b4227 | *ytfQ* | galactofuranose ABC transporter periplasmic binding protein |
| 26.79794 | 138.346 | 0.19370243 | -2.36809 | 82.5719315 | 6.367579548 | b4149 | *blc* | outer membrane lipoprotein Blc |
| 181.7529 | 936.437 | 0.19408991 | -2.3652 | 559.094842 | 9.126949225 | b3611 | *yibN* | putative sulfurtransferase YibN |
| 145.4887 | 748.584 | 0.19435189 | -2.36326 | 447.036338 | 8.804248296 | b2898 | *ygfZ* | folate-binding protein |
| 751.5539 | 3856.26 | 0.19489192 | -2.35925 | 2303.90682 | 11.16986665 | b3314 | *rpsC* | 30S ribosomal subunit protein S3 |
| 17.33936 | 88.9491 | 0.19493585 | -2.35893 | 53.1442091 | 5.73184059 | b2502 | *ppx* | exopolyphosphatase |
| 8.47441 | 43.3818 | 0.19534476 | -2.35591 | 25.9281115 | 4.696445226 | b0944 | *ycbF* | putative fimbrial chaperone YcbF |
| 216.4262 | 1100.96 | 0.19657961 | -2.34681 | 658.692889 | 9.363462166 | b0226 | *dinJ* | antitoxin/DNA-binding transcriptional repressor DinJ |
| 22.43308 | 114.016 | 0.19675325 | -2.34554 | 68.2247066 | 6.09222238 | b1492 | *gadC* | L-glutamate:4-aminobutyrate antiporter |
| 9.502939 | 48.1965 | 0.19717055 | -2.34248 | 28.8497427 | 4.850486548 | b3624 | *waaZ* | lipopolysaccharide core biosynthesis protein WaaZ |
| 30.03043 | 151.319 | 0.19845735 | -2.3331 | 90.6748737 | 6.502630925 | b3468 | *yhhN* | conserved inner membrane enzyme YhhN |
| 62.39331 | 313.521 | 0.1990082 | -2.3291 | 187.957317 | 7.554261267 | b0759 | *galE* | UDP-glucose 4-epimerase |
| 7.105313 | 35.673 | 0.19917901 | -2.32786 | 21.3891573 | 4.418807737 | b1512 | *lsrR* | DNA-binding transcriptional repressor LsrR |

| | | | | | | | | |
|---|---|---|---|---|---|---|---|---|
| 195.556 | 977.314 | 0.20009545 | -2.32124 | 586.434909 | 9.195827176 | b2330 | *prmB* | 50S ribosomal subunit protein L3 N(5)-glutamine methyltransferase |
| 11.71934 | 58.352 | 0.20083884 | -2.31589 | 35.0356635 | 5.130752313 | b0213 | *yafS* | putative S-adenosyl-L-methionine-dependent methyltransferase |
| 97.93286 | 485.195 | 0.20184247 | -2.3087 | 291.563679 | 8.1876672 | b0410 | *yajD* | HNH nuclease family protein |
| 69.76326 | 345.21 | 0.20208922 | -2.30694 | 207.486721 | 7.696875199 | b1329 | *mppA* | murein tripeptide ABC transporter periplasmic binding protein |
| 26.18913 | 129.05 | 0.2029382 | -2.30089 | 77.6194494 | 6.278346295 | b2949 | *yqgF* | ribonuclease H-like domain containing nuclease |
| 41.5041 | 204.393 | 0.20306021 | -2.30002 | 122.948581 | 6.941911274 | b1378 | *ydbK* | putative pyruvate-flavodoxin oxidoreductase |
| 112.9456 | 548.112 | 0.20606297 | -2.27884 | 330.528706 | 8.368631762 | b4138 | *dcuA* | C4-dicarboxylate transporter DcuA |
| 194.0844 | 940.887 | 0.20627801 | -2.27734 | 567.48584 | 9.148440584 | b0721 | *sdhC* | succinate:quinone oxidoreductase, membrane protein SdhC |
| 201.4712 | 976.429 | 0.20633467 | -2.27694 | 588.950083 | 9.202001553 | b3920 | *yiiQ* | DUF1454 domain-containing protein YiiQ |
| 31.65311 | 151.12 | 0.20945647 | -2.25528 | 91.3866749 | 6.513911915 | b1624 | *ydgJ* | putative oxidoreductase YdgJ |
| 42.33519 | 200.955 | 0.21067011 | -2.24694 | 121.645044 | 6.92653374 | b0281 | *intF* | CP4-6 prophage; putative phage integrase |
| 250.1827 | 1186.66 | 0.21082936 | -2.24585 | 718.421144 | 9.488686001 | b1064 | *grxB* | reduced glutaredoxin 2 |
| 7.654097 | 36.3004 | 0.21085442 | -2.24568 | 21.9772432 | 4.457938523 | b3541 | *dppD* | dipeptide ABC transporter ATP binding subunit DppD |
| 77.3956 | 365.998 | 0.21146467 | -2.24151 | 221.6967 | 7.792443486 | b2282 | *nuoH* | NADH:quinone oxidoreductase subunit H |
| 44.15346 | 207.309 | 0.21298422 | -2.23118 | 125.731037 | 6.974197018 | b4046 | *zur* | DNA-binding transcriptional repressor Zur |
| 39.96376 | 187.122 | 0.21357095 | -2.22721 | 113.542735 | 6.827091591 | b3496 | *dtpB* | dipeptide/tripeptide:H(+) symporter DtpB |
| 32.70036 | 153.037 | 0.21367667 | -2.2265 | 92.8685093 | 6.537117572 | b2751 | *cysN* | sulfate adenylyltransferase subunit 1 |
| 13.44934 | 62.3154 | 0.21582681 | -2.21205 | 37.8823918 | 5.243455515 | b3160 | *yhbW* | putative luciferase-like monooxygenase |
| 22.37494 | 103.571 | 0.21603448 | -2.21067 | 62.9730422 | 5.976662459 | b3566 | *xylF* | xylose ABC transporter periplasmic binding protein |
| 300.5411 | 1390.66 | 0.21611439 | -2.21013 | 845.599251 | 9.723830288 | b0250 | *ykfB* | CP4-6 prophage; protein YkfB |
| 28.47781 | 131.533 | 0.21650711 | -2.20751 | 80.0053647 | 6.322024836 | b3540 | *dppF* | dipeptide ABC transporter ATP binding subunit DppF |
| 9.71654 | 44.8494 | 0.21664831 | -2.20657 | 27.2829562 | 4.769928068 | b0862 | *artQ* | L-arginine ABC transporter membrane subunit ArtQ |
| 406.8813 | 1875.76 | 0.21691517 | -2.2048 | 1141.32169 | 10.15648977 | b0382 | *iraP* | anti-adaptor protein for sigma(S) stabilization |
| 13.35396 | 61.5516 | 0.21695553 | -2.20453 | 37.4527721 | 5.2270006 | b1919 | *dcyD* | D-cysteine desulfhydrase |
| 17.41184 | 80.143 | 0.21725962 | -2.20251 | 48.7774173 | 5.608141466 | b3211 | *yhcC* | radical SAM family oxidoreductase YhcC |
| 10.97809 | 50.3545 | 0.21801584 | -2.1975 | 30.6663081 | 4.938582586 | b2380 | *ypdA* | sensory histidine kinase YpdA |
| 331.3918 | 1518.79 | 0.21819528 | -2.19631 | 925.088606 | 9.853447744 | b3978 | *glyT* | tRNA-Gly |
| 169.4008 | 776.186 | 0.21824762 | -2.19596 | 472.79348 | 8.88506633 | b2147 | *preA* | NAD-dependent dihydropyrimidine dehydrogenase subunit PreA |
| 32.11423 | 146.716 | 0.21888745 | -2.19174 | 89.4149739 | 6.482444547 | b2264 | *menD* | 2-succinyl-5-enolpyruvyl-6-hydroxy-3- cyclohexene-1-carboxylate synthase |
| 192.4898 | 879.092 | 0.21896436 | -2.19123 | 535.790784 | 9.065525955 | b1208 | *ispE* | 4-(cytidine 5'-diphospho)-2-C-methyl-D-erythritol kinase |
| 139.4747 | 631.959 | 0.22070202 | -2.17983 | 385.716921 | 8.591398627 | b0050 | *apaG* | DUF525 domain-containing protein ApaG |
| 16.84929 | 76.3441 | 0.22070202 | -2.17983 | 46.596675 | 5.542155108 | b1789 | *yeaL* | conserved inner membrane protein YeaL |
| 378.6062 | 1713.25 | 0.22098772 | -2.17796 | 1045.92591 | 10.03056494 | b2696 | *csrA* | carbon storage regulator |
| 18.41567 | 83.3197 | 0.22102421 | -2.17772 | 50.8676744 | 5.668677232 | b2465 | *tktB* | transketolase 2 |
| 17.55766 | 79.3829 | 0.22117695 | -2.17673 | 48.4702712 | 5.599028249 | b3060 | *ttdR* | DNA-binding transcriptional activator Dan |
| 98.38618 | 444.436 | 0.2213731 | -2.17545 | 271.41111 | 8.084335965 | b4216 | *ytfJ* | protein YtfJ |
| 196.8011 | 888.792 | 0.22142541 | -2.17511 | 542.796511 | 9.084267638 | b4144 | *yjeI* | DUF4156 domain-containing lipoprotein YjeI |
| 39.06266 | 175.922 | 0.22204542 | -2.17107 | 107.492296 | 6.748089462 | b4461 | *yfjD* | putative inner membrane protein |
| 38.97648 | 175.232 | 0.22242737 | -2.16859 | 107.104439 | 6.74287446 | b1638 | *pdxH* | pyridoxal 5-phosphate synthase |
| 69.71462 | 313.22 | 0.22257408 | -2.16764 | 191.46723 | 7.580953681 | b2278 | *nuoL* | NADH:quinone oxidoreductase subunit L |
| 111.4216 | 500.141 | 0.22278052 | -2.16631 | 305.781045 | 8.256355169 | b1982 | *amn* | AMP nucleosidase |
| 300.7762 | 1341.75 | 0.22416676 | -2.15736 | 821.264134 | 9.681702485 | b3796 | *argX* | tRNA-Arg |

| | | | | | | | | |
|---|---|---|---|---|---|---|---|---|
| 16.42152 | 73.1031 | 0.22463491 | -2.15435 | 44.7623318 | 5.484213288 | b2789 | *gudP* | galactarate/glucarate/glycerate transporter GudP |
| 11.91346 | 52.9526 | 0.22498339 | -2.15211 | 32.4330422 | 5.019392447 | b2800 | *fucA* | L-fuculose-phosphate aldolase |
| 15.01843 | 66.7426 | 0.2250201 | -2.15187 | 40.88053 | 5.353341996 | b3538 | *bcsG* | cellulose phosphoethanolamine transferase |
| 380.5352 | 1678.68 | 0.22668705 | -2.14123 | 1029.60804 | 10.00787951 | b3831 | *udp* | uridine phosphorylase |
| 48.26557 | 212.45 | 0.22718544 | -2.13806 | 130.357852 | 7.02633367 | b1794 | *dgcP* | diguanylate cyclase DgcP |
| 135.9212 | 596.351 | 0.22792124 | -2.13339 | 366.136326 | 8.516237108 | b3237 | *argR* | DNA-binding transcriptional dual regulator ArgR |
| 116.705 | 510.178 | 0.2287537 | -2.12813 | 313.441281 | 8.292051389 | b0811 | *glnH* | L-glutamine ABC transporter periplasmic binding protein |
| 58.50844 | 255.17 | 0.22929187 | -2.12474 | 156.839292 | 7.293143222 | b3810 | *yigA* | conserved protein YigA |
| 4042.022 | 17621.8 | 0.22937633 | -2.12421 | 10831.9069 | 13.40299962 | b1243 | *oppA* | oligopeptide ABC transporter periplasmic binding protein |
| 14.69928 | 64.0117 | 0.22963424 | -2.12259 | 39.3554985 | 5.298493308 | b1290 | *sapF* | putrescine ABC exporter ATP binding protein SapF |
| 23.6882 | 102.724 | 0.23060004 | -2.11654 | 63.2061831 | 5.98199379 | b1610 | *tus* | DNA replication terminus site-binding protein |
| 62.99183 | 272.165 | 0.23144684 | -2.11125 | 167.578633 | 7.388694402 | b4025 | *pgi* | glucose-6-phosphate isomerase |
| 7.369006 | 31.7568 | 0.23204487 | -2.10752 | 19.5629094 | 4.290049039 | b4214 | *cysQ* | 3'(2'),5'-bisphosphate nucleotidase |
| 712.8259 | 3053.65 | 0.2334338 | -2.09891 | 1883.23962 | 10.87900086 | b1136 | *icd* | isocitrate dehydrogenase |
| 37.84276 | 161.768 | 0.2339316 | -2.09584 | 99.8056177 | 6.641049117 | b4353 | *yjiX* | conserved protein YjiX |
| 2084.225 | 8905.18 | 0.23404638 | -2.09513 | 5494.70263 | 12.42382569 | b0008 | *talB* | transaldolase B |
| 9.353007 | 39.9519 | 0.23410662 | -2.09476 | 24.6524594 | 4.623659674 | b0909 | *ycaL* | periplasmic protease YcaL |
| 15.2915 | 64.8879 | 0.23566028 | -2.08522 | 40.0896857 | 5.325159202 | b3809 | *dapF* | diaminopimelate epimerase |
| 21.37394 | 90.5132 | 0.23614161 | -2.08228 | 55.9435757 | 5.805900562 | b1834 | *yebT* | intermembrane transport protein YebT |
| 9.9588 | 42.1481 | 0.23628098 | -2.08142 | 26.0534613 | 4.70340315 | b2875 | *yqeB* | XdhC-CoxI family protein YqeB |
| 94.2874 | 397.634 | 0.23712124 | -2.0763 | 245.960551 | 7.942283134 | b2908 | *pepP* | proline aminopeptidase P II |
| 8.944571 | 37.214 | 0.24035479 | -2.05676 | 23.0793014 | 4.528527647 | b2622 | *intA* | CP4-57 prophage; integrase |
| 28.63589 | 118.948 | 0.24074277 | -2.05444 | 73.7919888 | 6.205392295 | b2703 | *srlE* | sorbitol-specific PTS enzyme IIBC1 component |
| 10.60793 | 43.795 | 0.24221769 | -2.04562 | 27.2014837 | 4.765613438 | b2251 | *nudI* | pyrimidine deoxynucleotide diphosphatase NudI |
| 78.65312 | 324.375 | 0.24247616 | -2.04409 | 201.513884 | 7.654735434 | b3410 | *feoC* | ferrous iron transport protein FeoC |
| 338.5153 | 1393.53 | 0.24291926 | -2.04145 | 866.022788 | 9.758261178 | b1241 | *adhE* | aldehyde-alcohol dehydrogenase |
| 21.99961 | 90.5305 | 0.24300784 | -2.04093 | 56.2650366 | 5.814166796 | b3555 | *yiaG* | putative DNA-binding transcriptional regulator YiaG |
| 20.4842 | 84.1648 | 0.24338201 | -2.03871 | 52.3245123 | 5.709415053 | b2132 | *bglX* | beta-D-glucoside glucohydrolase, periplasmic |
| 1323.18 | 5420.21 | 0.24411979 | -2.03434 | 3371.69423 | 11.71925799 | b2911 | *ssrS* | 6S RNA |
| 14.51408 | 59.3764 | 0.244442 | -2.03244 | 36.9452333 | 5.207316333 | b2197 | *ccmE* | periplasmic heme chaperone |
| 25.64758 | 104.654 | 0.2450712 | -2.02873 | 65.1505931 | 6.025706408 | b3893 | *fdoH* | formate dehydrogenase O subunit beta |
| 214.7996 | 875.894 | 0.24523479 | -2.02776 | 545.346757 | 9.091030044 | b0828 | *iaaA* | isoaspartyl dipeptidase proenzyme |
| 43.05875 | 175.513 | 0.24533108 | -2.0272 | 109.28578 | 6.771961889 | b4167 | *nnr* | NAD(P)HX epimerase/NAD(P)HX dehydratase |
| 12.05061 | 48.8761 | 0.24655407 | -2.02002 | 30.4633752 | 4.929003892 | b0826 | *moeB* | molybdopterin-synthase adenylyltransferase |
| 24.56865 | 99.2005 | 0.24766654 | -2.01353 | 61.8845746 | 5.951507941 | b0425 | *panE* | 2-dehydropantoate 2-reductase |
| 39.01211 | 157.396 | 0.24785918 | -2.01241 | 98.2041873 | 6.617712635 | b3835 | *ubiB* | ubiquinone biosynthesis protein UbiB |
| 77.82686 | 312.924 | 0.24870854 | -2.00747 | 195.375412 | 7.610105108 | b0880 | *cspD* | DNA replication inhibitor |
| 118.7419 | 477.207 | 0.24882688 | -2.00679 | 297.974508 | 8.2190451 | b3688 | *yidQ* | DUF1375 domain-containing putative lipoprotein YidQ |
| 12.05061 | 48.4178 | 0.24888806 | -2.00643 | 30.2342032 | 4.918109655 | b4473 | *smf* | protein Smf |
| 54.68512 | 219.529 | 0.24910243 | -2.00519 | 137.106874 | 7.099157098 | b0472 | *recR* | DNA repair protein RecR |
| 26.32022 | 105.345 | 0.24984736 | -2.00088 | 65.8327084 | 6.040732648 | b2785 | *rlmD* | 23S rRNA m(5)U1939 methyltransferase |

**Table S3. List of deletion mutants from KEIO collection showing decreased sensitivity to TAT-RasGAP$_{317-326}$.** Mutants were determined as less sensitive when the $NG^{m}_{6\ hours}$ (P) was higher than 0.719 (the mean $NG^{WT}_{6hours}$ (P) + 2 SDs ). Results of the second strain of the collection having the same gene deleted are presented for comparison (Replicate 2).

| Locus_ tag | Gene_ name | Product | Sample_ID Replicate 1 | $NG^{m}_{24\ hours}$(NoP) Replicate 1 | $NG^{m}_{6\ hours}$(P) Replicate 1 | Sample_ID Replicate 2 | $NG^{m}_{24\ hours}$(NoP) Replicate 2 | $NG^{m}_{6\ hours}$(P) Replicate 2 |
|---|---|---|---|---|---|---|---|---|
| b1338 | *abgA* | p-aminobenzoyl-glutamate hydrolase subunit A | KEIO_75_F10 | 1.17 | 0.75 | KEIO_76_F10 | 1.18 | 0.34 |
| b2195 | *ccmG* | holocytochrome c synthetase - thiol:disulfide oxidoreductase CcmG | KEIO_46_E11 | 0.86 | 0.75 | KEIO_45_E11 | 1.16 | 0.08 |
| b1731 | *cedA* | cell division modulator | KEIO_46_F10 | 0.81 | 0.75 | KEIO_45_F10 | 1.17 | 0.03 |
| b2417 | *crr* | Enzyme IIA(Glc) | KEIO_58_H8 | 0.95 | 0.75 | KEIO_57_H8 | 1.10 | 0.70 |
| b3806 | *cyaA* | adenylate cyclase | KEIO_53_F10 | 0.63 | 0.95 | KEIO_54_F10 | 0.79 | 0.30 |
| b0014 | *dnaK* | chaperone protein DnaK | KEIO_46_E8 | 0.79 | 0.73 | KEIO_45_E8 | 1.01 | -0.05 |
| b0592 | *fepB* | ferric enterobactin ABC transporter periplasmic binding protein | KEIO_53_F11 | 0.85 | 0.74 | KEIO_54_F11 | 0.91 | -0.03 |
| b0809 | *glnQ* | L-glutamine ABC transporter ATP binding subunit | KEIO_55_E1 | 1.20 | 0.73 | KEIO_56_E1 | 1.04 | 0.23 |
| b1132 | *hflD* | lysogenization regulator | KEIO_89_B12 | 0.67 | 0.72 | KEIO_90_B12 | 0.75 | 0.59 |
| b2830 | *nudH* | RNA pyrophosphohydrolase | KEIO_24_H2 | 1.01 | 0.76 | KEIO_23_H2 | 0.94 | 0.23 |
| b2285 | *nuoE* | NADH:quinone oxidoreductase subunit E | KEIO_04_H4 | 1.38 | 0.85 | KEIO_03_H4 | 0.90 | 0.05 |
| b2277 | *nuoM* | NADH:quinone oxidoreductase subunit M | KEIO_04_H3 | 1.49 | 0.89 | KEIO_03_H3 | 0.90 | 0.03 |
| b3167 | *rbfA* | 30S ribosome binding factor | KEIO_85_C8 | 1.03 | 0.79 | KEIO_86_C8 | 1.33 | 0.02 |
| b3631 | *rfaG* | lipopolysaccharide glucosyltransferase I | KEIO_46_C6 | 0.68 | 0.77 | KEIO_45_C6 | 1.06 | 0.27 |
| b3842 | *rfaH* | transcription antiterminator RfaH | KEIO_46_E7 | 0.92 | 0.93 | KEIO_45_E7 | 0.85 | 0.70 |
| b3630 | *rfaP* | lipopolysaccharide core heptose (I) kinase | KEIO_46_B6 | 0.60 | 0.87 | KEIO_45_B6 | 1.21 | 0.52 |
| b3632 | *rfaQ* | lipopolysaccharide core heptosyltransferase 3 | KEIO_46_D6 | 0.78 | 0.76 | KEIO_45_D6 | 1.19 | 0.20 |
| b3625 | *rfaY* | lipopolysaccharide core heptose (II) kinase | KEIO_46_F5 | 1.04 | 0.75 | KEIO_45_F5 | 1.27 | 0.49 |
| b3179 | *rrmJ* | 23S rRNA 2'-O-ribose U2552 methyltransferase | KEIO_46_E12 | 0.70 | 0.89 | KEIO_45_E12 | 1.14 | 0.26 |
| b4161 | *rsgA* | ribosome small subunit-dependent GTPase A | KEIO_63_D11 | 0.99 | 0.85 | KEIO_64_D11 | 0.70 | 0.47 |
| b3833 | *ubiE* | adenosylmethionine:2-DMK methyltransferase | KEIO_44_D8 | 0.60 | 0.76 | KEIO_43_D8 | 0.56 | 0.17 |
| b2311 | *ubiX* | flavin prenyltransferase | KEIO_44_E6 | 0.53 | 0.77 | KEIO_43_E6 | 1.22 | -0.01 |
| b0660 | *ybeZ* | PhoH-like protein | KEIO_66_E4 | 0.72 | 0.72 | KEIO_65_E4 | 1.03 | 0.40 |
| b3343 | *yheL* | sulfurtransferase complex subunit TusB | KEIO_36_E4 | 0.91 | 0.78 | KEIO_35_E4 | 1.23 | -0.01 |
| b3448 | *yhhA* | DUF2756 domain-containing protein YhhA | KEIO_36_F7 | 0.82 | 0.78 | KEIO_35_F7 | 1.33 | -0.01 |
| b4056 | *yjbQ* | UPF0047 protein YjbQ | KEIO_38_F8 | 0.76 | 0.93 | KEIO_37_F8 | 1.30 | 0.00 |
| b1826 | *yobG* | PhoQ kinase inhibitor | KEIO_32_E12 | 0.87 | 0.89 | KEIO_31_E12 | 1.21 | 0.40 |

**Table S4. List of genes which deletion caused hypersensitivity to TAT-RasGAP$_{317-326}$ as determined by screening of the KEIO collection.** Mutants are selected as "hypersensitive" when the two individual mutants from the Keio collection (designed as replicate 1 and replicate 2 and for which their position in the collection is indicated by their Sample_ID) showed a normalized growth at 24h in presence of the peptide ($NG^{m}_{24\ hours}$(P) lower than 0.212 (Average NG of wild-type - 3 SD).

| Locus_tag | Gene_ name | Product | Sample_ID Replicate 1 | $NG^{m}_{24\ hours}$(NoP) Replicate 1 | $NG^{m}_{24\ hours}$(P) Replicate 1 | Sample_ID Replicate 2 | $NG^{m}_{24\ hours}$(NoP) Replicate 2 | $NG^{m}_{24\ hours}$(P) Replicate2 |
|---|---|---|---|---|---|---|---|---|
| b0118 | *acnB* | hypothetical protein | KEIO_01_B11 | 0.765 | 0.025 | KEIO_02_B11 | 0.778 | 0.173 |
| b2916 | *argP* | DNA-binding transcriptional dual regulator ArgP | KEIO_01_B5 | 1.015 | -0.017 | KEIO_02_B5 | 1.006 | -0.008 |
| b3237 | *argR* | DNA-binding transcriptional dual regulator ArgR | KEIO_01_F1 | 1.079 | 0.204 | KEIO_02_F1 | 1.075 | -0.088 |
| b2078 | *baeS* | sensory histidine kinase BaeS | KEIO_03_F11 | 0.845 | -0.004 | KEIO_04_F11 | 1.273 | 0.041 |
| b3723 | *bglG* | transcriptional antiterminator BglG | KEIO_01_A2 | 0.876 | -0.034 | KEIO_02_A2 | 0.923 | -0.011 |
| b1735 | *chbR* | DNA-binding transcriptional dual regulator ChbR | KEIO_01_E2 | 0.817 | 0.000 | KEIO_02_E2 | 0.835 | 0.009 |
| b0620 | *citB* | DNA-binding transcriptional dual regulator DpiA | KEIO_03_C9 | 0.993 | 0.011 | KEIO_04_C9 | 1.186 | 0.090 |
| b0103 | *coaE* | dephospho-CoA kinase | KEIO_05_A9 | 1.024 | 0.008 | KEIO_06_A9 | 1.086 | 0.068 |
| b3911 | *cpxA* | sensory histidine kinase CpxA | KEIO_05_D2 | 0.642 | -0.021 | KEIO_06_D2 | 1.191 | 0.000 |
| b4398 | *creB* | DNA-binding transcriptional regulator CreB | KEIO_05_D3 | 0.762 | -0.012 | KEIO_06_D3 | 1.339 | 0.201 |
| b4399 | *creC* | sensory histidine kinase CreC | KEIO_05_E3 | 0.727 | -0.023 | KEIO_06_E3 | 1.259 | 0.027 |
| b0338 | *cynR* | DNA-binding transcriptional dual regulator CynR | KEIO_01_H2 | 0.923 | 0.055 | KEIO_02_H2 | 0.948 | 0.044 |
| b0430 | *cyoC* | cytochrome bo3 ubiquinol oxidase subunit 3 | KEIO_01_E11 | 0.781 | 0.000 | KEIO_02_E11 | 0.833 | 0.099 |
| b0429 | *cyoD* | cytochrome bo3 ubiquinol oxidase subunit 4 | KEIO_01_D11 | 0.824 | 0.018 | KEIO_02_D11 | 0.769 | -0.009 |
| b4124 | *dcuR* | DNA-binding transcriptional activator DcuR | KEIO_05_B3 | 0.800 | 0.023 | KEIO_06_B3 | 1.323 | 0.089 |
| b4125 | *dcuS* | sensory histidine kinase DcuS | KEIO_05_C3 | 0.772 | -0.004 | KEIO_06_C3 | 1.372 | 0.105 |
| b2133 | *dld* | D-lactate dehydrogenase | KEIO_05_C5 | 1.025 | 0.017 | KEIO_06_C5 | 1.279 | 0.194 |
| b2364 | *dsdC* | DNA-binding transcriptional dual regulator DsdC | KEIO_01_C3 | 0.814 | 0.077 | KEIO_02_C3 | 0.864 | 0.008 |
| b3264 | *envR* | DNA-binding transcriptional repressor EnvR | KEIO_01_E3 | 0.824 | 0.003 | KEIO_02_E3 | 0.860 | 0.072 |
| b0566 | *envY* | DNA-binding transcriptional activator EnvY | KEIO_01_F3 | 0.767 | -0.028 | KEIO_02_F3 | 0.760 | 0.027 |
| b3404 | *envZ* | sensory histidine kinase EnvZ | KEIO_05_B10 | 0.780 | -0.007 | KEIO_06_B10 | 1.254 | 0.018 |
| b1187 | *fadR* | DNA-binding transcriptional dual regulator FadR | KEIO_01_G3 | 0.824 | 0.198 | KEIO_02_G3 | 0.812 | 0.042 |
| b0588 | *fepC* | ferric enterobactin ABC transporter ATP binding subunit | KEIO_41_B2 | 0.926 | -0.009 | KEIO_42_B2 | 0.797 | -0.017 |
| b0585 | *fes* | enterochelin esterase | KEIO_41_H1 | 1.103 | 0.021 | KEIO_42_H1 | 0.995 | 0.073 |
| b0150 | *fhuA* | ferrichrome outer membrane transporter/phage receptor | KEIO_05_G6 | 0.795 | -0.001 | KEIO_06_G6 | 1.245 | 0.029 |

| | | | | | | | | |
|---|---|---|---|---|---|---|---|---|
| b4152 | *frdC* | fumarate reductase membrane protein FrdC | KEIO_03_B8 | 0.840 | 0.003 | KEIO_04_B8 | 1.135 | 0.157 |
| b1612 | *fumA* | fumarase A | KEIO_03_G2 | 0.814 | 0.021 | KEIO_04_G2 | 1.097 | 0.046 |
| b1416 | *gapC* | | KEIO_03_C2 | 0.806 | 0.035 | KEIO_03_D2 | 0.857 | 0.007 |
| b2388 | *glk* | glucokinase | KEIO_03_C5 | 1.030 | -0.015 | KEIO_04_C5 | 1.283 | -0.010 |
| b0720 | *gltA* | citrate synthase | KEIO_05_F5 | 0.901 | 0.030 | KEIO_06_F5 | 1.155 | -0.014 |
| b3612 | *gpmI* | 2%2C3-bisphosphoglycerate-independent phosphoglycerate | KEIO_03_E6 | 0.859 | 0.024 | KEIO_04_E6 | 1.298 | 0.066 |
| b4000 | *hupA* | DNA-binding protein HU-alpha | KEIO_01_H4 | 1.019 | 0.136 | KEIO_02_H4 | 1.030 | 0.055 |
| b4264 | *idnR* | DNA-binding transcriptional dual regulator IdnR | KEIO_01_C5 | 1.020 | 0.076 | KEIO_02_C5 | 0.989 | -0.028 |
| b0695 | *kdpD* | sensory histidine kinase KdpD | KEIO_03_E9 | 1.053 | -0.014 | KEIO_04_E9 | 1.278 | 0.000 |
| b0076 | *leuO* | DNA-binding transcriptional dual regulator LeuO | KEIO_01_E5 | 1.023 | 0.006 | KEIO_02_E5 | 1.004 | 0.117 |
| b0889 | *lrp* | DNA-binding transcriptional dual regulator Lrp | KEIO_01_F5 | 1.029 | 0.050 | KEIO_02_F5 | 1.101 | -0.033 |
| b2839 | *lysR* | DNA-binding transcriptional dual regulator LysR | KEIO_01_G5 | 1.051 | 0.009 | KEIO_02_G5 | 0.961 | -0.040 |
| b3938 | *metJ* | DNA-binding transcriptional repressor MetJ | KEIO_01_D6 | 0.799 | -0.050 | KEIO_02_D6 | 0.939 | -0.024 |
| b0346 | *mhpR* | DNA-binding transcriptional activator MhpR | KEIO_01_F6 | 0.808 | -0.039 | KEIO_02_F6 | 0.775 | -0.048 |
| b3226 | *nanR* | DNA-binding transcriptional dual regulator NanR | KEIO_01_B7 | 0.749 | 0.129 | KEIO_02_B7 | 0.757 | -0.051 |
| b0020 | *nhaR* | DNA-binding transcriptional activator NhaR | KEIO_01_C7 | 0.753 | 0.011 | KEIO_02_C7 | 0.812 | 0.011 |
| b2477 | *nlpB* | outer membrane protein assembly factor BamC | KEIO_07_D4 | 0.848 | -0.036 | KEIO_08_D4 | 1.122 | 0.030 |
| b2284 | *nuoF* | NADH:quinone oxidoreductase subunit F | KEIO_03_G4 | 0.741 | -0.008 | KEIO_04_G4 | 0.732 | 0.082 |
| b3405 | *ompR* | DNA-binding transcriptional dual regulator OmpR | KEIO_05_F1 | 0.918 | -0.020 | KEIO_06_F1 | 1.161 | -0.013 |
| b3403 | *pck* | phosphoenolpyruvate carboxykinase (ATP) | KEIO_03_D6 | 0.796 | 0.034 | KEIO_04_D6 | 1.120 | 0.181 |
| b0113 | *pdhR* | DNA-binding transcriptional dual regulator PdhR | KEIO_01_D7 | 0.724 | 0.135 | KEIO_02_D7 | 0.733 | -0.054 |
| b0903 | *pflB* | pyruvate formate-lyase | KEIO_03_D1 | 1.049 | -0.051 | KEIO_04_D1 | 1.239 | 0.083 |
| b4102 | *phnF* | putative transcriptional regulator PhnF | KEIO_01_F7 | 0.749 | -0.023 | KEIO_02_F7 | 0.750 | -0.044 |
| b1130 | *phoP* | DNA-binding transcriptional dual regulator PhoP | KEIO_03_B10 | 0.723 | -0.071 | KEIO_04_B10 | 1.173 | -0.005 |
| b1129 | *phoQ* | sensory histidine kinase PhoQ | KEIO_03_A10 | 0.931 | -0.040 | KEIO_04_A10 | 1.132 | -0.049 |
| b0333 | *prpC* | 2-methylcitrate synthase | KEIO_01_C11 | 0.797 | 0.053 | KEIO_02_C11 | 0.859 | -0.033 |
| b3753 | *rbsR* | DNA-binding transcriptional dual regulator RbsR | KEIO_01_E10 | 0.796 | 0.165 | KEIO_02_E10 | 0.790 | 0.009 |
| b2218 | *rcsC* | sensory histidine kinase RcsC | KEIO_01_A10 | 0.850 | 0.088 | KEIO_02_A10 | 0.790 | -0.086 |
| b2572 | *rseA* | anti-sigma factor | KEIO_07_C1 | 0.211 | -0.231 | KEIO_08_C1 | 0.200 | -0.295 |
| b0723 | *sdhA* | succinate:quinone oxidoreductase%2C FAD binding protein | KEIO_01_C12 | 0.921 | 0.053 | KEIO_02_C12 | 0.678 | 0.040 |
| b0724 | *sdhB* | succinate:quinone oxidoreductase%2C iron-sulfur cluster | KEIO_01_D12 | 0.694 | 0.074 | KEIO_02_D12 | 0.682 | 0.091 |
| b1916 | *sdiA* | DNA-binding transcriptional dual regulator SdiA | KEIO_01_F10 | 0.817 | -0.049 | KEIO_02_F10 | 0.771 | 0.029 |
| b4063 | *soxR* | DNA-binding transcriptional dual regulator SoxR | KEIO_01_C8 | 0.795 | 0.044 | KEIO_02_C8 | 0.794 | 0.046 |

| | | | | | | | | |
|---|---|---|---|---|---|---|---|---|
| b4062 | *soxS* | DNA-binding transcriptional dual regulator SoxS | KEIO_01_D8 | 0.726 | 0.158 | KEIO_02_D8 | 0.769 | -0.013 |
| b0726 | *sucA* | subunit of E1(0) component of 2-oxoglutarate dehydrogenase | KEIO_01_E12 | 0.732 | 0.015 | KEIO_02_E12 | 0.748 | 0.052 |
| b3839 | *tatC* | twin arginine protein translocation system - TatC protein | KEIO_61_B6 | 1.114 | 0.207 | KEIO_62_B6 | 0.981 | 0.006 |
| b3118 | *tdcA* | DNA-binding transcriptional activator TdcA | KEIO_01_F8 | 0.750 | 0.100 | KEIO_02_F8 | 0.804 | 0.051 |
| b2465 | *tktB* | transketolase 2 | KEIO_03_F5 | 0.977 | -0.019 | KEIO_04_F5 | 1.325 | 0.026 |
| b1914 | *uvrY* | DNA-binding transcriptional activator UvrY | KEIO_03_C11 | 0.777 | 0.069 | KEIO_04_C11 | 1.290 | 0.163 |
| b2059 | *wcaA* | putative colanic acid biosynthesis glycosyl transferase | KEIO_05_C7 | 0.855 | 0.025 | KEIO_06_C7 | 1.323 | -0.001 |
| b2405 | *xapR* | DNA-binding transcriptional activator XapR | KEIO_01_D9 | 0.972 | 0.079 | KEIO_02_D9 | 1.024 | -0.035 |
| b0208 | *yafC* | putative LysR family transcriptional regulator YafC | KEIO_02_F9 | 1.026 | -0.025 | KEIO_01_F9 | 0.966 | -0.022 |
| b0326 | *yahL* | uncharacterized protein YahL | KEIO_05_E9 | 0.965 | 0.096 | KEIO_06_E9 | 1.221 | 0.075 |
| b0603 | *ybdO* | putative LysR family DNA-binding transcriptional regulator | KEIO_01_G9 | 0.987 | -0.008 | KEIO_02_G9 | 1.062 | 0.050 |
| b1254 | *yciB* | inner membrane protein | KEIO_89_F10 | 0.983 | 0.075 | KEIO_90_F10 | 1.055 | 0.134 |
| b2109 | *yehB* | putative fimbrial usher protein YehB | KEIO_05_B7 | 0.747 | 0.027 | KEIO_06_B7 | 1.374 | 0.015 |
| b2337 | *yfcT* | | KEIO_05_B6 | 0.798 | -0.011 | KEIO_06_B6 | 1.241 | 0.002 |
| b2513 | *yfgM* | ancillary SecYEG translocon subunit | KEIO_05_B11 | 0.772 | 0.012 | KEIO_06_B11 | 1.144 | 0.010 |
| b3380 | *yhfW* | putative mutase YhfW | KEIO_03_B6 | 0.883 | 0.010 | KEIO_04_B6 | 1.198 | 0.084 |
| b4353 | *yjiX* | conserved protein YjiX | KEIO_39_C5 | 1.342 | 0.074 | KEIO_40_C5 | 0.947 | -0.043 |
| b0253 | *ykfA* | CP4-6 prophage%3B putative GTP-binding protein YkfA | KEIO_05_B9 | 0.977 | 0.097 | KEIO_06_B9 | 1.248 | 0.056 |
| b3292 | *zntR* | DNA-binding transcriptional activator ZntR | KEIO_01_E9 | 1.013 | -0.007 | KEIO_02_E9 | 0.996 | -0.028 |

**Table S5. Transposon library screening.** List of genes, whose disruption by transposon mutagenesis causes hypersensitivity to TAT-RasGAP$_{317\text{-}326}$ peptide in *P. aeruginosa* . UIC represents the unique insertion counts for each gene in absence (UIC_BM2) or in presence of the peptide (UIC_TATRasGAP). UID: Unique insertion density; NUID: normalized UID; FC: fold-change.

| Locus_tag | Locus_tag in PAO1 | Gene_ symbol | Description | Category | E. coli homologues | Hypersensitivity to PolB | UIC_BM2 | UIC_TATRasGAP | UID_BM2 | UID_TATRasGAP | NUID_BM2 | NUID_TATRasGAP | NUID_BM2vsTATRasGAP | $Log_2$(FC_BM2vsTATRasGAP) |
|---|---|---|---|---|---|---|---|---|---|---|---|---|---|---|
| PA14_31890 | PA2527 | | RND efflux transporter | Transmembrane transport | *mdtB* | | 172 | 1 | 0.055 | 0.000319 | 0.0063 | 4E-05 | 0.0062812 | -7.354948298 |
| PA14_31900 | PA2526 | | efflux transporter | Transmembrane transport | *mdtC* | | 387 | 3 | 0.124 | 0.000964 | 0.0143 | 0.0001 | 0.01419896 | -6.939910799 |
| PA14_13660 | PA3885 | *tpbA* | protein tyrosine phosphatase TpbA | Protein regulation | | | 86 | 1 | 0.131 | 0.001522 | 0.0151 | 0.0002 | 0.01487962 | -6.354948298 |
| PA14_63130 | PA4775 | | hypothetical protein | Unknown | | | 227 | 3 | 0.264 | 0.003484 | 0.0303 | 0.0004 | 0.02991901 | -6.17026953 |
| PA14_66140 | PA5002 | *dnpA* | de-N-acetylase involved in persistence | LPS biosynthesis | | | 60 | 1 | 0.042 | 0.000705 | 0.0049 | 9E-05 | 0.00478072 | -5.835574139 |
| PA14_57070 | PA4391 | | hypothetical protein | Unknown | | | 109 | 2 | 0.109 | 0.001996 | 0.0125 | 0.0002 | 0.01227727 | -5.696867868 |
| PA14_02390 | PA0191 | | transcriptional regulator | Transcription regulation | *cynR* | | 91 | 2 | 0.099 | 0.002179 | 0.0114 | 0.0003 | 0.01114422 | -5.436478184 |
| PA14_18350 | PA3554 | *arnA* | bifunctional UDP-glucuronic acid decarboxylase/UDP-4-amino-4-deoxy-L-arabinose formyltransferase | LPS biosynthesis | *arnA* | Yes | 312 | 7 | 0.157 | 0.003519 | 0.0181 | 0.0004 | 0.01762613 | -5.40673084 |
| PA14_31920 | PA2525 | *opmB* | outer membrane protein | Transmembrane transport | *cusC* | | 208 | 6 | 0.139 | 0.004008 | 0.016 | 0.0005 | 0.01550503 | -5.044160761 |
| PA14_31870 | PA2528 | | RND efflux membrane fusion protein | Transmembrane transport | *mdtA* | | 310 | 11 | 0.242 | 0.008587 | 0.0278 | 0.001 | 0.02681074 | -4.74537633 |
| PA14_58090 | PA4476 | | hypothetical protein | Unknown | | Yes | 317 | 14 | 0.083 | 0.003654 | 0.0095 | 0.0004 | 0.0090805 | -4.429667651 |
| PA14_71970 | PA5452 | *wbpW* | GDP-mannose pyrophosphorylase | LPS biosynthesis | *cpsB* | | 297 | 14 | 0.206 | 0.009722 | 0.0237 | 0.0012 | 0.02255958 | -4.335647742 |
| PA14_41260 | PA1799 | *parR* | two-component response regulator | Two-component regulator system | *rstA* | Yes | 243 | 12 | 0.343 | 0.016949 | 0.0395 | 0.002 | 0.03744822 | -4.268533546 |
| PA14_26770 | PA2884 | | hypothetical protein | membrane protease | | | 76 | 4 | 0.099 | 0.005229 | 0.0114 | 0.0006 | 0.0108005 | -4.176611057 |
| PA14_58990 | | | DNA helicase | Transcription regulation | | | 230 | 15 | 0.171 | 0.011136 | 0.0196 | 0.0013 | 0.01830332 | -3.867282999 |
| PA14_29710 | PA2659 | | hypothetical protein, putative peptidase | Protein regulation | | | 30 | 2 | 0.097 | 0.006472 | 0.0112 | 0.0008 | 0.01039015 | -3.835574139 |
| PA14_63160 | PA4777 | *pmrB* | two-component regulator system signal sensor kinase | Two-component regulator system | *qseC* | Yes | 201 | 17 | 0.14 | 0.011855 | 0.0161 | 0.0014 | 0.01469697 | -3.492272393 |
| PA14_27190 | | *RNA_23* | tRNA-Ser | Translation regulation | | Yes | 23 | 2 | 0.264 | 0.022989 | 0.0304 | 0.0028 | 0.0276437 | -3.452245499 |
| PA14_13650 | PA3886 | | hypothetical protein | Unknown | | | 189 | 18 | 0.21 | 0.02 | 0.0242 | 0.0024 | 0.02174843 | -3.321000966 |
| PA14_23890 | PA3110 | | hypothetical protein | Peptidoglycan binding | | Yes | 85 | 11 | 0.131 | 0.016975 | 0.0151 | 0.0021 | 0.01304278 | -2.878642861 |
| PA14_59590 | | | hypothetical protein | Unknown | | | 23 | 3 | 0.064 | 0.008333 | 0.0074 | 0.001 | 0.0063447 | -2.867282999 |
| PA14_49410 | PA1159 | | cold-shock protein | Transcription regulation | *cspA* | | 15 | 2 | 0.071 | 0.009524 | 0.0082 | 0.0012 | 0.00706842 | -2.835574139 |
| PA14_42920 | PA1667 | *hsiJ2* | HsiJ2 | Protein secretion | | Yes | 251 | 40 | 0.188 | 0.03003 | 0.0217 | 0.0036 | 0.0180544 | -2.578299002 |
| PA14_17140 | PA3649 | | membrane-associated zinc metalloprotease | Quorum sensing | *rseP* | | 91 | 15 | 0.067 | 0.011086 | 0.0077 | 0.0013 | 0.00639952 | -2.529587588 |

| | | | | | | | | | | | | | | |
|---|---|---|---|---|---|---|---|---|---|---|---|---|---|---|
| PA14_33880 | PA2380 | | hypothetical protein | Unknown | | | 12 | 2 | 0.051 | 0.008439 | 0.0058 | 0.001 | 0.00480645 | -2.513646044 |
| PA14_51320 | PA1005 | | hypothetical protein, putative peptidase | Protein regulation | | | 149 | 26 | 0.104 | 0.018131 | 0.012 | 0.0022 | 0.00976508 | -2.447412346 |
| PA14_18480 | PA3546 | *algX* | alginate biosynthesis protein AlgX | Alginate biosynthesis | | | 141 | 25 | 0.099 | 0.017544 | 0.0114 | 0.0021 | 0.00926555 | -2.424378706 |
| PA14_18520 | PA3543 | *algK* | alginate biosynthetic protein AlgK | Alginate biosynthesis | | | 71 | 14 | 0.05 | 0.009804 | 0.0057 | 0.0012 | 0.00453633 | -2.271075741 |
| PA14_59060 | | | hypothetical protein | Unknown | | | 15 | 3 | 0.059 | 0.011765 | 0.0068 | 0.0014 | 0.00534689 | -2.250611638 |
| PA14_65300 | PA4943 | *hflX* | GTP-binding protein | Translation regulation | *hflX* | Yes | 68 | 15 | 0.052 | 0.011521 | 0.006 | 0.0014 | 0.0046173 | -2.109255789 |
| PA14_22490 | PA3223 | *acpD* | ACP phosphodiesterase | Lipid metabolism | *azoR* | | 77 | 17 | 0.12 | 0.02648 | 0.0138 | 0.0032 | 0.01060065 | -2.108007243 |
| PA14_02380 | PA0190 | | acid phosphatase | Phosphatase activity | | | 73 | 19 | 0.101 | 0.026171 | 0.0116 | 0.0032 | 0.008407 | -1.870580589 |
| PA14_23240 | PA3170 | | N-ethylammeline chlorohydrolase | Amino acid metabolism | *guaD* | | 351 | 96 | 0.263 | 0.07191 | 0.0303 | 0.0087 | 0.02156209 | -1.799048263 |
| PA14_10710 | PA4116 | *bphO* | heme oxygenase | Oxido-reduction | | | 149 | 41 | 0.253 | 0.069728 | 0.0292 | 0.0084 | 0.02073039 | -1.790300059 |
| PA14_67100 | PA5079 | | D-tyrosyl-tRNA(Tyr) deacylase | Translation regulation | *dtd* | | 90 | 26 | 0.205 | 0.059361 | 0.0236 | 0.0072 | 0.01646907 | -1.720096922 |
| PA14_70450 | PA5337 | *rpoZ* | DNA-directed RNA polymerase subunit omega | Transcription regulation | *rpoZ* | | 153 | 45 | 0.573 | 0.168539 | 0.0659 | 0.0204 | 0.04556607 | -1.69421829 |
| PA14_55330 | PA0695 | | hypothetical protein | Unknown | | | 203 | 62 | 0.272 | 0.082999 | 0.0313 | 0.01 | 0.02123778 | -1.63982315 |
| PA14_65200 | PA4937 | *rnr* | exoribonuclease RNase R | Translation regulation | *rnr* | | 327 | 101 | 0.12 | 0.037078 | 0.0138 | 0.0045 | 0.00933147 | -1.623618886 |
| PA14_69250 | PA5244 | | hypothetical protein | Unknown | | Yes | 116 | 36 | 0.196 | 0.060914 | 0.0226 | 0.0074 | 0.01522234 | -1.616739537 |
| PA14_16370 | PA3712 | | hypothetical protein | Unknown | | | 193 | 61 | 0.277 | 0.087644 | 0.0319 | 0.0106 | 0.02131428 | -1.590403243 |
| PA14_08680 | PA4277 | *tufB* | elongation factor Tu | Translation regulation | *tufB* | | 102 | 34 | 0.085 | 0.028476 | 0.0098 | 0.0034 | 0.00638787 | -1.513646044 |
| PA14_43900 | PA1592 | | hypothetical protein | Unknown | | | 27 | 9 | 0.114 | 0.037975 | 0.0131 | 0.0046 | 0.00851875 | -1.513646044 |
| PA14_60250 | PA4546 | *pilS* | two-component sensor PilS | Two-component regulator system | *atoS* | | 195 | 66 | 0.122 | 0.041431 | 0.0141 | 0.005 | 0.00907743 | -1.491619738 |
| PA14_08830 | PA4265 | *tufA* | elongation factor Tu | Translation regulation | *tufA* | | 103 | 35 | 0.086 | 0.029313 | 0.0099 | 0.0035 | 0.00638299 | -1.485901054 |
| PA14_49020 | | | hypothetical protein | Unknown | | Yes | 117 | 40 | 0.438 | 0.149813 | 0.0504 | 0.0181 | 0.03231401 | -1.477120168 |
| PA14_51620 | PA0979 | | transposase | Others | | | 139 | 48 | 0.45 | 0.15534 | 0.0518 | 0.0188 | 0.03298478 | -1.462662115 |
| PA14_63040 | PA4766 | | hypothetical protein | Unknown | | Yes | 23 | 8 | 0.075 | 0.026144 | 0.0086 | 0.0032 | 0.00548868 | -1.452245499 |
| PA14_42700 | PA1686 | *alkA* | DNA-3-methyladenine glycosidase II | DNA repair | *alkA* | | 112 | 44 | 0.125 | 0.049217 | 0.0144 | 0.006 | 0.00846622 | -1.276606847 |
| PA14_04160 | PA0319 | | transcriptional regulator | Transcription regulation | *yjiK* | | 185 | 75 | 0.187 | 0.075758 | 0.0215 | 0.0092 | 0.0123448 | -1.231246313 |
| PA14_11670 | PA4033 | *mucE* | hypothetical protein | Unknown | | | 64 | 26 | 0.237 | 0.096296 | 0.0273 | 0.0116 | 0.01563478 | -1.228243825 |
| PA14_59990 | | | hypothetical protein | Unknown | | | 549 | 225 | 0.585 | 0.239617 | 0.0673 | 0.029 | 0.03831052 | -1.215564691 |
| PA14_12560 | PA3966 | | hypothetical protein | Unknown | | | 17 | 7 | 0.081 | 0.033333 | 0.0093 | 0.004 | 0.00528558 | -1.208791463 |
| PA14_48040 | PA1250 | *aprI* | alkaline proteinase inhibitor AprI | Protein regulation | *sapA* | | 29 | 12 | 0.073 | 0.030303 | 0.0084 | 0.0037 | 0.00476356 | -1.201702038 |
| PA14_14540 | | | hypothetical protein | Unknown | | | 33 | 14 | 0.175 | 0.074074 | 0.0201 | 0.009 | 0.01113685 | -1.165722741 |
| PA14_15410 | | | hypothetical protein | Unknown | | | 42 | 18 | 0.209 | 0.089552 | 0.024 | 0.0108 | 0.01321858 | -1.151075965 |
| PA14_28600 | PA2747 | | hypothetical protein | Unknown | | | 35 | 15 | 0.122 | 0.052083 | 0.014 | 0.0063 | 0.00768789 | -1.151075965 |

| | | | | | | | | | | | | | | |
|---|---|---|---|---|---|---|---|---|---|---|---|---|---|---|
| PA14_59140 | | | hypothetical protein | Unknown | | | 107 | 46 | 0.2 | 0.086142 | 0.0231 | 0.0104 | 0.01264341 | -1.146588574 |
| PA14_18430 | PA3549 | *algJ* | alginate o-acetyltransferase AlgJ | Alginate biosynthesis | | | 421 | 181 | 0.358 | 0.153912 | 0.0412 | 0.0186 | 0.02258801 | -1.146514079 |
| PA14_21530 | PA3287 | | ankyrin domain-containing protein | Unknown | | | 58 | 25 | 0.112 | 0.04845 | 0.0129 | 0.0059 | 0.00707718 | -1.142808349 |
| PA14_26485 | PA2906 | | oxidoreductase | Oxido-reduction | | | 117 | 51 | 0.08 | 0.034908 | 0.0092 | 0.0042 | 0.00499509 | -1.126622921 |
| PA14_18690 | PA3529 | | peroxidase | Oxido-reduction | *ahpC* | | 48 | 21 | 0.08 | 0.034826 | 0.0092 | 0.0042 | 0.00494971 | -1.121328621 |
| PA14_11320 | PA4063 | | hypothetical protein | Unknown | | | 54 | 24 | 0.091 | 0.040609 | 0.0105 | 0.0049 | 0.00560477 | -1.098608545 |
| PA14_41280 | PA1797 | | putative beta-lactamase | Unknown | | | 158 | 71 | 0.086 | 0.038734 | 0.0099 | 0.0047 | 0.00523614 | -1.082717172 |
| PA14_51830 | PA0962 | | DNA-binding stress protein | Transcription regulation | | | 62 | 28 | 0.132 | 0.059448 | 0.0151 | 0.0072 | 0.00796053 | -1.075524932 |
| PA14_13630 | | | hypothetical protein | Unknown | | | 46 | 21 | 0.187 | 0.085366 | 0.0215 | 0.0103 | 0.01119723 | -1.059928077 |
| PA14_22340 | PA3235 | | hypothetical protein | Unknown | | | 35 | 16 | 0.112 | 0.051282 | 0.0129 | 0.0062 | 0.00670898 | -1.05796656 |
| PA14_63250 | PA4785 | | acetyl-CoA acetyltransferase | Carbon metabolism | | | 192 | 88 | 0.15 | 0.068858 | 0.0173 | 0.0083 | 0.00896327 | -1.054214425 |
| PA14_21680 | PA3273 | | hypothetical protein | Unknown | | | 98 | 45 | 0.163 | 0.075 | 0.0188 | 0.0091 | 0.00972797 | -1.051540291 |
| PA14_52890 | PA0880 | | ring-cleaving dioxygenase | Antibiotic resistance | *gloA* | | 153 | 71 | 0.402 | 0.186352 | 0.0462 | 0.0225 | 0.02368099 | -1.036324267 |
| PA14_52800 | PA0887 | *acsA* | acetyl-CoA synthetase | Carbon metabolism | *acs* | | 299 | 140 | 0.153 | 0.071575 | 0.0176 | 0.0087 | 0.00893721 | -1.023402201 |
| PA14_41270 | PA1798 | *parS* | two-component sensor | Two-component regulator system | *rstB* | Yes | 74 | 0 | 0.057 | 0 | 0.0066 | 0 | 0.00661684 | #N/A |
| PA14_18330 | PA3556 | *arnT* | 4-amino-4-deoxy-L-arabinose transferase | LPS biosynthesis | *arnT* | Yes | 105 | 0 | 0.064 | 0 | 0.0073 | 0 | 0.00732323 | #N/A |
| PA14_56950 | PA4381 | *colR* | two-component response regulator | Two-component regulator system | *cusR* | Yes | 49 | 0 | 0.072 | 0 | 0.0082 | 0 | 0.00824399 | #N/A |
| PA14_18370 | PA3552 | *arnB* | UDP-4-amino-4-deoxy-L-arabinose--oxoglutarate aminotransferase | LPS biosynthesis | *arnB* | Yes | 97 | 0 | 0.084 | 0 | 0.0097 | 0 | 0.00971514 | #N/A |
| PA14_49180 | PA1179 | *phoP* | two-component response regulator PhoP | Two-component regulator systen | *phoP* | Yes | 103 | 0 | 0.152 | 0 | 0.0175 | 0 | 0.01748255 | #N/A |

**Table S6.** MICs of passages of *E.coli* selected for resistance to TAT-RasGAP$_{317-326}$ peptide. Bacteria were treated as described in Figure 8. MIC was measured for each passage, in absence of selection (MIC non selected) or with selection (MIC selected). Concentrations of TAT-RasGAP$_{317-326}$ each passage was exposed to are indicated.

| Passages | MIC non selected | MIC selected | μM TAT-RasGAP$_{317-326}$ |
|---|---|---|---|
| 1 | 4 | 8 | 2 |
| 2 | 4 | 4 | 2 |
| 3 | 4 | 8 | 2 |
| 4 | 4 | 8 | 2 |
| 5 | 4 | 16 | 5 |
| 6 | 4 | 16 | 10 |
| 7 | 4 | 16 | 10 |
| 8 | 4 | 16 | 10 |
| 9 | 4 | 16 | 10 |
| 10 | 4 | 16 | 10 |
| 11 | 4 | 32 | 20 |
| 12 | 4 | 32 | 20 |
| 13 | 4 | 32 | 30 |
| 14 | 4 | 32 | 30 |
| 15 | 4 | 32 | 30 |
| 16 | 4 | 64 | 40 |
| 17 | 4 | 64 | 40 |
| 18 | 4 | 64 | 60 |
| 19 | 4 | 64 | 60 |
| 20 | 4 | 64 | 80 |

**Table S7:** MICs of passages of *P.aeruginosa* selected for resistance to TAT-RasGAP$_{317-326}$ peptide. Bacteria were treated as described in Figure 8. MIC was measured for each passage, in absence of selection (MIC non selected) or with selection (MIC selected). Concentrations of TAT-RasGAP$_{317-326}$ each passage was exposed to are indicated.

| Passages | MIC non selected | MIC selected | μM TAT-RasGAP$_{317-326}$ |
|---|---|---|---|
| 1 | 2 | 2 | 2 |
| 2 | 2 | 2 | 4 |
| 3 | 2 | 2 | 4 |
| 4 | 2 | 2 | 4 |
| 5 | 2 | 2 | 4 |
| 6 | 2 | 2 | 8 |
| 7 | 2 | 2 | 8 |
| 8 | 2 | 4 | 8 |
| 9 | 2 | 4 | 4 |
| 10 | 2 | 4 | 8 |
| 11 | 2 | 4 | 12 |
| 12 | 2 | 4 | 12 |
| 13 | 2 | 4 | 16 |
| 14 | 4 | 4 | 24 |
| 15 | 2 | 8 | 24 |
| 16 | 2 | 4 | 20 |
| 17 | 2 | 8 | 20 |
| 18 | 2 | 4 | 20 |
| 19 | 4 | 4 | 20 |
| 20 | 4 | 8 | 20 |

**Table S8:** MICs of passages of *S. capitis* selected for resistance to TAT-RasGAP$_{317-326}$ peptide. Bacteria were treated as described in Figure 8. MIC was measured for each passage, in absence of selection (MIC non selected) or with selection (MIC selected). Concentrations of TAT-RasGAP$_{317-326}$ each passage was exposed to are indicated.

| Passages | MIC non selected | MIC selected | μM TAT-RasGAP$_{317-326}$ |
|---|---|---|---|
| 1 | 2 | 2 | 0.5 |
| 2 | 2 | 2 | 1 |
| 3 | 2 | 2 | 1 |
| 4 | 2 | 2 | 2 |
| 5 | 2 | 8 | 4 |
| 6 | 2 | 8 | 4 |
| 7 | 2 | 8 | 4 |
| 8 | 2 | 16 | 8 |
| 9 | 2 | 8 | 4 |
| 10 | 2 | 16 | 8 |
| 11 | 2 | 16 | 12 |
| 12 | 2 | 16 | 16 |
| 13 | 2 | 32 | 20 |
| 14 | 2 | 32 | 20 |
| 15 | 2 | 32 | 24 |
| 16 | 2 | 64 | 24 |
| 17 | 2 | 64 | 24 |
| 18 | 2 | 64 | 24 |
| 19 | 2 | 64 | 24 |
| 20 | 2 | 64 | 24 |

**Table S9:** MICs of passages of *S. aureus* selected for resistance to TAT-RasGAP$_{317\text{-}326}$ peptide. Bacteria were treated as described in Figure 8. MIC was measured for each passage, in absence of selection (MIC non selected) or with selection (MIC selected). Concentrations of TAT-RasGAP$_{317\text{-}326}$ each passage was exposed to are indicated.

| Passages | MIC non selected | MIC selected | μM TAT-RasGAP$_{317\text{-}326}$ |
|---|---|---|---|
| 1 | 128 | 128 | 2 |
| 2 | 64 | 64 | 5 |
| 3 | 64 | 64 | 10 |
| 4 | 64 | >256 | 20 |
| 5 | 128 | >256 | 40 |
| 6 | 128 | >256 | 60 |
| 7 | 64 | >256 | 80 |
| 8 | 128 | >256 | 120 |
| 9 | 64 | >256 | 180 |
| 10 | 128 | >256 | 240 |
| 11 | 128 | >256 | 300 |
| 12 | 128 | >256 | 450 |
